# Oncogene-induced matrix reorganization controls CD8^+^ T cell function in the soft-tissue sarcoma microenvironment

**DOI:** 10.1101/2022.03.31.486627

**Authors:** Ashley M. Fuller, Hawley C. Pruitt, Ying Liu, Valerie Irizarry-Negron, Hehai Pan, Hoogeun Song, Ann DeVine, Rohan Katti, Samir Devalaraja, Gabrielle E. Ciotti, Michael Gonzalez, Erik F. Williams, Ileana Murazzi, Dimitris Ntekoumes, Nicolas Skuli, Hakon Hakonarson, Daniel Zabransky, Jose Trevino, Ashani Weeraratna, Kristy Weber, Malay Haldar, Joseph A. Fraietta, Sharon Gerecht, T. S. Karin Eisinger-Mathason

## Abstract

CD8^+^ T cell dysfunction impedes anti-tumor immunity in solid cancers but the underlying mechanisms are diverse and poorly understood. Extracellular matrix (ECM) composition has been linked to both impaired T cell migration and enhanced tumor progression; however, impacts of individual ECM molecules on T cell function in the tumor microenvironment (TME) are only beginning to be elucidated. Upstream regulators of aberrant ECM deposition and organization in solid tumors are equally ill-defined. Therefore, we investigated how ECM composition modulates CD8^+^ T cell function in undifferentiated pleomorphic sarcoma (UPS), an immunologically active and desmoplastic tumor. Using an autochthonous murine model of UPS and data from multiple human patient cohorts, we discovered a multifaceted mechanism wherein the transcriptional co-activator YAP1 promotes collagen VI (COLVI) deposition in the UPS TME. In turn, COLVI induces CD8^+^ T cell dysfunction and immune evasion by remodeling fibrillar collagen and inhibiting T cell autophagic flux. Unexpectedly, collagen I (COLI) opposed COLVI in this setting, promoting CD8^+^ T cell function and acting as a tumor suppressor. Thus, CD8^+^ T cell responses in sarcoma depend upon oncogene-mediated ECM composition and remodeling.

## INTRODUCTION

Immunosuppression in the solid tumor microenvironment (TME) is a barrier to T cell-mediated anti-tumor immunity. Tumors evade host adaptive immune responses by inducing CD8^+^ T cell dysfunction, a hypofunctional state characterized by overexpression of inhibitory cell-surface receptors (e.g., PD1, TIM-3, LAG3), reduced effector function, and impaired proliferative capacity (1). Molecular mechanisms underlying CD8^+^ T cell dysfunction in solid cancers are of significant interest due to their impact on immunotherapy strategies. However, most prior studies in this area have focused on the roles of continuous antigen exposure/repetitive T cell receptor stimulation, immune checkpoint-mediated inhibitory signaling, and immunosuppressive cytokines (2). Moreover, the importance of TME contexture in the setting of T cell-based therapies is poorly described. Thus, a more comprehensive and physiological evaluation of CD8^+^ T cell dysfunction in solid tumors is critical for improving our understanding of immune evasion mechanisms in the TME and advancing actionable interventions.

Soft tissue sarcoma (STS) is a heterogeneous group of solid mesenchymal tumors comprised of ∼70 distinct histologic subtypes (3). These lesions are characterized by mesenchymal gene expression, extensive extracellular matrix (ECM) deposition, and increased ECM stiffness relative to normal tissues (4–7). Interestingly, these features are also observed in high-grade, poorly differentiated epithelial tumors where they are linked to progression, therapeutic resistance, and poor clinical outcomes (8–16). Recent studies have shown that the ECM facilitates cancer progression in part by inhibiting T cell migration/infiltration (17–20). However, the roles of individual ECM proteins in this process are only beginning to be defined. Moreover, little research has addressed the effects of ECM molecules on T cell function, particularly in solid tumors, or identified upstream regulators of aberrant ECM deposition and composition/organization in this context. This paucity of available data indicates that further study, particularly in vivo, is necessary.

Members of the collagen superfamily, of which there are 28 distinct molecular species, are some of the most abundant and diverse ECM constituents in both normal tissues and solid tumors (21). Although the roles of specific collagen species in cancer-associated processes are ill-defined, a growing body of literature indicates that individual collagen molecules can have context-specific functions in the TME. For example, type I collagen (ColI), a fibrillar collagen that forms prototypical collagen fibers, promotes or is associated with malignant progression in some tumor settings, but has anti-tumor effects in others (22–28). These findings underscore the need to systematically interrogate the roles of individual collagen molecules, particularly with respect to their potential impacts on adaptive immunity, in specific tumor contexts.

Undifferentiated pleomorphic sarcoma (UPS) is a relatively common STS subtype that predominantly arises in adult skeletal muscle and has a 10-year survival rate of only ∼25% (3, 29). Although some STS are considered immunologically “cold”, recent clinical trials have revealed that patients with UPS can exhibit objective clinical responses to immune checkpoint inhibition (30, 31). These encouraging findings suggest that studies of UPS may provide valuable insights into strategies for enhancing T cell function and immunotherapy responses in solid tumors. Our previous work linked the intrinsic oncogenic functions of the transcriptional co-regulator Yes-associated protein 1 (YAP1), the central Hippo pathway effector, to UPS cell proliferation, tumor growth, and reduced human patient survival (32–35). However, we had not investigated the contribution of YAP1 to the broader UPS microenvironment or immune cell activity. In some epithelial tumors, cancer cell-intrinsic YAP1 modulates recruitment and differentiation of macrophages and myeloid-derived suppressor cells, suggesting a role in immunomodulation (36, 37). However, this observation has not been confirmed in mesenchymal cancers. YAP1 also possesses mechanosensory functions, and its nuclear localization and activity increase in response to stiff environments such as those found in tumor tissue (38). Therefore, in this study, we interrogated the role of UPS cell-intrinsic YAP1 signaling in the regulation of ECM deposition/organization and adaptive immune cell function in the TME. We discovered a novel role for YAP1 in the regulation of ECM composition and cytotoxic T cell function, and found that collagen type VI (ColVI), a microfibrillar collagen, indirectly modulates effector T cell function by opposing and remodeling collagen type I (ColI). We further identify COLVI as a putative ECM-associated biomarker of diagnosis and survival in human UPS. Our findings implicate YAP1 inhibition, in combination with immunotherapy, as a promising approach to mitigate immune evasion mechanisms in the TME of patients with solid tumors.

## RESULTS

### UPS cell-intrinsic Yap1 inhibits T cell activation and promotes CD8^+^ T cell dysfunction

Using the genetically engineered mouse model (GEMM) of skeletal muscle-derived UPS, *Kras*^G12D/+^; *Trp53*^fl/fl^ (KP) (39, 40), we previously showed that Yap1 promotes UPS tumorigenesis and progression via activation of sarcoma cell-intrinsic NF-κB (33). In this GEMM system, tumors are generated by injecting adenovirus-expressing Cre recombinase into the gastrocnemius muscle. Recombination initiates oncogenic *Kras* expression and deletes *Yap1* floxed alleles in infected muscle progenitor cells (39, 40). *TP53* mutations and deletion are prevalent in human UPS (41), as is hyperactivation of the MAPK pathway downstream of KRAS (42). Consistent with our previous work (33), we observed significantly increased tumor latency and similar rates of tumor development when we introduced *Yap1*^fl/fl^ alleles into the KP GEMM, creating *LSL*-*Kras*^G12D/+^; *Trp53*^fl/fl^; *Yap1*^fl/fl^ (KPY) animals (**Figure 1A, B**, Supp. Figure 1A).

**Figure 1.**
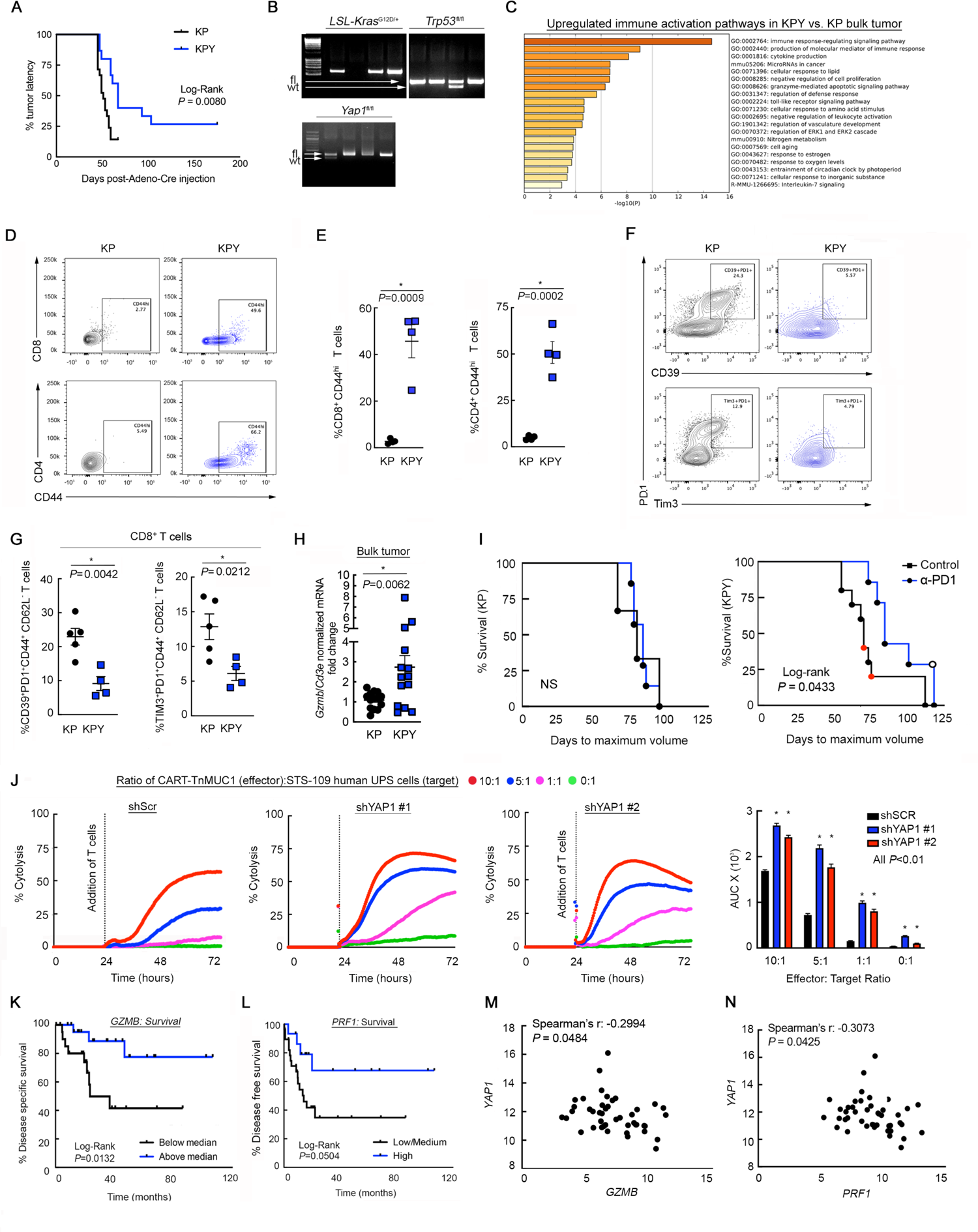
YAP1^+^ UPS cells inhibit CD8^+^ T cell activation and promote dysfunction. (**A**) Kaplan-Meier latency curves of KP and KPY UPS tumors (n >10 per genotype). Log-rank test. (**B**) Validation of genotypes from **A**. (**C**) Metascape pathway analysis of 5 unique bulk KP and KPY tumors. Includes all genes with >2-fold expression increase in KPY vs. KP, identified via microarrays. (**D, E**) Representative contour plots (**D**) and quantification (**E**) of CD8^+^CD44^hi^ and CD4^+^CD44^hi^ T cells in KP and KPY tumors. Each point represents an individual tumor; two-tailed unpaired t-tests. (**F, G**) Representative contour plots (**F**) and quantification (**G**) of CD39, Tim-3, and Pd1 expression in CD8^+^ T cells from KP and KPY tumors. Each point represents an individual tumor. Two-tailed unpaired t-tests. (**H**) qRT-PCR of *Gzmb* in bulk KP and KPY tumors; two-tailed unpaired t-test. (**I**) Kaplan-Meier survival curves of KP and KPY tumor-bearing mice treated with α-Pd1 or control. Red and black circles in control curves indicate IgG- and un-injected mice, respectively. Open circle: mouse with durable tumor regression. X-axis: days since adeno Cre injection. Log-rank test. (**J**) Longitudinal cytolysis of shScr or shYAP1 human STS-109 UPS cells during co-culture with CART-TnMUC1 cells. Measurements indicate % target (UPS) cell cytolysis. Normalized cell index for STS-109 cells alone (not shown) controls for differences in target cell growth across conditions. Quantification: area under the curve (AUC); one-way ANOVA with Dunnett’s (vs. shScr) for each ratio. shScr data are identical to those in Fig. 3D**-E** (performed in the same experiment). (**K**, **L**) Kaplan-Meier survival curves of UPS patients (n = 44) in TCGA-Sarcoma stratified by intratumoral *GZMB* (**K**) and *PRF1* (**L**) expression. Each tertile (low, medium, high) represents 1/3 of patients. Log-rank test. (**M**, **N**) Correlation of *YAP1* with *GZMB* (**M**) and *PRF1* (**N**) gene expression in UPS tumors from TCGA-Sarcoma.

Our previous studies have focused on mechanisms by which Yap1 impacts sarcoma cell-autonomous signaling and phenotypes such as proliferation, differentiation, and metastasis (32–34). Therefore, in the present study, we tested the hypothesis that UPS-cell intrinsic Yap1 also impacts the TME. To identify potential mechanisms of Yap1-mediated TME modulation, we explored a publicly available microarray-based gene expression dataset previously published by our group (33) comparing 5 unique KP and KPY bulk tumors. Loss of *Yap1* enhanced expression of numerous pathways associated with immune activation, suggesting that UPS cell-intrinsic Yap1 contributes to immunosuppression through an unknown mechanism (**Figure 1C**). To investigate how Yap1 controls immunosuppression in UPS, we performed flow cytometric and automated immunohistochemical (IHC) analyses of KP and KPY tumors. We did not detect changes in myeloid cell (dendritic cells, macrophages, neutrophils) infiltration or polarization, nor differences in B cell content (Supp. Figure 1B**-D**). However, we did observe increased proportions of CD44^hi^CD8^+^ and CD44^hi^CD4^+^ T cells in KPY relative to KP tumors, indicating enhanced T cell activation (**Figure 1D, E**). Furthermore, the percentage of dysfunctional effector CD8^+^ T cells (CD39^+^/Pd1^+^ and Tim-3^+^/Pd1^+^ CD8^+^ T cells) was higher in KP tumors compared to KPY (**Figure 1F, G**). Markers of central CD8^+^ memory T cell differentiation (CD62L, CD127) remained unchanged (Supp. Figure 2A). Importantly, in KPY mice, the observed increases in T cell activation could not be attributed to reduced immunosuppressive Foxp3^+^ T regulatory cell content, nor to enhanced effector T cell infiltration (Supp. Figure 1B-D). In fact, CD4^+^ and CD8^+^ T cell content was modestly decreased in KPY relative to KP tumors. Therefore, we conclude that Yap1 may promote CD8^+^ T cell dysfunction but likely does not impact T cell recruitment to the TME.

To further explore the relationship between Yap1^+^ UPS cells and T cell activation, we evaluated expression of the T cell cytolysis marker, granzyme B (*Gzmb*) (43), in GEMM tumors (**Figure 1H**). *Gzmb* mRNA expression (normalized to total T cells; *Cd3e*) was significantly increased in KPY tumors, providing further evidence that Yap1^+^ UPS cells are associated with immunosuppression. Therefore, we determined if UPS cell-intrinsic Yap1 modulates T cell effector function in addition to inhibitory surface marker expression. To this end, we treated tumor-bearing KP and KPY mice with α-Pd1 or isotype control antibody. We hypothesized that immune checkpoint blockade would show increased efficacy in KPY animals due to enhanced T cell activation, but have no effect in KP mice. Consistent with this hypothesis, time to maximum tumor volume was significantly increased in KPY, but not KP, animals (**Figure 1I**). Notably, one KPY mouse experienced complete and lasting tumor regression. We further evaluated the effect of YAP1^+^ UPS cells on T cell function by leveraging human chimeric antigen receptor T (CART) cells that target the Tn glycoform of mucin 1 on human cancer cells (TnMUC1 CART cells (44)). This antigen is expressed on human STS-109 cells, derived from a UPS patient tumor (Supp. Figure 2B). We co-cultured TnMUC1-CART cells with STS-109 cells expressing control or *YAP1-* specific shRNAs (shYAP1) at multiple effector:target ratios and analyzed longitudinal cytolysis (**Figure 1J, Supp.** Figure 2C). We observed that *YAP1* deficiency in UPS cells enhances cytotoxic T cell function, confirming that YAP1*^+^* UPS cells promote immunosuppression. To explore this data in a human patient context, we leveraged The Cancer Genome Atlas (TGCA) Sarcoma dataset (41). Consistent with our experimental findings, gene expression levels of T cell cytolysis markers (*GZMB* and perforin [*PRF1*]) in human UPS tumors were associated with improved survival (**Figure 1K, L**, Supp. Figure 2D) and negatively correlated with *YAP1* levels (**Figure 1M, N**). Thus, although some sarcomas are considered immunologically “cold”, our data suggest that cytotoxic T cell activation is a critical factor in UPS patient survival, and that modulating Yap1 and T cell activity may improve clinical outcomes.

### UPS cell-intrinsic Yap1 promotes collagen VI deposition in the TME

We next sought to define the mechanism of crosstalk between Yap1^+^ UPS cells and infiltrating CD8^+^ T cells. Recent studies in epithelial tumors have shown that Yap1 can influence the cancer cell “secretome” (45, 46). Therefore, we measured 31 cytokines and chemokines in supernatants from the same samples used in our CAR T cell cytolysis assays (**Figure 1J**). Many analytes were below the lower limit of detection of the assay, but those we could detect were generally stable in UPS cell mono-cultures vs. co-cultures, and in cultures with *YAP1*-sufficient vs. *YAP1*-deficient UPS cells (Supp. Figure 3A). UPS cells co-cultured with normal human donor T cells (ND T cells) yielded similar results (Supp. Figure 3B). These findings support the conclusion that YAP1 likely does not control CD8^+^ T cell function in the TME via cytokines or chemokines; thus, we focused on other potential mechanisms.

YAP1 is a known modulator of mechanosensing properties associated with ECM remodeling (47). Therefore, we investigated whether ECM-related processes are required for YAP1-mediated T cell suppression in the UPS TME. Using our microarray dataset of KP and KPY tumors to identify Yap1-dependent matrix genes, we found that many pathways associated with ECM and tissue remodeling were altered in KPY tumors relative to KP (Supp. Figure 3C). We also observed that genes encoding many members of the collagen superfamily, particularly collagen type VI (ColVI), were downregulated in KPY relative to KP tumors (**Figure 2A, Supp.** Figure 3D). Multiple genes encode ColVI (e.g., *Col6a1*, *Col6a2*, *Col6a3*), each of which results in a unique protein chain. *Col6a1* is indispensable for ColVI protein expression (48). qRT-PCR and IHC analysis of bulk tumor specimens revealed that KPY tumors exhibited a trend toward reduced ColVI deposition overall (**Figure 2B, C**, Supp. Figure 3E-H). We did observe some heterogeneity in expression, potentially due to ColVI secretion by multiple cell types including macrophages (28, 49) and UPS cells themselves; however, IHC analysis clearly showed that KPY tumors exhibited significantly less strong-positive (3+) and significantly more moderately positive (2+) staining than KP tumors (**Figure 2B, C**). We validated these findings in vitro by qRT-PCR and immunoblotting of UPS cell lines derived from multiple unique KP GEMM tumors (KP230 and SKPY42.1 cells, referred to hereafter as “KP cells”). Specifically, KP cells transduced with one of multiple *Yap1*-specific shRNAs expressed substantially less *Col6a1* and *Col6a2* than control cells; *Col6a3* was more modestly reduced (**Figure 2D, E**). In contrast, we could not validate a role for Yap1 in the modulation of collagen type III expression (*Col3a1*; Supp. Figure 3I), nor that of other matrix genes such as fibronectin (*Fn1*; Supp. Figure 3J), indicating the potential specificity of this regulation.

**Figure 2.**
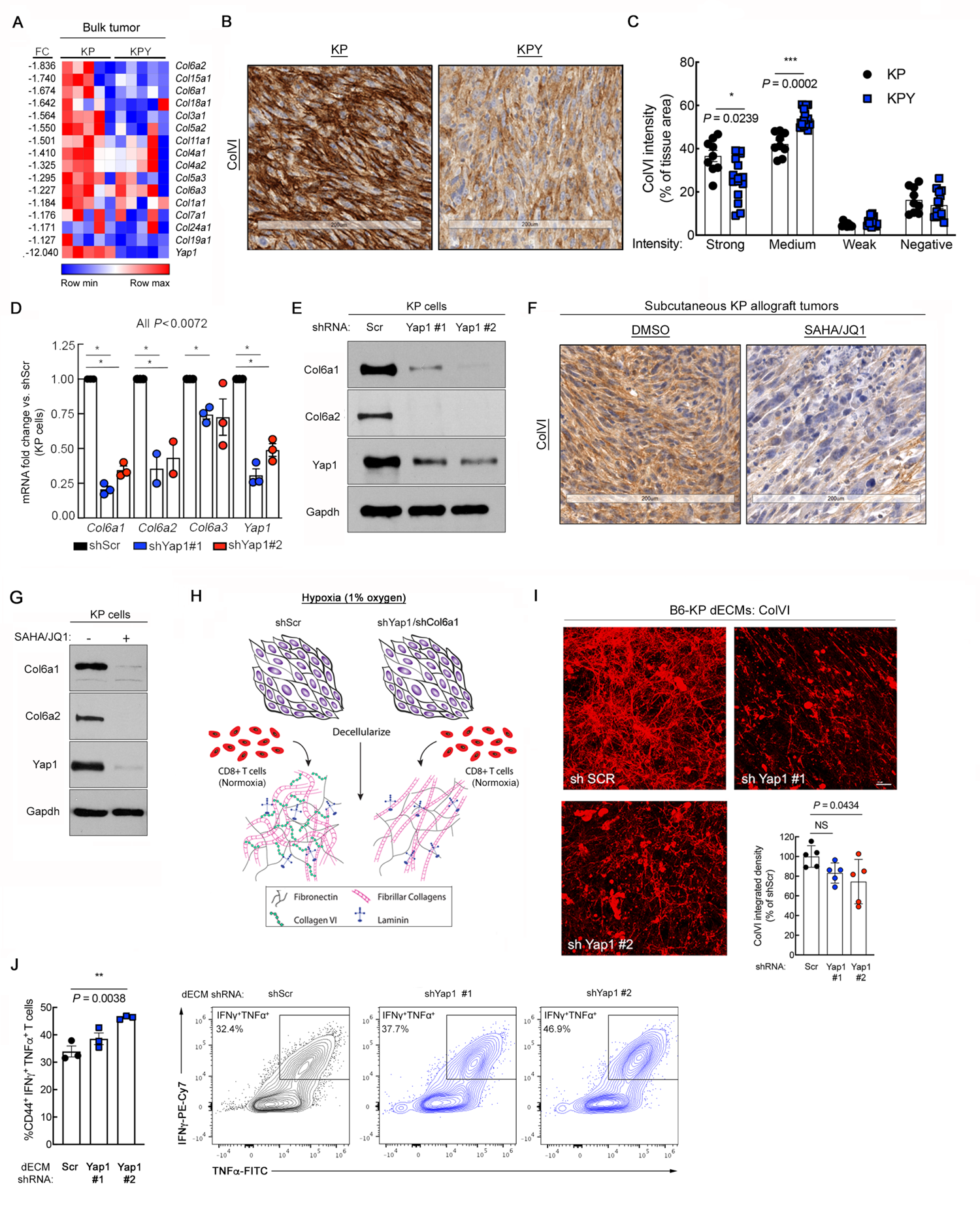
UPS-cell intrinsic Yap1 mediates collagen VI (ColVI) deposition in the TME. (**A**) Heat map of gene expression microarray data comparing 5 unique KP and KPY bulk tumors. The top 1/3 of collagen-encoding genes modulated by *Yap1* deletion is displayed. Supp. Figure 3D shows the remaining 2/3 of collagen-encoding genes. FC = fold change. (**B, C**) Representative images (**B**) and quantification (**C**) of ColVI IHC in KP and KPY tumors. Two-tailed unpaired t-tests with Welch correction and Holm-Sidak multiple comparisons test (n = 3-5 mice per genotype with 3 sections per mouse). (**D**) qRT-PCR of *Col6a1*, *Col6a2*, *Col6a3,* and *Yap1* gene expression in KP cells expressing a control or one of multiple independent *Yap1-*targeting shRNAs. One-way ANOVA with Dunnett’s (vs. shScr) for each gene. (**E**) Representative immunoblot of KP cells treated as in **D**. (**F**) Representative images of ColVI IHC in KP tumor-bearing mice treated with 25 mg/kg SAHA + 50 mg/kg JQ1 or vehicle control for 20 days. Quantification is in Supp. Figure 3M. (**G**) Representative immunoblot of KP cells treated with SAHA (2 μM) + JQ1 (0.5 μM) or vehicle control for 48 hours. (**H**) Schematic of experimental model to assess immunomodulatory role of UPS cell-derived decellularized extracellular matrix (dECM). (**I**) Representative widefield images and quantification of ColVI deposition in dECM from KP cells expressing control or *Yap1*-targeting shRNAs. One-way ANOVA with Dunnett’s (vs. shScr). Scale bars = 25 μM. Image brightness and contrast were adjusted for publication. (**J**) Quantification and representative contour plots showing IFNγ and TNFα co-expression in CD44^+^CD8^+^ T cells incubated on dECMs from control and shYap1-expressing KP cells. Each point represents T cells isolated from an individual mouse. One-way ANOVA with Dunnett’s (vs. shScr). The shScr plot and data are identical to those shown in Figure 5A (performed in the same experiment).

To confirm the relationship between Yap1 and ColVI in UPS with a pharmacological approach, we treated tumor-bearing KP mice and KP cells in vitro with a combination of the histone deacetylase inhibitor (HDACi) Vorinostat (also known as Suberoylanilide Hydroxamic Acid; SAHA) and the BRD4 inhibitor JQ1, or vehicle control. We and others have reported that treatment with JQ1/SAHA (or other HDACi) inhibits Yap1 expression in UPS and other cancers (33, 50–52). Microarray, IHC, immunoblotting, and qRT-PCR analyses confirmed that ColVI gene and protein expression were substantially downregulated in SAHA/JQ1-treated cells and tumors (**Figure 2F, G, Supp.** Figure 3K-M). Taken together, these data support the conclusion that UPS-cell intrinsic Yap1 promotes aberrant ColVI deposition in the TME, and that genetic and non-specific pharmacologic inhibition of Yap1 can reverse this process.

As a transcriptional co-activator, Yap1 lacks a DNA-binding domain and must interact with Tea Domain (TEAD) family transcription factors to stimulate gene expression; Tead1 is enriched in skeletal muscle tissue (53) and potentially muscle-derived tumors. Therefore, to explore the mechanism by which Yap1 promotes ColVI deposition, we leveraged publicly available Tead1 ChIP-seq data (GSE55186) from Yap1-driven embryonal rhabdomyosarcoma (eRMS) (54), skeletal muscle-derived tumors that lie on morphological and transcriptional continua with UPS (55). In murine Yap1-driven eRMS (54), Tead1-ChIP signal was enriched in a region that overlapped with the *Col6a1* 5’ untranslated region (UTR), likely corresponding to the *Col6a1* promoter (Supp. Figure 4A). A second peak ∼5 kb upstream of the *Col6a1* 5’ UTR was also observed, potentially representing an enhancer region. Similarly, in cultured human eRMS cells (RD cells), TEAD1-ChIP signal was enriched ∼9 kb upstream of the *COL6A1* 5’ UTR (Supp. Figure 4B). These data suggest that transcriptionally active Yap1 upregulates ColVI deposition in skeletal muscle-derived sarcomas by directly stimulating *Col6a1* transcription.

### Yap1-mediated ColVI deposition promotes CD8^+^ T cell dysfunction

Based on our findings, we hypothesized that Yap1-mediated ColVI deposition in the UPS TME promotes CD8^+^ T cell inhibitory marker expression and dysfunction. To test this idea, we developed a novel system in which C57BL/6 KP cells (B6-KP cells) were seeded at 90% confluency and cultured under hypoxic conditions (1% O_2_), stimulating them to deposit ECM. This ECM was then decellularized (decellularized ECM; dECM) and incubated with activated syngeneic CD8^+^ T cells (splenocytes; C57BL/6 background) under normoxic conditions (21% O_2_) (**Figure 2H**). dECMs were generated under hypoxic conditions because it is well-established that hypoxia stimulates robust ECM gene/protein expression and matrix remodeling in the TME (4, 56); indeed, ColVI deposition was significantly increased in dECMs generated under hypoxia vs. normoxia (Supp. Figure 4C**, D**). Thus, UPS dECMs generated under hypoxic conditions were used in all experiments given our focus on Yap1-mediated ECM deposition and not the role of hypoxia vs. normoxia *per se*. Subsequent CD8^+^ T cell culture on dECMs was conducted under normoxic conditions given previous reports that hypoxia can either enhance or suppress CD8^+^ T cell expansion and function depending on tissue/experimental context and extent of T cell receptor (CD3) stimulation (57–60).

To determine if Yap1-mediated ColVI deposition in UPS enhances T cell dysfunction, we first generated dECMs from control and *Yap1*-deficient B6-KP cells. Immunofluorescent staining revealed that ColVI deposition was somewhat heterogeneous, but was generally decreased in *Yap1-*deficient dECMs compared to controls, confirming the regulatory role of UPS cell-intrinsic Yap1 in ColVI secretion (**Figure 2I**). Although the observed reductions in ColVI were modest and only attained statistical significance for 1 shRNA, these results were unsurprising because culturing cells at high confluence – which is required for matrix deposition in our system – is a well-established Yap1 suppressor (61) and can thereby minimize differences in Yap1 activity between shScr and shYap1 UPS cells. We cultured syngeneic CD8^+^ T cells on these dECMs and measured the surface expression of the T cell inhibitory receptors Pd1 and Tim-3 by flow cytometry. The proportion of CD8^+^ T cells co-expressing Pd1 and Tim-3 was modestly reduced following culture on dECMs from *Yap1*-deficient compared to control UPS cells (Supp. Figure 4E**, F**). We also observed significantly higher percentages of CD8^+^ T cells co-expressing the cytolytic markers IFNγ and TNFα following culture on dECMs from shYap1 cells (**Figure 2J**), consistent with the results of our CART-TnMUC1 assay (**Figure 1J)**. Together, these results support the conclusion that UPS cell-intrinsic Yap1 promotes CD8^+^ T cell dysfunction.

We then determined the specific effects of ColVI, downstream of Yap1, on CD8^+^ T cell surface marker expression by generating dECMs from ColVI-deficient KP cells (**Figure 3A**). In this assay, we targeted *Col6a1,* rather than other ColVI-encoding genes, because *Col6a1* is indispensable for ColVI protein synthesis (48). *Col6a1* depletion in KP cells significantly reduced CD8^+^ T cell dysfunction in this assay as measured by co-expression of Pd1 and Tim-3 (**Figure 3B, C**). We also confirmed that ColVI depletion does not affect KP tumor-derived cell proliferation in vitro (Supp. Figure 5A**, B**), consistent with the hypothesis that the dominant role of ColVI is specific to immune modulation in the TME. To directly test the effect of COLVI on T cell-mediated killing, we employed the human CART-TnMUC1 system introduced in **Figure 1J** (44). Longitudinal T cell-mediated cytolysis of STS-109 UPS cells expressing control or *COL6A1*-specific shRNAs revealed that COLVI depletion enhanced cytotoxic T cell function, phenocopying the effects of *YAP1* depletion (**Figure 3D, E**, Supp. Figure 5C). To address the possibility that shCOL6A1 UPS cells (and shYAP1 cells in **Figure 1J**) are simply more susceptible than shScr cells to T cell-mediated apoptosis, we treated them with recombinant human TNFα or IFNγ, two cytolytic cytokines known to be produced by CD8^+^ T cells, and evaluated apoptosis by flow cytometry. We reasoned that equivalent doses of purified cytokines should elicit similar levels of apoptosis in shYAP1/shCOL6A1 cells and controls if CD8^+^ T cell function is truly enhanced in the setting of UPS cell-intrinsic *YAP1* or *COL6A1* deficiency. We observed that purified cytokines did not increase shYAP1/shCOL6A1 UPS cell apoptosis relative to shScr cells, confirming enhanced CART cell cytotoxicity in the presence of reduced UPS cell-derived COLVI (Supp. Figure 5D**, E**). In fact, IFNγ elicited lower rates of late apoptosis in shYAP1 and shCOL6A1 UPS cells compared to controls; however, these effect sizes were modest and inconsistent across cytokine concentrations and independent shRNAs. Consistent with these results, COLVI protein was detected extracellularly and in UPS cell culture-conditioned medium (Supp. Figure 5F-H), where it can suppress CART-TnMUC1-mediated cytolysis and promote T cell dysfunction.

**Figure 3.**
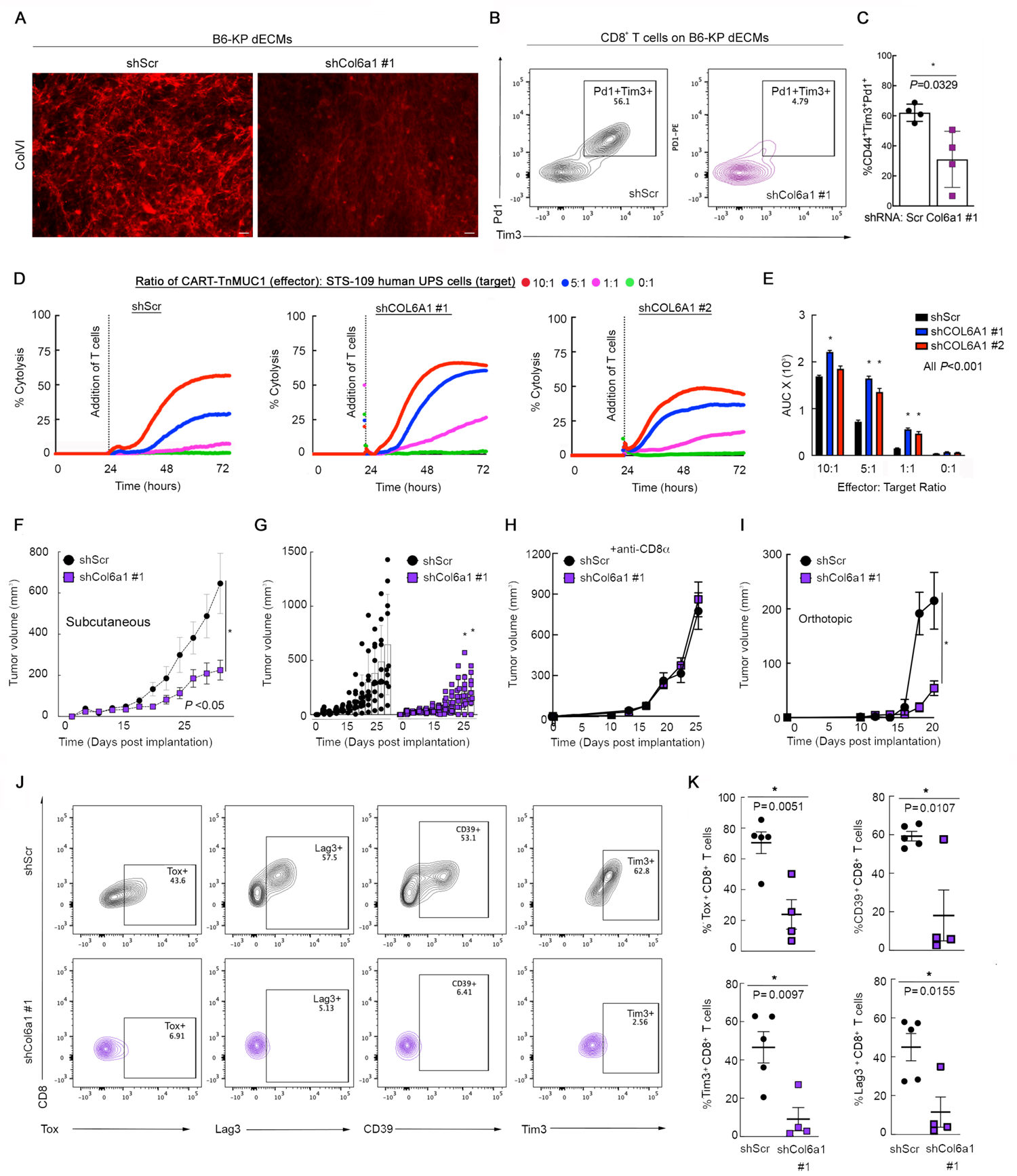
ColVI in the UPS TME promotes CD8^+^ T cell dysfunction. (**A**) Representative widefield images of ColVI immunofluorescence in dECMs generated from control and shCol6a1-expressing B6-KP cells. Scale bar = 20 μm. Image brightness and contrast were adjusted for publication. (**B**) Representative contour plots and (**C**) quantification of Pd1 and Tim-3 co-expression in CD44^+^CD8^+^ T cells incubated on dECM derived from control or shCol6a1 KP cells. (**D**) Longitudinal cytolysis of shScr or shCOL6A1-expressing human STS-109 UPS cells co-cultured with CART-TnMUC1 cells. Measurements indicate % target cell (UPS cell) cytolysis. Normalized cell index for STS-109 cells alone (not shown) controls for differences in target cell growth across conditions. (**E**) Quantification of area-under-the-curve (AUC) from **D**; one-way ANOVA with Dunnett’s (vs. shScr) for each ratio. In **D, E**, shScr data are identical to those in Fig. 1J (performed in the same experiment). (**F**) Tumor growth curves from subcutaneous (flank) syngeneic transplant of 3 x 10^4^ B6-KP cells in Matrigel expressing control or *Col6a1*-targeting shRNAs in syngeneic C57BL/6 mice. Two-way ANOVA. (**G**) Visualization of individual tumors from **F**. (**H**) Tumor growth curves depicting subcutaneous (flank) syngeneic transplant of 5 x 10^5^ KP cells (SKPY42.1 cell line) expressing control or *Col6a1*-targeting shRNAs in C57BL/6 mice treated with α-CD8α every three days. (**I**) Tumor growth curves depicting syngeneic orthotopic transplant (into the gastrocnemius muscle) of 2.5 x 10^5^ KP cells (SKPY42.1 cell line) expressing control or *Col6a1*-targeting shRNAs in C57BL/5 mice. Two-way repeated-measures ANOVA, SEM. (**J, K**) Representative contour plots (**J**) and quantification (**K**) of T cell dysfunction markers in CD8^+^ T cells from control and shCol6a1 orthotopic tumors from **I**. Each point in **K** represents an individual tumor. Two-tailed unpaired t-tests.

In light of our in vitro findings that ColVI suppresses CD8^+^ T cell function, we investigated this relationship in vivo by generating control and *Col6a1* shRNA-expressing UPS tumors (syngeneic allograft of SKPY42.1 KP cells on a pure C57BL/6 background) in C57BL/6 hosts. In these immunocompetent mice, ColVI-deficient tumors were significantly smaller and slower growing than control tumors (**Figure 3F, G**). We demonstrated that ColVI-dependent tumor growth is mediated by T cell inactivation by depleting CD8^+^ T cells in the syngeneic transplant system (**Figure 3H**, Supp. Figure 5I), where control and shCol6a1 tumors grew at the same rate. We also generated syngeneic orthotopic tumors by injecting control and shCol6a1-expressing KP cells into the gastrocnemius muscles of immunocompetent C57BL/6 mice (**Figure 3I**, Supp. Figure 5J). Flow cytometric analysis indicated that the proportion of CD8^+^ T cells expressing dysfunction markers, including Tox, Tim-3, CD39, and Lag3, was significantly decreased in ColVI-deficient tumors compared to controls (**Figure 3J, K**). Taken together, these findings confirm that Yap1-mediated ColVI deposition in the UPS TME promotes CD8^+^ T cell dysfunction and immune evasion.

### ColVI colocalizes with and remodels collagen I fibers in the UPS TME

Next, we explored the mechanism by which COLVI promotes CD8^+^ T cell dysfunction in the UPS TME. We first asked whether CD8^+^ T cell dysfunction is induced following direct interaction with deposited COLVI via known COLVI receptors. To test this hypothesis, we developed a second novel in vitro system by incorporating purified human COLVI into COLI-containing hydrogels. COLI, a fibrillar collagen, is one of the most widely used hydrogel scaffolds due to its mechanical stability, hydrophilicity, versatility, in vivo abundance, and ease of extraction (62). COLVI, which forms microfilaments instead of fibers, does not possess all of these properties and thus cannot be used to generate hydrogels independently; moreover, unlike COLI, which can be found in isolation and need not interact with other matrix proteins in vivo, COLVI is always found bound to other ECM molecules and/or cell surface proteins (63). Activated human CD8^+^ T cells were then cultured on these COLVI-containing hydrogels, allowing us to assess the impact of purified matrix proteins on CD8^+^ T cells in a three-dimensional environment. Subsequently, we genetically or pharmacologically blocked the known COLVI receptors ITGB1, NG2 (CSPG4), CMG2, and ITGAV (Supp. Figure 6A). ITGAV and ITGB1 can bind multiple collagen species, including both COLVI and COLI; however, to our knowledge, NG2 and CMG2 are specific for COLVI (63–68). Neutralization of ITGB1 or NG2 with blocking antibodies had no effect on CD8^+^ T cell dysfunction as measured by co-expression of TIM-3 and PD1, nor did it restore CD8^+^ T cell proliferation (KI67 positivity). Similar results were obtained following treatment of human CD8^+^ T cells with cilengitide, a selective inhibitor of αvβ3 and αvβ5 integrins (69), and with activated CD8^+^ T cells from *Cmg2*^-/-^ mice (70) (Supp. Figure 6B-F). From these results, we conclude that CD8^+^ T cell dysfunction is not strongly modulated by canonical COLVI receptors. However, we cannot exclude the possibility that ITGAV and/or ITGB1 may be involved, as neutralization of these receptors would likely block both T cell-COLVI and T cell-COLI interactions in our hydrogel system.

In the absence of a direct mechanism connecting ColVI receptors to T cell dysfunction, we investigated potential indirect mechanisms. ColVI binds to a number of ECM proteins, including fibrillar collagens such as ColI, one of the most prevalent collagens in mammalian tissues (71–73). In one study of resting peripheral blood T cells, COLI induced CD8^+^ T cell proliferation in vitro when used in conjunction with CD3/T cell receptor stimulation (74). Therefore, we hypothesized that Yap1-mediated ColVI deposition promotes CD8^+^ T cell dysfunction by altering ColI content and/or organization in the UPS ECM. We first examined the effect of *Col6a1* depletion on ColI levels in KP cells in vitro. qRT-PCR demonstrated no consistent changes in ColI-related gene expression, whereas immunoblot, dot blot, and co-immunofluorescence assays indicated that ColI protein levels generally did not change in *Col6a1*-deficient vs. control cells (Supp. Figure 7A-G). However, we did observe marked colocalization between ColI and ColVI in control (shScr) KP dECMs, with 26.4% of ColI colocalizing with ColVI, indicating a physical interaction between these two ECM components (**Figure 4A, Supp.** Figure 7H**, Mov. S1-3**). Therefore, we considered the possibility that ColVI remodels ColI in the UPS TME, with potential implications for CD8^+^ T cell function.

**Figure 4.**
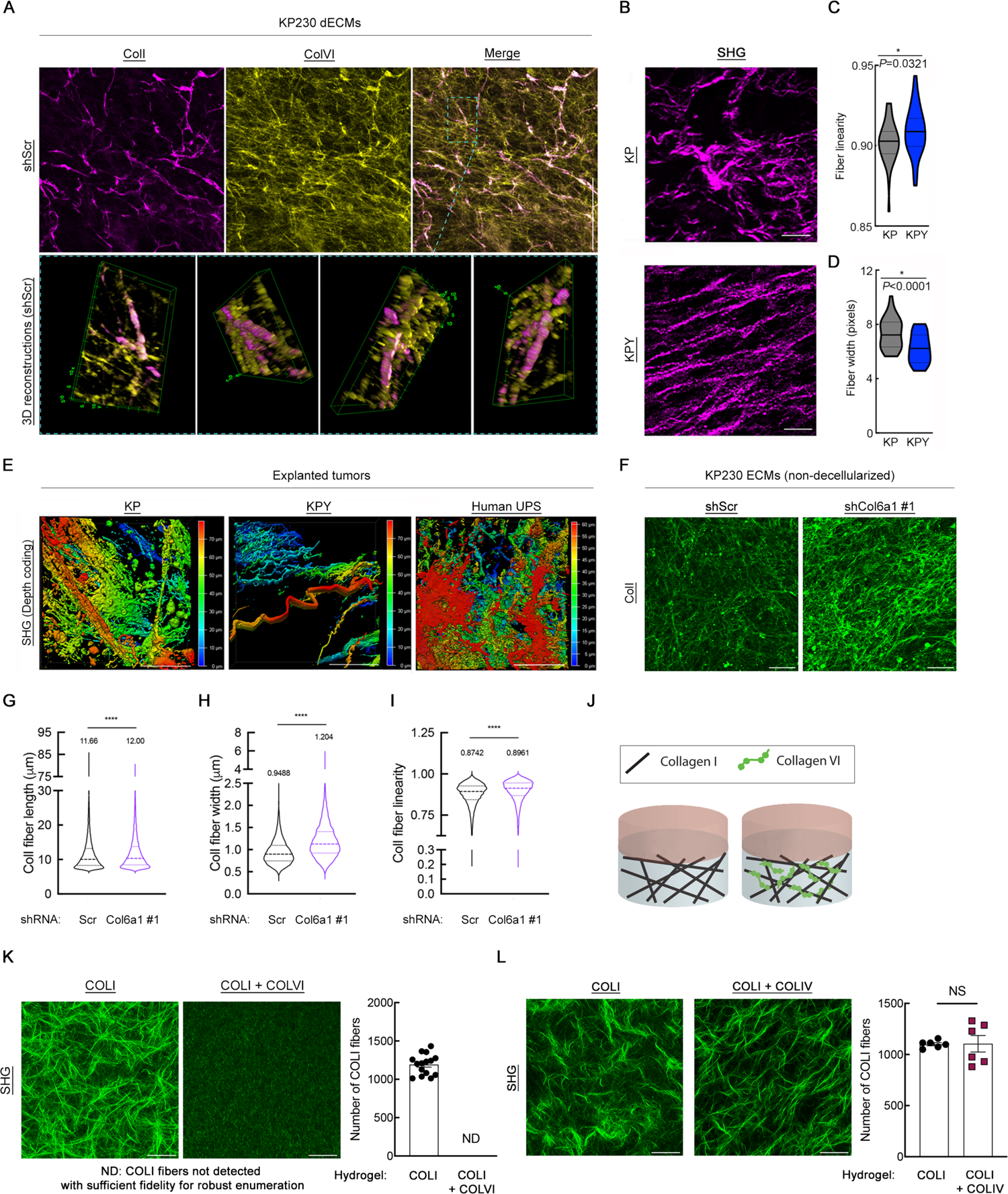
ColVI interacts with and remodels ColI in the UPS TME. (**A**) Representative confocal micrographs (maximum-intensity Z-projections) and 3D reconstructions showing ColI and ColVI co-immunofluorescence in KP cell-derived dECMs. Scale bar = 100 μM (**B**) Representative multiphoton second-harmonic generation (SHG) images (maximum-intensity Z-projections) of KP and KPY tumor sections. Scale bars = 50 μM. (**C, D**) Violin plots of CT-FIRE analysis of images from **B**. Mean fiber width and linearity were plotted for ≥5 separate fields (n = 5 mice per genotype); two-tailed unpaired t-test. (**E**) Representative depth-coded SHG images of human UPS, KP, and KPY explanted live tumors. Red = SHG signal farthest from the objective/greatest relative tissue depth, blue = SHG signal closest to the objective/shallowest relative tissue depth. Scale bars = 50 μM. (**F**) Representative confocal micrographs (maximum-intensity Z-projections) of ColI immunofluorescence in ECMs (non-decellularized) generated from control and shCol6a1 KP cells. Scale bars = 50 μM. (**G-I**) Violin plots depicting CT-FIRE analysis of images in **F**. ColI fiber length (**G**), width (**H**), and linearity (**I**) were plotted from 7 independent fields across multiple dECMs per condition. Numbers above violin plots indicate means. Thick and thin dotted lines within the shapes denote medians and quartile 1/3, respectively. (**J**) Schematic of in vitro hydrogel system to assess how purified COLVI impacts purified COLI structure/organization. (**K**) Representative SHG images (maximum-intensity Z-projections with 2x optical zoom) and quantification of COLI fiber number in COLI-alone and COLI + COLVI hydrogels. Scale bars = 50 μM. (**L**) Representative SHG images (maximum-intensity Z-projections with 2x optical zoom) and quantification of COLI fiber number in COLI-alone and COLI + COLIV hydrogels. Scale bars = 50 μM. For **K-L**, quantification was performed for ≥6 independent fields across multiple hydrogels per condition. Brightness and contrast of all micrographs in Figure 4 were adjusted for publication.

To test this hypothesis, we began by examining the architecture of fibrillar collagen molecules in explanted GEMM tumors. Using multiphoton second harmonic generation (SHG) imaging, we identified significant alterations to fibrillar collagen organization, including significantly thinner and straighter fibers in KPY tumors compared to KP (**Figure 4B-D**). Importantly, these changes in fibrillar collagen structure occurred despite similar levels of ColI-related gene expression (**Figure 2A**, Supp. Figure 3D**, 8A**) and ColI protein deposition (Supp. Figure 8B**, C**) in KP and KPY tumors. Similar results were observed in SAHA/JQ1-treated tumors compared to controls (Supp. Figure 8D-F). We also evaluated human UPS tumors and found that their fibrillar collagen structure recapitulated that of KP tumors, confirming that our GEMMs successfully reproduce this aspect of tumor biology (**Figure 4E**, **Mov. S4-6**). To explore the impact of ColVI on ColI organization more directly, we examined extracellular ColI immunofluorescent staining patterns in control and *Col6a1*-deficient KP cell-derived ECMs (**Figure 4F**). In this experiment, matrices were not decellularized in order to circumvent the potential (albeit minor) changes in ECM structure induced by the decellularization process. We observed that ColI fibers in shCol6a1 ECMs were significantly longer, straighter, and wider than those in shScr ECMs, and exhibited significantly different orientation distributions, confirming ColVI-mediated remodeling (**Figure 4G-I, Supp.** Figure 8G). Finally, we asked whether COLVI alters COLI structure directly or indirectly through other mechanisms by performing SHG imaging of hydrogels containing purified COLI alone, or COLI together with purified COLVI (**Figure 4J**). As SHG can only detect fibrillar collagen molecules, COLI, but not COLVI, is imaged in this assay. Remarkably, the addition of COLVI (250 μg/mL) to our hydrogel system nearly abolished the formation of COLI fibers and higher-level structures (e.g., fiber bundles; **Figure 4K**). In contrast, COLI fibers remained abundant in the presence of a different non-fibrillar collagen, collagen type IV (COLIV; used at the same concentration as COLVI), underscoring the potential specificity of the COLI-COLVI relationship (**Figure 4L**). Taken together, we conclude that ColVI directly modifies ColI fiber architecture in the UPS TME.

### ColI opposes ColVI and abrogates CD8^+^ T cell dysfunction

We next sought to understand mechanistically how ColI-ColVI interactions impact CD8^+^ T cells, hypothesizing that ColVI triggers dysfunction indirectly by remodeling ColI in the TME. To test the impact of ColI on CD8^+^ T cell function, we incubated activated murine CD8^+^ T cells on dECMs from control or *Col1a1*-deficient B6-KP cells. Unlike in the setting of *Yap1* and *Col6a1* deficiency (**Figure 2, 3**), the proportion of IFNγ^+^TNFα^+^ CD8^+^ T cells was reduced following culture on *Col1a1-*deficient dECMs, indicating decreased cytolytic capacity (**Figure 5A, B**). We then compared the effects of COLI and COLVI on activated human CD8^+^ T cells by culturing them on hydrogels containing purified COLI alone, or COLI together with purified COLVI (**Figure 5C)**. The proportion of CD8^+^ T cells co-expressing PD1 and TIM-3 was significantly reduced on COLI gels compared to on COLVI-containing hydrogels (**Figure 5D, E**); TIM-3 median fluorescence intensity was similarly decreased (Supp. Figure 9A). Additionally, CD8^+^ T cell proliferative capacity was improved on COLI gels relative to COLI + COLVI gels as indicated by greater KI67 positivity (**Figure 5F, G**). The proportion of cytolytic IFNγ^+^TNFα^+^ CD8^+^ T cells was also modestly elevated in the presence of COLI alone (Supp. Figure 9B). We then tested the specificity of COLI-COLVI interactions on CD8^+^ T cell function by substituting COLIV for COLVI in this assay. Remarkably, CD8^+^ T cell dysfunction was not significantly impacted by the addition of COLIV to COLI-containing hydrogels (Supp. Figure 9C), consistent with our SHG data (**Figure 4L**), and further illustrating the specificity of the COLI-

**Figure 5.**
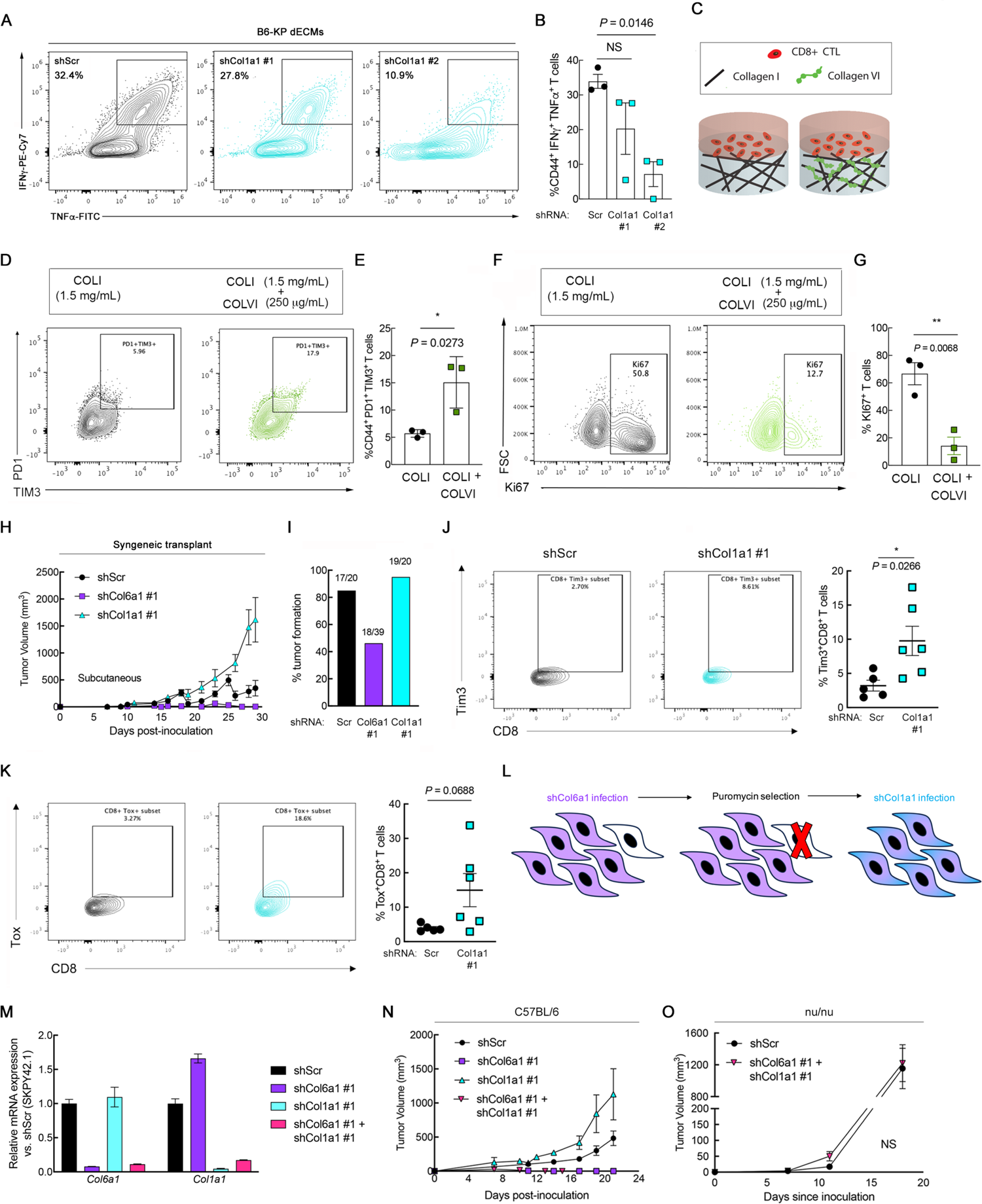
ColVI-mediated CD8^+^ T cell dysfunction is restored in the presence of ColI. (**A**) Representative flow cytometry plots and (**B**) quantification of IFNγ and TNFα co-expression in CD44^+^CD8^+^ T cells incubated on dECMs from control or shCol1a1 KP cells. Each point represents T cells from an individual mouse. One-way ANOVA with Dunnett’s vs. shScr. The shScr plot and data are identical to those in Figure 2J (performed in the same experiment). (**C**) Schematic of in vitro hydrogel system to test how ColI impacts ColVI-mediated CD8^+^ T cell dysfunction. CTL = cytotoxic T lymphocyte. (**D, E**) Representative flow cytometry plots (**D**) and quantification (**E**) of activated human CD8^+^CD44^+^ T cells showing TIM-3 and PD1 co-expression after incubation on hydrogels containing purified COLI with or without purified COLVI. Two-tailed unpaired t-test. (**F, G**) Representative flow cytometry plots (**F**) and quantification (**G**) of KI67 in activated human CD8^+^CD44^+^ T cells cultured as in **D, E**. Two-tailed unpaired t-test. (**H**) Tumor growth curves from subcutaneous (flank) syngeneic transplant of 5 x 10^5^ KP cells (SKPY42.1 cell line) expressing control, *Col6a1*, or *Col1a1*-targeting shRNAs in C57BL/6 mice. Data are from two independent animal cohorts (total n = 20 for shScr and shCol1a1; n = 39 for shCol6a1). (**I**) Tumor formation rates from **H**. (**J, K**) Representative contour plots and quantification of Tim-3 (**J**) and Tox (**K**) expression in CD8^+^ T cells in tumors from **H**. Each point represents an individual tumor. Two-tailed unpaired t-test with Welch correction. (**L**) Schematic depicting strategy for depleting *Col6a1* and *Col1a1* in the same UPS cell population. (**M**) Validation of *Col6a1* and *Col1a1* expression in UPS cells from **L** (SKPY42.1 cell line). (**N, O**) Tumor growth curves from subcutaneous syngeneic transplant of 1 x 10^6^ KP cells (SKPY42.1 cell line) from **M** in C57BL/6 (**N**) and nu/nu (**O**) mice.

COLVI relationship.

Stiffened, fibrotic microenvironments such as those in desmoplastic solid tumors are well-known to activate Yap1 (38). Therefore, we ascertained whether COLVI drives CD8^+^ T cell dysfunction by increasing ECM stiffness and potentiating Yap1 signaling. We queried the expression and subcellular localization of Yap1 in control, shCol6a1, and shCol1a1 KP cells, but did not detect significant differences in Yap1 gene expression, protein levels, or S127 phosphorylation, an established surrogate for cytoplasmic retention and degradation (Supp. Figure 9D-F). Consistent with this observation, hydrogel stiffness was not significantly altered by the addition of COLVI (Supp. Figure 9G).

We also investigated the involvement of a ColI receptor, Lair1, in CD8^+^ T cell dysfunction, given a recent report that Lair1 negatively regulates CD8^+^ T cell activity and may promote immunotherapy resistance in lung cancer (75). However, analysis of publicly available single-cell RNA-sequencing data from KP UPS tumors (GSE144507; (76)) demonstrated that *Lair1* was predominantly expressed on tumor-associated macrophages and very minimally on CD8^+^ T cells (Supp. Figure 9H). Moreover, *LAIR1* (but not *COL1A1* itself), was associated with improved overall survival in UPS patients (Supp. Figure 9I**, J**), inconsistent with its putative role promoting CD8^+^ T cell immunosuppression (75). These results argue against the involvement of *Lair1* in UPS matrix-mediated immune evasion. Finally, we confirmed that neither molecular diffusion rates throughout, nor oxygen concentrations within, the hydrogels were substantially impacted by the addition of COLVI, demonstrating that differential nutrient and/or oxygen availability likely does not underlie the observed COLVI-induced CD8^+^ T cell dysfunction in this system (Supp. Figure 9K**, L**). Taken together, these observations clearly indicate that COLI can abrogate the CD8^+^ T cell dysfunction mediated by COLVI, and demonstrate the relative importance of ECM signaling over Yap1 hyperactivation in this process.

Our findings thus far suggest that ColVI functions to restrain ColI-mediated CD8^+^ T cell activity and proliferation. To test this hypothesis in vivo, we generated subcutaneous syngeneic tumors by injecting SKPY42.1 cells expressing control, *Col1a1*-, or *Col6a1*-targeting shRNAs in C57BL/6 mice (**Figure 5H**). In this immunocompetent setting, *Col1a1*-deficient tumors grew more rapidly than both control and *Col6a1*-deficient tumors. shCol1a1 and shScr tumors also developed with similar efficiency (85% and 95%, respectively), whereas shCol6a1 tumors only formed in 46.2% of mice (**Figure 5I**). Of the shCol6a1 tumors that did form, 83.3% (15/18) rapidly regressed before they reached 100 mm^3^. Similar results were obtained in immunocompetent syngeneic orthotopic tumor models (Supp. Figure 10A-C). Importantly, shCol1a1-tumor-bearing mice experienced significantly worse survival than mice bearing control tumors, whereas survival of shCol6a1-tumor bearing mice was improved (Supp. Figure 10D). Furthermore, evaluation of dysfunction marker expression on CD8^+^ T cells revealed upregulation of Tim-3 and Tox in *Col1a1*-deficient UPS tumors compared to controls (**Figure 5J, K**), indicative of increased dysfunction due to loss of ColI. Impressively, when we assessed the impact of *Col1a1* depletion on KP cell growth in vitro, *Col1a1*-deficient cells proliferated more slowly than control cells, reflecting a discrepancy between effects of ColI depletion in vitro and in vivo in this system (Supp. Figure 10E). These results confirm that the presence of ColI in the UPS ECM controls tumor growth by enabling host anti-tumor immunity, whereas the aberrant deposition of ColVI opposes ColI and promotes immune evasion.

To ascertain whether ColVI or ColI plays the dominant role in matrix-mediated CD8^+^ T cell dysfunction, we depleted *Col6a1* and *Col1a1* in the same population of SKPY42.1 cells and injected them into recipient syngeneic C57BL/6 mice (**Figure 5L, M**). Cells deficient for both collagens formed tumors at intermediate rates between those deficient for shCol6a1 or shCol1a1 individually (Supp. Figure 10F), but the resulting tumors (herein referred to as “double knockdown tumors”) phenocopied shCol6a1 tumor growth, rapidly regressing before they reached ∼100 mm^3^ (**Figure 5N, Supp.** Figure 10G). To confirm that this observed regression was T cell-dependent, we generated control and double knockdown tumors in nu/nu mice, in which mature T cells are lacking but innate immune cells are present. In this critical experiment, no statistically significant differences in tumor formation or growth were observed (**Figure 5O, Supp.** Figure 10H**, I**), confirming T cell-mediated double knockdown tumor regression in the syngeneic model (**Figure 5N**). From these data, we conclude that ColVI plays a dominant role over that of ColI in UPS matrix-mediated immune evasion.

### ColVI promotes T cell dysfunction by disrupting CD8^+^ T cell autophagic flux

To date, an immunosuppressive role of ColVI has not been documented in tumors. We therefore aimed to identify the specific downstream mechanism by which ColVI deposition causes T cell dysfunction. ColVI most notably regulates mesenchymal cell adhesion and migration (77, 78); however, we were intrigued by its ability to modulate autophagy in fibroblasts and muscle tissue (79). Autophagy is a central regulator of T cell metabolism and is essential for T cell activation (80, 81). To assess whether ColVI impacted CD8^+^ T cell autophagy, we encapsulated T cells in purified COLVI-containing hydrogels and visualized autophagosomes *in situ*. In the presence of COLVI, T cells contained significantly more and brighter autophagosomes than T cells encapsulated in ColI gels (**Figure 6A-C**). To determine whether COLVI caused autophagosome accumulation by inducing autophagy, or by disrupting autophagic flux and autophagosome clearance, we treated activated CD8^+^ T cells with chloroquine (CQ) in the presence or absence of COLVI and evaluated LC3B-II expression. In murine T cells, inhibiting autophagic flux with CQ did not further increase Lc3b-II, indicating that COLVI disrupts autophagic flux, but does not induce autophagy (**Figure 6D, E**). Activated human peripheral blood CD8^+^ T cells generally showed the same trend (**Figure 6F-I**). We confirmed this result by staining for p62, a protein rapidly degraded during autophagy induction. In the presence of COLVI, p62 accumulated within T cells as evidenced by increased mean p62 signal intensity (**Figure 6J, K**). Together, these results indicate that extracellular COLVI inhibits autophagic flux in CD8^+^ T cells.

**Figure 6.**
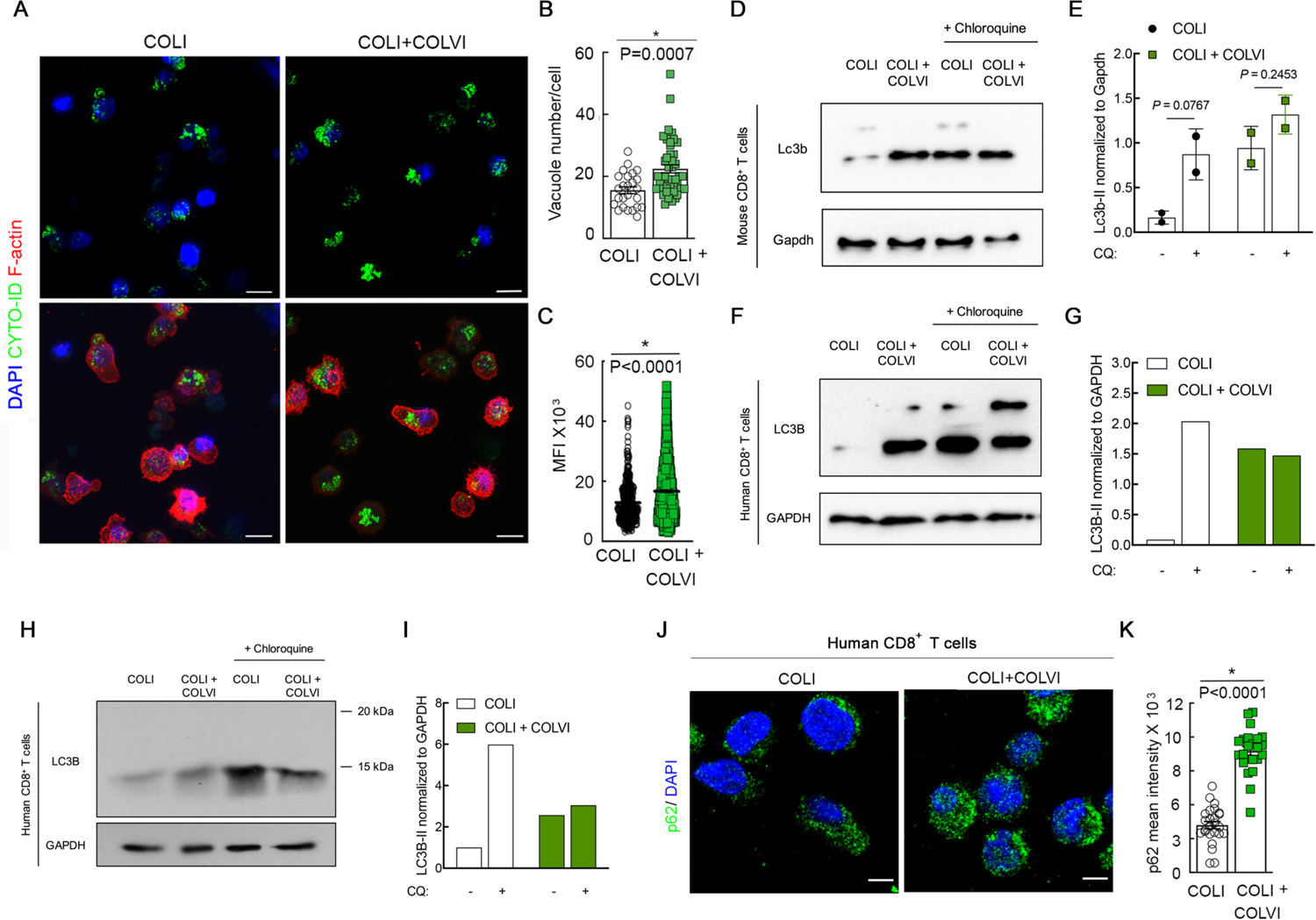
ColVI disrupts CD8^+^ T cell autophagic flux. (**A-C**) Visualization (**A**) and quantification (**B, C**) of autophagosomes in CD8^+^ human T cells cultured on hydrogels containing purified COLI with or without purified COLVI. Two-tailed unpaired t-test. (**D, E**) Western blot (**D**) and quantification (**E**) of Lc3b-II expression in murine CD8^+^ T cells cultured on purified COLI-containing hydrogels in the presence or absence of purified COLVI, with or without chloroquine (CQ) treatment. Two-tailed unpaired t-test. n = 2. SD. (**F, G**) Western blot (**F**) and quantification (**G**) of LC3B-II expression in human CD8^+^ T cells cultured on purified COLI-containing hydrogels in the presence or absence of purified COLVI, with or without CQ treatment. (**H, I**) Western blot (**H**) and quantification (**I**) of LC3B-II expression in human CD8^+^ T cells cultured on purified COLI-containing hydrogels in the presence or absence of purified COLVI, with or without CQ treatment. Molecular weight marker positions are shown to demonstrate that the single LC3B band detected in this experiment corresponds to the reported molecular weight for LC3B-II (14-16 kDa for LC3B-II vs. 16-18 kDa for LC3B-I). Samples from **F, G** and **H, I** were generated from cells from different donors, and shown separately because the analyses were conducted at different institutions using different detection methods (digital fluorescent detection vs. chemiluminescence on film). (**J, K**) Representative images (**J**) and quantification (**K**) of p62 immunofluorescence in human CD8^+^ T cells cultured on purified COLI-containing hydrogels with or without purified COLVI. Two-tailed unpaired t-test.

To explore the broader impacts of our findings, we next considered whether COLVI might also impact CD8^+^ T cell function in other cancer types. Analysis of TCGA PanCancer RNA-seq data revealed that, like in sarcomas, *COL6A1*, *COL6A2*, and *COL6A3* are highly expressed in pancreatic ductal adenocarcinoma (PDAC) (Supp. Figure 11A). Indeed, PDAC tumors contain abundant desmoplastic stroma, most of which is secreted by cancer-associated fibroblasts (CAFs) in the TME (24). Therefore, we generated dECMs from PDAC-CAFs isolated from three independent human tumors and confirmed their ability to secrete COLVI (Supp. Figure 11B). We then generated dECMs from control and shCOL6A1-expressing PDAC-CAFs and incubated them with activated human CD8^+^ T cells. Surprisingly, the proportion of CD8^+^ T cells co-expressing PD1 and TIM-3^+^ was not altered by exposure to *COL6A1*-deficient vs. -replete PDAC-CAF-derived matrix (Supp. Figure 11C**, D**). Thus, unlike in UPS, COLVI does not appear to modulate CD8^+^ T cell function in PDAC. Given a recent report that ColI produced by fibroblasts (containing Col1a1/Col1a2 heterotrimers) is distinct from that produced by PDAC cancer cells (containing Col1a1 homotrimers that possess oncogenic properties) (82), we then asked whether the differential immunomodulatory capacity of CAF-derived vs. UPS-derived ECM resulted from the production of heterotrimeric vs. homotrimeric ColI, respectively. However, like fibroblasts (and unlike PDAC cancer cells) (82), *Col6a1*-sufficient and -deficient UPS cells secreted both Col1a1 and Col1a2 (Supp. Figure 11E), demonstrating that differences in the composition of ColI trimers do not likely underlie the divergent effects of CAF-derived vs. UPS-derived matrix on CD8^+^ T cell function. Taken together, these data underscore critical differences in matrix protein composition between sarcomas and carcinomas, and highlight the potential specificity of the ColVI-CD8^+^ T cell relationship to mesenchymal tumors.

### COLVI as a potential prognostic and diagnostic biomarker in human STS

To evaluate our experimental findings in UPS in the clinical setting, we used data from multiple human UPS patient cohorts: the Detwiller et al. dataset (83), TCGA-Sarcoma, and surgical specimens from the Hospital of the University of Pennsylvania (HUP). Like *YAP1* (33), *COL6A1*, *COL6A2*, and *COL6A3* were upregulated in human UPS relative to normal muscle tissue and strongly correlated with poor outcome in UPS patients (**Figure 7A-I**). We also interrogated the relationship between YAP1 and COLVI in these datasets. Consistent with our in vitro and GEMM data demonstrating that Yap1 promotes ColVI deposition in the UPS TME, COLVI expression highly correlated with nuclear YAP1 staining, a surrogate for YAP1 transcriptional activity (**Figure 7J, K**). Moreover, the *COL6A1*, *COL6A2*, and *COL6A3* promoters appeared transcriptionally active in these specimens based upon the presence of H3K27Ac marks at these loci (**Figure 7L**). Finally, *COL6A3* gene expression positively tracked with that of *YAP1* and *FOXM1* (**Figure 7M, N**), the latter of which is a *YAP1* target gene in sarcoma.

**Figure 7.**
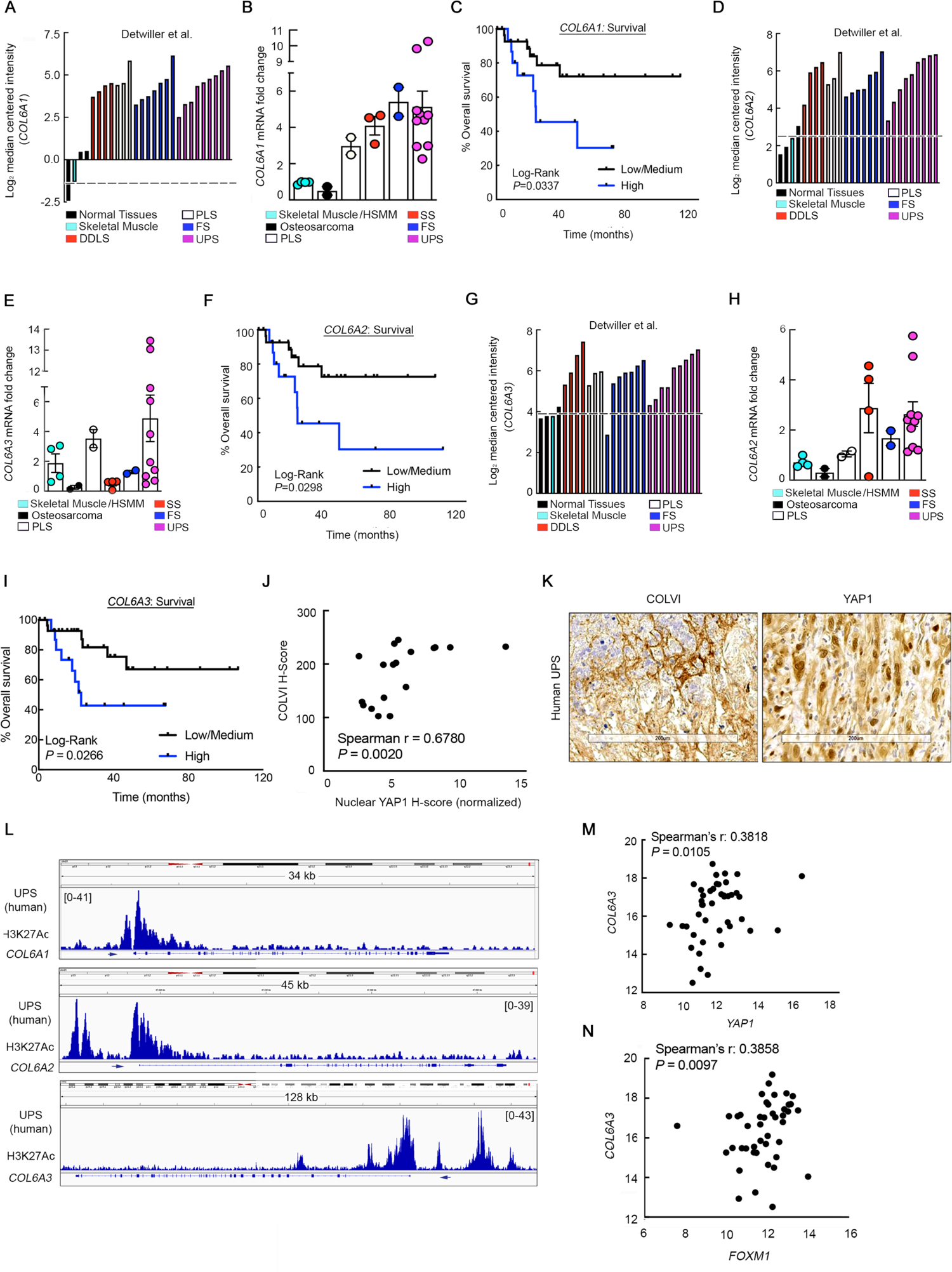
YAP1 and COLVI expression/activity are correlated in human UPS tumors. (**A**) *COL6A1* gene expression levels in specimens from the Detwiller sarcoma dataset (Oncomine) (83). DDLS = dedifferentiated liposarcoma, PLS = pleomorphic liposarcoma, FS = fibrosarcoma. (**B**) qRT-PCR analysis of *COL6A1* expression in human sarcoma and normal skeletal muscle tissue specimens (Hospital of the University of Pennsylvania; HUP). PLS = pleomorphic liposarcoma, SS = synovial sarcoma, FS = fibrosarcoma. (**C**) Kaplan-Meier overall survival curves of UPS patients in TCGA-Sarcoma (n = 44) stratified by intratumoral *COL6A1* gene expression levels. (**D**) *COL6A2* gene expression levels in specimens from the Detwiller sarcoma dataset. (**E**) qRT-PCR analysis of *COL6A2* expression in human sarcoma and normal skeletal muscle tissue specimens (HUP). (**F**) Kaplan-Meier overall survival curves of UPS patients in TCGA-Sarcoma stratified by intratumoral *COL6A2* gene expression levels. (**G**) *COL6A3* gene expression levels in specimens from the Detwiller sarcoma dataset. (**H**) qRT-PCR analysis of *COL6A3* expression in human sarcoma and normal skeletal muscle tissue specimens (HUP). (**I**) Kaplan-Meier overall survival curves of UPS patients in TCGA-Sarcoma dataset stratified by intratumoral *COL6A3* gene expression levels. (**J**) Correlation of COLVI and nuclear YAP1 immunostaining in UPS tumor specimens (HUP). Each point represents an individual specimen. (**K**) Representative IHC images from **J**. Scale bar = 200 μm. (**L**) Publicly available ChIP-seq data (GSE97295) of *COL6A1*, *COL6A2*, and *COL6A3* promoter H3K27 acetylation in human UPS samples (HUP). (**M, N**) Correlation of *YAP1* with *COL6A3* (**M**) and *FOXM1* (**N**) gene expression in UPS tumors from TCGA-Sarcoma.

Finally, we explored the relationship between COLVI expression in UPS and other sarcoma subtypes using a sarcoma tissue microarray (TMA). UPS tumors exhibited the highest mean COLVI H-score of all tumor types (88.52; range: 105.73), significantly greater than that of leiomyosarcomas, neurofibromas, and synovial sarcomas (**Figure 8A, B**). Importantly, the dynamic range of COLVI staining in the UPS TMA cohort was similar to that observed in the HUP cohort (**Figure 7J-K**, range: 142.94). Given that UPS is more prevalent among older adults and presents with aggressive clinical features (29), associations between COLVI H-score and histology were then adjusted for patient age, tumor grade, and tumor stage (**Figure 8A**). After controlling for these variables, the relationships between COLVI H-score and histologic subtype were attenuated but remained statistically significant. Tumor grade was the only other variable that exhibited significant associations with COLVI H-score in univariate models. Furthermore, in TCGA dataset, *COL6A1* gene expression was significantly associated with reduced long-term survival among liposarcoma patients, where tumor COLVI expression levels are similar to those in UPS, but not leiomyosarcoma patients, where tumor COLVI levels are significantly lower (**Figure 8C, D**). These data indicate that COLVI expression may be a biomarker of long-term clinical outcomes and sensitivity to immunotherapy in some human sarcoma patients.

**Figure 8.**
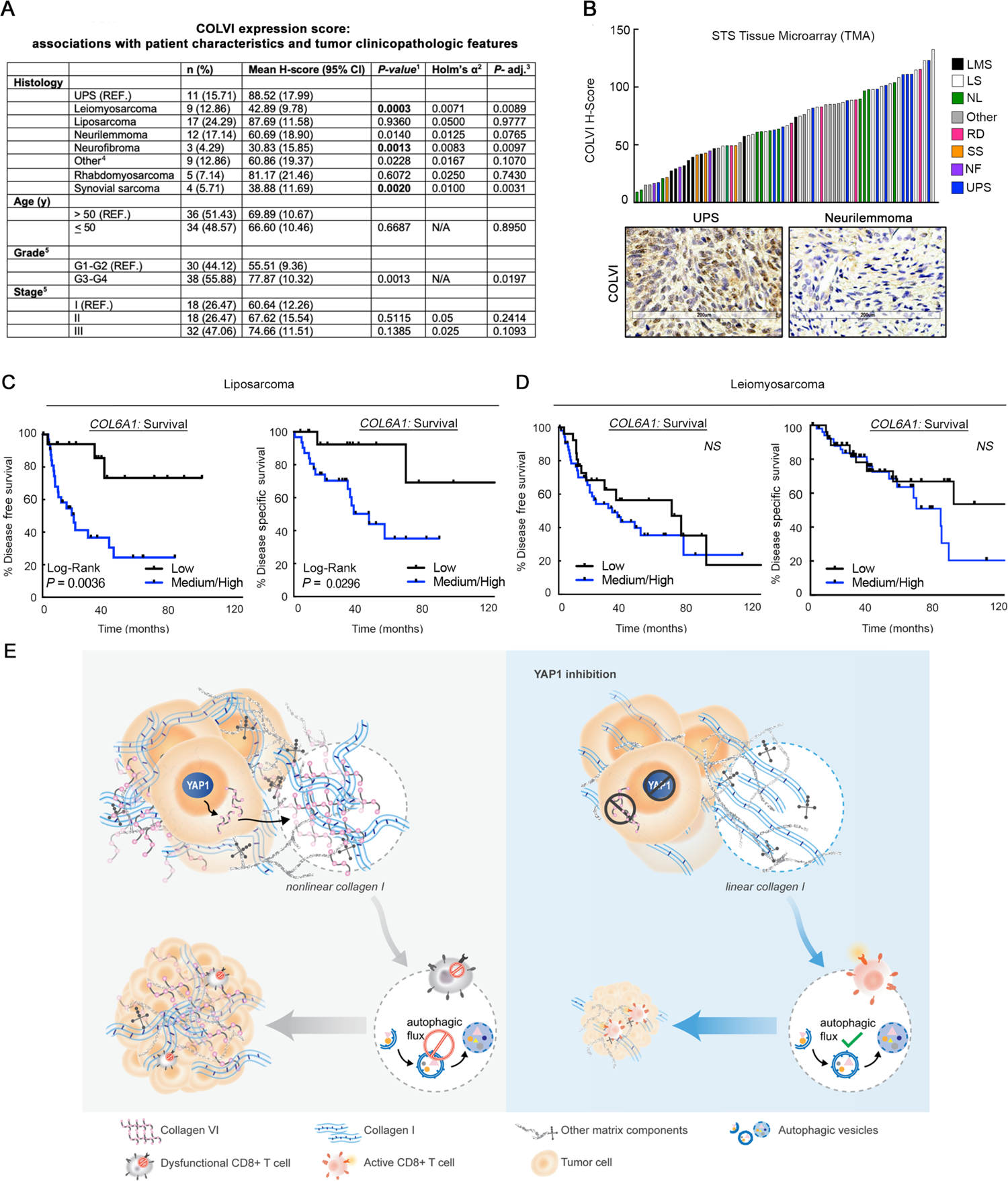
COLVI expression in the microenvironments of UPS and other soft-tissue sarcoma subtypes. (**A**) Association of IHC-based COLVI expression score with tumor subtype and clinicopathologic features. Sarcoma tissue microarray (TMA). ^1^Univariate linear models. ^2^In univariate analyses, the Holm-Bonferroni adjustment for multiple comparisons was performed for demographic or clinicopathologic variables with greater than two levels, with α = 0.05. Results are considered statistically significant (bold text) if the univariate *P-*value is smaller than the corresponding Holm’s alpha. ^3^Fully adjusted model (age, grade, stage, and histology). Correction for multiple comparisons was not performed due to insufficient statistical power. ^4^Includes 2 alveolar soft part sarcomas, 1 epithelioid hemangioendothelioma, 1 fibroma, 1 glomus tumor, 1 hemangioendothelial sarcoma, 1 hemangiopericytoma, 1 osteosarcoma, and 1 tenosynovial giant cell tumor. ^5^Excludes two benign cases. (**B**) Waterfall plot depicting IHC-based COLVI expression scores in individual tumors from **A**. LMS = leiomyosarcoma; LS = liposarcoma, NL = neurilemmoma, RD = rhabdomyosarcoma, SS = synovial sarcoma, NF = neurofibroma. “Other” as described in **A**. Representative images of UPS and neurilemmoma are also shown. (**C, D**) Kaplan-Meier disease-free and disease-specific survival curves of (**C**) liposarcoma and (**D**) leiomyosarcoma patients in TCGA-Sarcoma stratified by *COL6A1* gene expression. Each tertile (low, medium, high) represents 1/3 of patients. Log-rank test. (**E**) Model depicting study findings.

## DISCUSSION

Until now, our understanding of the role of the ECM in anti-tumor immunity was primarily limited to ECM-mediated T cell and macrophage migration. Additionally, upstream mediators of aberrant ECM protein composition in the TME were poorly defined. Herein, we establish a more specific and mechanistic understanding of individual collagen molecules in the ECM and how they impact adaptive immune cell function in solid tumors (**Figure 8E**). We discovered that the highly expressed transcriptional co-regulator Yap1 promotes the deposition of a pro-tumor matrix protein, ColVI, in the UPS ECM. In turn, ColVI opposes anti-neoplastic ColI molecules in the TME, altering their organization/architecture, thereby disrupting CD8^+^ T cell autophagic flux. Ultimately, this cascade facilitates CD8^+^ T cell dysfunction as measured by upregulation of co-inhibitory receptors, suppressed proliferation, and reduced effector function. Here, we report a novel, non-canonical role of Yap1 in the TME and establish a direct mechanistic link between specific ECM constituents and modulation of immune cell function.

Despite the incredible diversity of the collagen superfamily and the abundance of collagen molecules in solid tumors (21, 28), the effects of specific collagens and other matrix proteins on T cell effector function, differentiation, and anti-tumor efficacy are only beginning to be characterized. For example, using murine lung cancer models, Peng et al. (75) demonstrated that extracellular collagen molecules induced CD8^+^ T cell exhaustion and attenuated responses to α-Pd1 checkpoint therapy. Although these phenotypes were reversible following inhibition of the collagen cross-linking enzyme LOXL2, the authors did not attribute them to a specific collagen type, in part because LOXL2 inhibition disrupts the synthesis and deposition of multiple collagen species. Additionally, Liu and colleagues have implicated laminin-111 as an inhibitor of CD8^+^ T cell expansion and function in vitro; however, they did not pursue in vivo validation of these results (84). The authors did show that Matrigel, the primary component of which is laminin-111, may accelerate syngeneic mammary tumor growth in immunocompetent mice, but did not address whether Matrigel also directly stimulates cancer cell proliferation in vivo (84). Furthermore, Robertson et al. (85) recently showed that incubation of splenocytes with mammary carcinoma cells on collagen type IV (COLIV)-containing matrices may reduce T cell-mediated cancer cell clearance. However, this study (85) did not establish a direct mechanistic link between COLIV and suppression of T cell function, instead suggesting that COLIV may induce a more immunosuppressive transcriptional/secretory profile in mammary carcinoma cells. Moreover, the authors’ use of mammary carcinoma cells and unstimulated splenocytes from mice of different genetic backgrounds, as well as their reliance on transcriptional profiles from mixed carcinoma cell-T cell co-cultures as readouts of T cell function, make it challenging to interpret their results (85). Conversely, in the present study, we uncovered specific immunomodulatory roles of two distinct collagen species in UPS, and directly linked aberrant ECM composition/organization to induction of CD8^+^ T cell dysfunction. Using multiple orthogonal in vitro functional assays and in vivo readouts, we discovered that ColVI and ColI possess opposing roles in this tumor context, promoting and opposing immune evasion, respectively. Whereas a direct immunosuppressive role for ColVI has not been previously documented, the ColVI-mediated dysfunction program observed herein upregulated multiple T cell inhibitory receptors and dysfunction markers (Pd1, Tim-3, Lag3, CD39, Tox), suppressed CD8^+^ T cell proliferation, and blunted CD8^+^ T cell cytolytic capacity. In contrast, ColI was a tumor suppressor in vivo and reduced CD8^+^ T cell dysfunction relative to ColVI. These observations are consistent with recent studies demonstrating the stimulatory effects of COLI on CD8^+^ T cell function. For example, COLI co-stimulation enhanced peripheral blood-derived effector T cell expansion in vitro (74), and increased intratumoral T cell content and activation gene expression in pancreatic cancer models in vivo (25). In contrast, another study of three-dimensional culture models reported that high COLI density suppressed T cell proliferation and cytolytic marker gene expression, thereby impairing their ability to lyse melanoma cells (86). However, many of the experiments in (86) were performed with mixed populations of CD4^+^ and CD8^+^ T cells and are challenging to interpret. Nevertheless, taken together, these studies suggest that the effects of ColI on CD8^+^ T cells are complex and potentially context-specific. However, our work herein clearly shows that ColI is a requisite factor for CD8^+^ T cell function in UPS. By extension, stromal depletion strategies seeking to reduce COLI deposition in the TME could elicit detrimental outcomes in sarcomas.

One of the most intriguing findings from our study is that ColVI in the UPS TME directly remodels extracellular ColI. We suspect that this ColVI-mediated matrix remodeling masks binding motifs on ColI, such as RGD (Arg-Gly-Asp) sites or GXXGER consensus sequences, that would otherwise facilitate tumoricidal ColI-CD8^+^ T cell interactions. However, the identity of the receptor on CD8^+^ T cells mediating interactions with ColI in the UPS TME remains an open question. We excluded the possibility that Lair1 may be involved given it was not expressed on CD8^+^ T cells in KP tumors. Similarly, the involvement of another ColI receptor, Ddr1, is unlikely, given Ddr1’s previously reported role in the negative regulation of CD8^+^ T cell migration/infiltration in carcinomas (87, 88). Conversely, certain ColI-binding integrins such as Itga1, Itgav, and Itgb1 may be candidates given their putative roles in/associations with promoting CD8^+^ T activity (89–92). However, these proteins are challenging to study in UPS because they are receptors for both ColI and ColVI (63–68). Therefore, careful biochemical studies will be required to fully elucidate the mechanism by which ColI promotes CD8^+^ T cell activity and inhibits immune evasion in UPS.

Senescence, functional exhaustion, insufficient homeostatic proliferation, deletion, and altered metabolism have all been proposed as largely T cell-intrinsic mechanisms that hamper endogenous and engineered T cell-mediated anti-tumor immunity (93). Our findings offer a novel alternative model in which cancer cell-intrinsic biology drives failure of cytotoxic T cell activity by indirectly interfering with T cell autophagic flux. Previous studies have shown that autophagy is rapidly induced upon T cell activation, and that the essential autophagy genes *Atg5* and *Atg7* are critical for the survival, activation, and expansion of mature T cells (94, 95). Moreover, disrupting T cell autophagic flux hinders clearance of damaged mitochondria, resulting in increased reactive oxygen species (ROS) generation and T cell apoptosis (95). Thus, whether and how aberrant ColVI deposition influences ROS production in T cells with dysregulated autophagic flux is an important direction for future research.

Our study has multiple implications for the clinical management of UPS in human patients. First, as UPS is a diagnosis of exclusion, some pleomorphic neoplasms are incorrectly classified as “UPS” when they are more likely to be pseudosarcomas or other high-grade sarcomas (96). Thus, our observation that COLVI levels are significantly increased in UPS relative to several other soft-tissue sarcomas indicates that it may be a useful diagnostic tool for distinguishing UPS from other dedifferentiated pleomorphic tumors. Second, our study revealed that Pd1 blockade extended survival of KPY, but not KP mice. This result indicates that anti-Pd1 treatment was not sufficient to reinvigorate dysfunctional effector T cells in KP mice, but did preserve CD8^+^ T cells with robust cytolytic function in the context of Yap1 deficiency. Taken together, our findings show that Yap1-mediated signaling can contribute to immune evasion by modulating the composition and organization of the TME, and indicate that individual collagen species may have unique or opposing effects on UPS patient responses to T cell-based therapies. Specifically, COLVI in the UPS ECM may be detrimental to the efficacy of anti-PD1 or anti-CTLA4 therapy, whereas COLI may potentiate responses to immune checkpoint inhibition. As a result, this study underscores the critical need to systematically evaluate the roles of individual ECM components in the regulation of immune cell function. Furthermore, our data specifically implicate YAP1 and/or COLVI targeting as promising strategies by which to improve the efficacy of checkpoint blockade and other T cell-based therapies in UPS, and potentially other desmoplastic solid tumors.

## METHODS

Detailed methods are provided in the Supplementary Methods file.

### Statistics

Analyses were performed using Prism (Graph Pad Software). Data are shown as mean ± SEM unless otherwise specified. Data were reported as biological replicates as indicated in the figure legends; in vitro experiments were replicated at least three times unless otherwise specified. Unpaired two-tailed t-tests and one-way ANOVAs were performed to determine if differences between two or three group means, respectively, were statistically significant. 2-way repeated-measures ANOVA, mixed models, or non-linear regression were used for in vivo tumor growth curves. For correlations, Spearman’s coefficient was used if at least one dataset was not normally distributed. Pearson’s coefficient was used if both datasets were normally distributed. Shapiro-Wilk test was used to assess normality. *P*-values of <0.05 were considered statistically significant.

### Study approval

All experiments were performed in accordance with NIH guidelines and approved by the University of Pennsylvania Institutional Animal Care and Use Committee (approval number 805758). Studies performed with human specimens were not considered human-subjects research because all samples were de-identified and not collected exclusively for the purposes of this research.

## AUTHOR CONTRIBUTIONS

Conceptualization: TSKEM, SG, JAF, MH

Methodology: HCP, AMF, YL, JAF

Validation: HCP, AMF, YL, VIN, HS, AD, RK, GEC, EFW, IM, NS

Formal Analysis: HCP, AMF, YL, RK, SD, GEC

Investigation: HCP, AMF, YL, VIN, HP, HS, AD, RK, SD, GEC, EFW, IM, DN, NS

Data Curation: AMF, YL, MG

Provision of resources: HH, DZ, JT, AW, KW, MH, JAF, SG, TSKEM

Writing-original draft preparation: AMF, HCP, TSKEM

Writing-review and editing: AMF, HCP, TSKEM

Visualization: HCP, AMF, YL, EFW, JAF, TSKEM

Supervision: MH, JAF, SG, TSKEM

Project administration: TSKEM

Funding acquisition: SG, TSKEM

## ACKNOWLEDGEMENTS

We acknowledge Rebecca Gladdy, MD, for STS-109 sarcoma cells, and Steven Leppla, PhD, for *Cmg2^-/-^*cells. We also thank James Hayden and Frederick Keeney of the Wistar Institute Imaging Facility for assistance with multiphoton microscopy and analysis; Gordon Ruthel, PhD, of the PennVet Imaging Core for assistance with SHG imaging of collagen hydrogels; and John Tobias, Ph.D., of the UPenn Molecular Profiling Facility for bioinformatic assistance. We further acknowledge the UPenn Human Immunology Core for procuring human CD8^+^ T cells. Lastly, we thank Martha Jordan, PhD, and Sydney Drury for their assistance with in vivo flow cytometry, and Linnea T. Olsson, MSPH, for assistance with molecular epidemiologic analysis of human sarcoma tumors. We apologize to those we were unable to cite due to space constraints. **Funding:** This work was funded by The University of Pennsylvania Abramson Cancer Center, The Penn Sarcoma Program, Steps to Cure Sarcoma, R01CA229688, The Johns Hopkins Physical Sciences–Oncology Center (U54CA210173), T32HL007971, DoD RA200237, and the American Cancer Society – Roaring Fork Valley Research Circle Postdoctoral Fellowship (PF-21-111-01-MM). The UPenn Molecular Pathology and Imaging Core, which provided routine histology services, is supported by P30DK050306. The Penn Cytomics and Cell Sorting Shared Resource Laboratory is supported in part by an NCI Grant (P30016520) to the Abramson Cancer Center. Use of the Leica SP8 MP at the PennVet Imaging Core was made possible by NIH grant S10OD021633-01. The UPenn Human Immunology Core is supported in part by P30AI045008 and P30CA016520.

