## Supplementary materials for "Oncogene-induced matrix reorganization controls CD8^+^ T cell function in the soft-tissue sarcoma microenvironment"

**Supplemental materials:**

- Supplementary methods and associated references
- Histopathology appendix and associated references
- Supplementary Figures 1-11

**SUPPLEMENTARY METHODS**

**Cell lines**

HEK-293T cells were purchased from ATCC (Manassas, VA, USA). STS-109 cells were derived from a human UPS tumor by Rebecca Gladdy, M.D. (University of Toronto). KP230 cells were derived from a UPS mouse tumor as described previously by David Kirsch M.D., Ph.D. (Duke University). STR analysis was performed at the time of derivation and confirmed in April 2015. Cells were purchased, thawed, and expanded; multiple aliquots were frozen down within 10 days of initial resuscitation. For experimental use, aliquots were resuscitated and cultured for up to 20 passages (4-6 weeks) before being discarded. Human PDAC-CAF cells were isolated and cultured as previously described (1). STS109 cells were cultured in DMEM with 20% FBS, 1% L-glutamine, and 1% penicillin/streptomycin (P/S). Other cells were cultured in DMEM with 10% FBS, 1% L-glutamine, and 1% P/S. All cells were negative for mycoplasma contamination (testing performed every 6-12 months).

**In vitro Drug Treatments and Reagents**

Cells were treated with SAHA (2 μM) (Sigma Aldrich) and JQ1 (0.5 μM) (gift from Jun Qi, Ph.D., Harvard University) or vehicle control for 48 hours.

**Lentiviral Transduction**

Glycerol stocks for shRNA-mediated knockdown of *Yap1* TRCN: 0000095864, 0000095866, 0000095867, 0000095868; *YAP1* TRCN: 0000107266, 0000107267; *Col6a1* TRCN: 0000091533, 0000091535, 0000091536; *COL6A1* TRCN: 0000116957, 0000116961; *Col1a1* TRCN: TRCN0000090505, TRCN0000090507; and scramble shRNA were obtained from Sigma Aldrich or Dharmacon. shRNA plasmids were packaged using the third-generation lenti-vector system (pMDLg/pRRE, pRSV-Rev, and pMD2.G/VSVG) and expressed in HEK-293T cells. Supernatant was collected 24 and 48 hours after transfection and concentrated using 10-kDa Amicon Ultra-15 centrifugal filter units (Millipore) or polyethylene glycol-8000 precipitation. Target cell transduction was performed in the presence of 7.5–8.0 μg/mL polybrene (Sigma-Aldrich, #H9268). Puromycin selection (1-3 μg/mL) was performed 48 hours after infection. Cells were used in experiments 48 hours after selection. For experiments with knockdown of both *Col6a1* and *Col1a1* in the same cell population, transduction was done sequentially (*Col6a1* followed by *Col1a1*) because both constructs contain the same selection marker.

**Immunoblots**

Protein whole-cell lysates were prepared in SDS/Tris (pH 7.6) or RIPA lysis buffer, separated by SDS-PAGE gel electrophoresis, transferred to nitrocellulose or PVDF membranes, blocked according to standard protocols, and probed with the following antibodies: rabbit anti-YAP1 (Cell Signaling #4912; 1:1,000), rabbit anti-COL6A1 (Proteintech #17023-1-AP; 1:500–1:1,000), rabbit anti-COL6A2 (NOVUS #NBP1-90951; 1:500), rabbit anti-GAPDH (Cell Signaling #2118; 1:5000-1:10,000), rabbit anti-LC3B (R&D MAB85582 1:5000), and rabbit anti-collagen I (Boster Bio #PA2140-2, 1:500). HRP-linked anti-rabbit secondary antibodies (Cell Signaling #7074;

1:2500) were used for chemiluminescent detection. Immunoblotting for Col1a1/Col1a2 in cell culture-conditioned medium (CM) was performed as described in (2). For immuno-dot blots of CM, cells were cultured in normal growth medium for 24 hours, then switched to phenol red- and serum-free medium for 24 hours. CM was then concentrated in Amicon filter concentrators, spotted directly onto nitrocellulose membranes, and probed with antibodies as indicated above.

**Cell proliferation assay**

1 x $10^3$ cells per well were plated in 96-well plates and growth was monitored using the PrestoBlue Cell viability kit (Invitrogen #A13261).

**RNA isolation and qRT-PCR**

Total RNA was isolated from cells using the RNeasy Mini Kit (Qiagen) and from tissue using Trizol reagent (Life Technologies; 10 mg samples cut over dry ice). Reverse transcription of mRNA was performed using the High-Capacity RNA-to-cDNA Kit (Life Technologies). qRT-PCR was performed using a ViiA7 real-time PCR instrument (Applied Biosystems). All probes were TaqMan "best coverage" probes (Life Technologies). *HPRT* and/or *SDHA* were used as endogenous controls.

**Gene expression microarray and ChIP-seq analyses**

The publicly available KP/KPY tumor microarray and human UPS ChIP-seq datasets were first reported in (3). The gene expression microarray of control and SAHA/JQ1-treated KP cells can be found in NCBI GEO under the accession number GSE109923. The gene expression microarray dataset capturing 5 independent KP and KPY bulk tumors can be found under the GEO accession number GSE109920. The ChIP-seq dataset of H3K27 acetylation can be found under the GEO accession number GSE97295. The publicly available Yap1-driven embryonal rhabdomyosarcoma (eRMS) dataset was first reported by Tremblay et al. (4) and can be found in GEO under accession number GSE55186. The single-cell RNA sequencing dataset of $CD45^+$ cells in KP UPS tumors is from (5) and can be accessed in GEO under GSE144507.

***In vivo* tumor models**

GEMMs: LSL-*Kras*$^{G12D+}$; *Trp53*$^{fl/fl}$; *YAP1*$^{fl/fl}$ (KPY) mice were generated by crossing LSL-*Kras*$^{G12D+}$; *Trp53*$^{fl/fl}$ (KP) with *YAP1*$^{fl/fl}$ animals as previously described (3). We generated tumors in KP and KPY mice via injection of a calcium phosphate precipitate of adenovirus-expressing Cre recombinase (University of Iowa) into the right gastrocnemius muscle of 3-month-old male and female mice (3). Male and female GEMMs were examined in our study. Similar findings were observed in both sexes so results are reported for both sexes together.

Syngeneic transplant models: Cultured KP cells (SKPY42.1 cells on a pure C57BL/6J background, a gift from Sandra Ryeom, Ph.D., University of Pennsylvania) were detached using 0.25% trypsin (GIBCO), washed with 1x PBS, and counted in preparation for implantation. Cells were implanted subcutaneously (s.c.) or orthotopically (intramuscular, i.m.) into recipient syngeneic C57BL/6J or nu/nu mice (The Jackson Laboratory; strain codes:

000664 and 002019, respectively); specific cell numbers injected in each experiment are noted in the text and figure legends. To maintain tumorigenicity in immunocompetent mice due to genetic drift issues, SKPY42.1 cell lines were re-derived in March 2022: 2 x $10^6$ cells were injected s.c. into recipient syngeneic C57BL/6J mice and allowed to form tumors for 5 days. Tumors were excised, minced, and digested with 2.5% trypsin for 20 minutes at 37°C with vortexing every 5 minutes. Cell suspensions were then quenched with sarcoma cell culture medium (DMEM + 10% FBS + 1% L-glutamine + 1% P/S), filtered through 40-μm strainers, washed with PBS, and plated in tissue-culture plates. Re-derived cells were passaged 10 times before validation and use in experiments. Validation consisted of genotyping for recombined *Kras*$^{G12D}$ and *Trp53*$^{fl}$ alleles; assessment of *Yap1*, *Col6a1*, and *Col1a1* gene expression levels; and observation of shScr and shCol6a1 tumor growth kinetics in C57BL/6J mice. The re-derived cell line that most closely resembled parental SKPY42.1 cells was used in further experiments. For all studies, SKPY42.1 cells (parental and re-derived) were not used beyond passage 20.

Tumor growth measurements and survival analyses: Tumor dimensions were measured using calipers every 2-3 days, and volumes were calculated using the formula $(ab^2)\pi/6$, where a is the longest measurement and b is the shortest (3). Tumor volumes of 1500mm$^3$ were used as endpoints for survival analyses.

### ***In vivo* checkpoint studies**

In checkpoint studies, 200 μg of α-mouse Pd-1 monoclonal blocking antibody (clone RMP1-14, BioXCell) or isotype control antibody (clone 2A3, BioXCell) was administered i.p. every 3 days for 9 days (3 doses total) once tumors became palpable. In SAHA/JQ1 studies, autochthonous KP mice were treated as in (3). JQ1 was provided by Jun Qi (Dana-Farber Cancer Institute) and SAHA was purchased from Cayman Chemical. Vehicles (HP-β-CD and PEG400) were obtained from Sigma-Aldrich. For *in vivo* depletion of CD8$^+$ T cells, 200 μg of α-mouse CD8 (clone 2.43, BioXCell) or isotype control (clone LTF-2, BioXCell) antibody was administered intraperitoneally (i.p.) beginning three days prior to tumor implantation and repeated every three days until mouse sacrifice. CD8$^+$ depletion was confirmed in peripheral blood.

### **Immunohistochemistry**

Paraffin embedded tissue (5-μm sections) was stained using the BOND-RXm automated IHC/ISH stainer (Leica). The sarcoma tissue microarray was purchased from US Biomax (SO801a). Stained slides were digitally scanned by the Pathology Core Laboratory at the Children's Hospital of Philadelphia or the University of Pennsylvania School of Veterinary Medicine Comparative Pathology Core. Aperio ImageScope (Leica Biosystems) software was used for slide quantification. In murine tissue, CD4, CD8, Foxp3, and B220 positivity was assessed with the nuclear v9 algorithm; the positive pixel count algorithm was used to quantify F4/80 staining; and the Color Deconvolution algorithm was used to quantify ColVI and ColI staining. Digital histopathologic analysis of YAP1 and COLVI staining in human tissue is described in detail in the Immunohistochemistry Appendix. The following antibodies and dilutions were used: rat anti-Foxp3 (eBioscience #17-5773-82, 1:100), rat anti-F4/80 (Invitrogen #MF48000, 1:25), rabbit anti-CD4 (Cell Signaling #25229, 1:100), rabbit anti-CD8 (Cell Signaling #98941, 1:100), rat anti-B220 (BD Pharmingen #557390, 1:100), rabbit anti-collagen VI (Abcam #6588, 1:100 [human]), rabbit anti-collagen VI (abcam #182744, 1:2000 [mouse]), rabbit anti-YAP1 (Cell Signaling #14074, 1:100), anti-collagen I (Boster Bio #PA2140-2, 5 ug/mL), anti-Ki67 (Cell Signaling #12202, 1:500), anti-caspase 3 (Cell Signaling #9662, 1:500), and anti-cleaved caspase-3 (Cell Signaling #9661, 1:1500).

**Two photon second-harmonic generation (SHG) imaging and analysis of tumors**

Orientation and distribution of collagen fibers in tumors were visualized with SHG using a Leica TCS SP8MP 2-photon microscope (Leica Microsystems, Buffalo Grove, Il) equipped with a Chameleon femtosecond-pulsed laser (Coherent, Inc, Santa Clara, CA.). 960nm excitation was used to create better signal separation from endogenous autofluorescence. SHG signal was collected with a HyD detector through a DAPI filter cube, while autofluorescent signals were collected through FITC and TRITC filter cubes with standard PMTs and used to determine background separation. Freshly excised tumors, maintained in PBS, were cut in half and imaged to a depth of 100 μm in random areas across the cut surface using a 25X 1.0NA water immersion objective. Selected 5 μm stacks with even illumination, derived from the original, were used for quantification. Alternatively, tumor tissues were fixed for 48 hours in 4% neutral buffered paraformaldehyde, embedded in paraffin, sectioned at 7-10 μm, collected on glass slides and allowed to set. The unstained and unmounted tissues were then imaged as described above. 5-7 μm stacks were acquired in 1 μm steps at each location and 5 μm subset stacks were further processed for analysis. Each image stack was deconvolved with Huygens software (Scientific Volume Imaging, Hilversum, The Netherlands) and quantified with Leica LASX software using a standardized macro algorithm to determine total area of collagen within each measured area. Collagen fiber width and linearity from SHG images were analyzed using CT-FIRE, a previously established method for analysis of SHG images of tumors reported in (6). The software segments the image and calculates intensity gradients within subregions of images and uses them to track the overall directions of fibers. Linearity of fibers was calculated by overall displacement between the start and end of the fiber divided by the total length of the fiber.

**Flow cytometry of murine tumor samples**

GEMM tumors: Tumors were harvested and minced, and single cell suspensions were generated by digestion with 2 mg/mL collagenase B (Roche #11088823103) and 30 U/ml DNase I for 45 minutes at 37°C. RBCs were lysed using ACK lysis buffer. Samples were filtered through 70-μM cell strainers and incubated for 5 minutes at room temperature with anti-mouse CD16/32 Fc Block, and subsequently stained on ice with fluorophore-conjugated primary antibodies for identification of cell populations by flow cytometry. 7AAD (BioLegend) was used for dead cell discrimination. Flow cytometry was performed on an LSR II Flow Cytometer (BD Biosciences) and analyzed using FlowJo software (BD Biosciences).

T cell dysfunction panel (syngeneic transplant): Single-cell suspensions from bulk tumors were generated by digestion with 2 mg/mL collagenase B and 0.01 mg/mL DNase I. RBC lysis was performed as indicated above, after which samples were filtered through 40-μM cell strainers and incubated with anti-mouse CD16/32 Fc block (BioLegend #101320) for 10 minutes on ice (1 uL FC block/1 x $10^6$ cells). T cell enrichment was then performed using the mouse CD3ε microbead kit with LD columns (Miltenyi Biotech). T cell-enriched cell populations were then eluted from the columns and stained with LIVE/DEAD Fixable Aqua Dead Cell Dye (ThermoFisher) or eBiosience Fixable Viability Dye eFluor 455UV (ThermoFisher, 1:1000) for dead cell discrimination. Subsequently, extracellular staining was performed with pre-titrated fluorescent-conjugated antibodies on ice for 30 minutes, followed by fixation and permeabilization (eBioscience Foxp3/Transcription Factor Staining Buffer Set) and intracellular staining (1 hour at room temperature). Data were acquired with a FACSymphony A3 Lite cytometer (BD) and analyzed with FlowJo software. Single-color compensation controls were generated with UltraComp eBeads Plus Compensation beads or murine splenocytes, and positive and negative populations were identified using "fluorescence-minus-one" controls. The following antibodies were used: CD45 BUV563 (clone 30-F11, BD Biosciences, 1:200), CD3e BV650 (clone 145-2C11, BD Biosciences, 1:100), CD4 BUV395 (clone GK1.5, BD Biosciences, 1:200), CD8a AlexaFluor 700 (clone 53-6.7, BioLegend, 1:100), CD44 APC-Cy7 (clone IM7, BioLegend, 1:100), CD62L PE-eFluor610 (clone MEL-14, ThermoFisher, 1:100), PD1 BUV737 (clone RMP1-30, BD Biosciences, 1:100), Tim-3 SB436 (clone RMT3-23, ThermoFisher, 1:50), Lag3 SB600 (clone C9B7W, ThermoFisher, 1:25), CD69 BV711 (clone H1.2F3, BioLegend, 1:50), CD39 PE-Cy7 (clone Duha59, BioLegend, 1:100), B220 FITC (clone RA3-6B2, BioLegend, 1:200), LY6G FITC (clone 1A8, BioLegend, 1:100), CD11b FITC (clone M1/70, BioLegend, 1:100), Tox APC (clone REA473, Miltenyi, 1:100), TCF1 PE (clone S33-966, BD Biosciences, 1:100), and GZMB PerCP-Cy5.5 (clone QA16A02, BioLegend, 1:50).

### T cell isolation and activation

Murine T cells: $CD8^+$ T cells were isolated from spleens of 6-12-week-old C57BL/6J mice. Mice were maintained per guidelines approved by the Johns Hopkins University or University of Pennsylvania Animal Care and Use Committee. Cells were macerated through cell filters using a syringe and treated with ACK lysis buffer for erythrocyte lysis. $CD8^+$ T lymphocytes were isolated from splenocytes by negative selection using $CD8^+$ isolation kits (130-104-075) and magnetic LS columns from Miltenyi Biotech according to the manufacturer's protocol. Cells were activated by culture in RPMI with 10% FBS, 1% P/S, and IL-2 (40 U/mL) for 4 days with mouse T cell stimulator (CD3/CD28) Dynabeads (ThermoFisher 11453D) prior to use in experiments.

Human T cells: Human $CD8^+$ T cells were purchased from Astarte Bioscience or obtained from the University of Pennsylvania Human Immunology Core and cultured in RPMI with 10% FBS, 100 U/mL IL-2, and 1% penicillin/streptomycin. T cells were activated for 4 days using human T cell stimulator (CD3/CD28) Dynabeads (ThermoFisher 11161D) prior to use in experiments.

### Generation of decellularized ECMs (dECMs)

Glass-bottom chamber slides, tissue culture plates, or polymer-coated chamber slides (ibidi, Fitchburg, WI) were coated with 0.2% gelatin for 1h at 37°C, followed by fixation with 1.0% glutaraldehyde and 1 mM ethanolamine at room temperature for 30 minutes each. Plates were washed 2-3 times with PBS after each step, and cells were seeded such that they would reach 90% confluency the following day. Subsequently, culture media was supplemented with 50 $\mu$g/mL L-ascorbic acid (Sigma a7506) and cells were maintained for 5-7 days under hypoxic conditions (1% oxygen) in a CellXpert hypoxia incubator (Eppendorf). Culture media containing L-ascorbic acid and puromycin (to maintain selection of shRNA-infected cells) was changed daily. The decellularization protocol is adapted from Harris et. al (7). Briefly, cells were washed twice with wash buffer 1 as previously described prior to lysis with NP-40 based lysis buffer. Cell debris was removed with wash buffer 2 (three washes) prior to four final washes with DI water. ECMs were stored at 4°C in PBS for up to 2 weeks until use in experiments.

### Fabrication of collagen hydrogels

Collagen gels were prepared as reported previously in (8) with the following modifications: Rat tail collagen type I (Corning 354249) was mixed with native human collagen type VI (Rockland 009-001-108), native human collagen type IV (Rockland 009-001-106), or an equimolar amount of acetic acid, as well as 10x Medium 199 buffer (ThermoFisher). The solution was neutralized with 1M NaOH dropwise and mixed with 1x M199 buffer or RPMI culture medium (ThermoFisher). The final concentration of collagen I was 1.5 mg/mL, whereas collagen VI and collagen IV were each used at 250 μg/mL. Gels were polymerized at 37°C for 30-45 minutes, washed with PBS, and incubated overnight with culture medium to help remove/equilibrate excess salts from the 10x M199 and residual sodium azide from the COLVI.

### Hydrogel stiffness, diffusion, and oxygen concentration assessments

Collagen hydrogels (COLI ± COLVI) were fabricated as described above. Bulk stiffness was measured using an HR-20 Discovery Hybrid Rheometer (TA Instruments) equipped with an 8-mm cross-hatched parallel plate at 37°C. Storage modulus G' and loss modulus G'' were monitored at a 1%-20% strain range and fixed frequency of 0.1 Hz. All hydrogels were prepared as discs measuring 8 mm in diameter. Rhodamine B diffusion assays in COLI ± COLVI hydrogels were performed as described in (9). Non-invasive measurements of hydrogel oxygen concentrations were obtained as in (8).

### Flow cytometry of isolated T cells

Murine T cells: Murine $CD8^+$ T cells were activated as described above and incubated with decellularized ECMs prior to flow cytometry analysis. To assess cell-surface dysfunction marker expression, cells were incubated in IL2-replete culture medium for 48 hours and stained with Zombie NIR viability dye (BioLegend, 1:2500) and the following antibodies: CD3 APC (clone 17A2, BioLegend, 1:100), CD8a PerCP (clone 53-6.7, BioLegend, 1:100), CD44 FITC (clone IM7, BioLegend, 1:50), CD62L APC-Cy7 (clone MEL-14, BD Biosciences, 1:50), TIM-3 PE-Cy7 (clone B8.2C12, BioLegend, 1:25), and PD-1 PE (clone RMP1.14 BioLegend, 1:20). Cells were gated on single cells, then T cell markers (CD3 vs. CD8), then activated effector cells (CD44+/CD62L-). Non-viable cells were not gated out because 1) many antigen-experienced effector T cells, including dysfunctional T cells, die via apoptosis or express cell death markers (10-12), and 2) IL2 deprivation is well known to cause extensive T cell apoptosis (13), and gating out large numbers of non-viable cells would substantially hinder our ability to robustly evaluate dysfunction phenotypes. To evaluate intracellular cytokine expression, cells were incubated on hydrogels in IL2-containing culture medium for 48 hours, and subsequently treated with Cell Stimulation Cocktail

+ Brefeldin A (eBioscience) for 4 hours. Cells were then retrieved and stained with Zombie NIR viability dye (1:2500); fixed using the Cyto-Fast Fix-Perm Buffer Set (BioLegend); and stained with the following intracellular antibodies: Ki67 AlexaFluor 488 (clone SolA15, ThermoFisher, 1:100), TNFα FITC (clone MP6-XT22, BioLegend, 1.5 uL/100 uL staining volume), and IFNγ PE-Cy7 (clone XMG1.2, BioLegend, 1:400). All staining and washes were performed in the presence of GolgiPlug (Protein Transport Inhibitor containing Brefeldin A; BD). Data were acquired on a BD FACS Canto II or FACSymphony A3 Lite and analyzed with FCS Express or FlowJo Software. Cells were gated on single cells (FSC-A versus FSC-H), then viable cells, then T cell markers (CD3 vs. CD8). Gating on positive populations was determined using "fluorescence-minus-one" controls.

Human T cells: Human $CD8^+$ T cells were activated as described above, incubated on hydrogels for 48 hours, and analyzed via flow cytometry. To assess cell-surface dysfunction marker expression, cells were incubated in IL2-replete culture medium and stained with the following antibodies: CD44 BB515 (clone G44-26, BD Biosciences, 1:20), CD62L APC (clone DREG-56, BioLegend, 1:20), PD1 PE-Cy7 (clone NAT105, BioLegend, 1:20), and TIM3 PE-Cy5 (clone F38-2E2, BioLegend, 1:20). Dead cells were not removed from the analysis as discussed in "Murine T cells" above. Intracellular cytokine expression was evaluated as described in "Murine T cells" above, except the following antibodies were used: Ki67 AlexaFluor 700 (clone B56, BD Biosciences), IFNγ Pacific Blue (clone B27, BioLegend, 1:100), and TNFα FITC (clone MAb11, BioLegend, 1:50).

Receptor blocking assays: Activated human $CD8^+$ T cells were incubated with anti-NG2 (clone 9.2.27, Santa Cruz Biotechnology, 40 μg/mL), anti-ITGB1 (clone P5D2, R&D Systems, 10 μg/mL), or the corresponding isotype control (IgG2a and IgG1, respectively) for 1 hour at 37°C prior to encapsulation in hydrogels. Alternatively, cells were treated with cilengitide (10 μg/mL, Tocris Bioscience) to block integrin αvβ3 or vehicle control. Cells were analyzed for flow cytometric expression of dysfunction markers and KI67 as described above 24 hours after encapsulation in hydrogels.

**Two photon SHG imaging and analysis of collagen hydrogels**

Collagen hydrogels were constructed as described above in 35-mm glass-bottom dishes with a 10-mm micro-well and #1.5 cover glass (Cellvis, Mountain View, CA). COLI fiber architecture (in COLI ± COLVI or COLI ± COLIV gels) was visualized with SHG using a Leica SP8 MP 2-photon microscope (Leica Microsystems) equipped with a Chameleon Vision II Ti:Sapphire laser (Coherent, Inc.). Forward and backward SHG signal was collected using an excitation wavelength of 910 nm; however, only backward signal was analyzed because robust forward signal was not detectable in all replicate experiments. Collagen hydrogel image stacks (15-19 images per stack at steps of 1 μM; 2-5 stacks per hydrogel) were acquired with a 20X 1.0 NA water-immersion lens with 2X optical zoom. The same instrument settings were used across replicate experiments. Max projection images of each stack were generated and analyzed with CT-FIRE.

**Immunofluorescence**

dECMs (single stains) and hydrogels: dECMs were fixed in 4% PFA, permeabilized, and blocked according to standard protocols. Staining was performed with rabbit anti-collagen VI antibodies (Abcam #ab6588, 1:100) overnight at 4°C, followed by goat anti-rabbit (H+L) AlexaFluor 546-conjugated secondary antibodies (ThermoFisher, 1:1000) for 30-60 minutes at room temperature. For hydrogels, mouse anti-p62 (R&D MAB8028 1:250) was incubated overnight at 4°C followed by goat anti-mouse (H+L) AlexaFluor 488-conjugated secondary antibodies (ThermoFisher, 1:500) for 1 hour at room temperature.

Non-decellularized ECMs: Analysis of ColI structure/organization in control and shCol6a1 ECMs was performed on matrices that did not undergo decellularization in order to preserve matrix structure to the highest possible degree. Matrices were fixed with 4% PFA and blocked with 3% BSA. To visualize extracellular ColI, matrix was stained with AlexaFluor 488-conjugated anti-collagen I antibodies (Abcam #ab309248) overnight at 4°C. No Triton X-100 was included at any point to minimize cell permeabilization and intracellular staining. Matrix was then mounted with Prolong Diamond and images were acquired on a Leica TCS SP8 confocal microscope. Images were analyzed with CT-FIRE.

dECM co-immunofluorescence: dECMs were fixed in 4% PFA and blocked in 3% BSA and 0.3% Triton X-100 in PBS. Antibodies were diluted in the same blocking buffer and all incubations were carried out at room temperature. Matrix was first incubated with anti-collagen VI antibody (Abcam ab6588, 1:200) for 3 hours, and then with donkey anti-rabbit monovalent F(ab) Rhodamine red secondary antibody (Jackson ImmunoResearch, 711-297-003, 1:50) for 2 hours. Next, matrix was incubated with anti-collagen I antibody for 2 hours (Boster Bio PA2140-2, 1:200), and then with AlexaFluor 488-conjugated goat anti-rabbit divalent F(ab) secondary antibody (ThermoFisher, A-11008, 1:1200) for 1 hour. Labeled matrix was then mounted with Prolong Diamond (Thermo Scientific, P36961). Images were acquired on a Leica TCS SP8 confocal microscope. Images and 3D reconstructions were analyzed with FIJI software, and 3D rotation movies were made with Leica LAS X 3D viewer. For ColVI-ColI co-localization (BIOP JaCoP plugin) and intensity analyses, intensity thresholds (500 and 800 for ColVI and ColI, respectively) were set based on evaluation of shCol6a1 and shCol1a1 controls.

Cytolysis arrays: To visualize extracellular COLVI and COLI expression in the setting of the cytolysis assay, STS-109 cells were seeded on cytolysis arrays, fixed in 4% PFA, and blocked in 10% normal goat serum in PBS. Cells were then incubated with antibodies diluted in 1% normal goat serum in PBS overnight at 4°C (anti-collagen VI: Abcam ab6588, 1:200; anti-collagen I: Boster Bio #PA2140-2, 1:200). Cells were then permeabilized and incubated with phalloidin-rhodamine red (Biotium #00027, 1:800) and Hoechst 33342 (NucBlue Live ReadyProbes Reagent, ThermoFisher #R37605) diluted in 0.3% Triton X-100 in PBS at room temperature for 30 min. The cell culture surface of the array was then removed and mounted onto a glass coverslip. Images were acquired on an Olympus IX81 wide-field microscope.

**Flow cytometric apoptosis assay**

STS-109 human UPS cells were infected with control, *YAP1*-, or *COL6A1*-targeting shRNAs and selected with puromycin as described above. Ninety-six hours after transduction, cells were trypsinized, plated on 6-cm plates, and allowed to recover overnight. Cells were then treated with vehicle control (0.1% BSA/PBS) or recombinant human TNF$\alpha$ (300-01A, PreproTech) or IFN$\gamma$ (300-02, PreproTech) at 10 ng/mL or 100 ng/mL for 48 hours (14-16). Subsequently, cells were harvested by trypsinization. Dead/floating cells that remained in cell culture supernatants prior to trypsinization were retained. Annexin V-APC/Fire 750 (5 μL/$10^6$ cells/100 μL staining volume; BioLegend 640953) and propidium iodide (4.5 μL/3 x $10^5$ cells/300 μL staining volume; R&D systems) staining were then performed sequentially; cells were incubated for 15 minutes at room temperature in the dark after the addition of each stain. Propidium iodide staining was performed in the presence of 0.4 mg/mL RNaseA (NEB T3018). Flow cytometric data (gated on single cells) was acquired on a BD LSRII instrument, with gating on positive populations guided by "fluorescence-minus-one" controls. Single-color controls contained a ~50/50 mixture of viable and heat-killed cells to ensure sufficient positive signal for instrument compensation. Data were analyzed with FlowJo Software.

**xCELLigence real-time target cell cytotoxicity analysis**

CART cell preparation: CART-TnMUC1 T cells (17) and donor-matched untransduced (NTD) cells were thawed in a 37°C water bath. T cells were resuspended in STS-109 culture medium and centrifuged twice for 5 minutes at 500 x g to remove residual cryopreservants. Cell pellets were then resuspended in culture media at a final concentration of 1 x $10^6$ cells/mL and rested overnight at 37°C in 5% $CO_2$.

xCELLigence real-time cell cytotoxicity assay: For the xCELLigence cytolysis assay, STS109 UPS cells (target cells) were harvested and seeded in triplicate into 96-well polyethylene terephthalate plates (E-Plate VIEW 96 PET) at 5,000 total targets per well. Cells were allowed to adhere for 24 hours (37°C, 5% $CO_2$). Adhesion of cells to the gold microelectrodes in the E-Plates impedes the flow of electric current between electrodes. This impedance value, plotted as a unitless parameter known as "Cell Index", increases as cells proliferate and then plateaus as cells approach 100% confluence. Effector cells (CART-TnMUC1cells or NTD T cells) were added to wells containing the appropriate targets at E:T cell ratios of 10:1, 5:1, 1:1, and 0:1. These non-adherent effector cells in suspension do not cause impedance changes in and of themselves (due to lack of adherence to the gold microelectrodes). In these experiments, target cells lysed by 1% Triton-X-100 were used as the positive control, while NTD cells at the same E:T ratios listed above were assayed as negative controls. Cytotoxic activity of the T lymphocytes was determined via continuous acquisition of impedance data for each well containing target cells over 6 days. Raw impedance data was analyzed using RTCA 2.1.0 software. Curves indicate the percent (%) Cytolysis using NTD controls as a reference. The curves also subtract the signal from wells containing effector cells alone and from wells containing target cells treated with a rapid killing agent (i.e., Triton-X-100) to compensate for residual background signal. The general equation for calculating the % Cytolysis for a sample at a time point *t* is: % $Cytolysis_{st}$= [1-($NCI_{st}$)/($AvgNCI_{Rt}$)]x100, where, $NCI_{st}$ is the Normalized Cell Index for the sample and $NCI_{Rt}$ is the average of Normalize Cell Index for the matching reference wells.

**Autophagy detection**

Murine or human $CD8^+$ T cells were incubated on hydrogels for 4 hours to capture the effects of COLVI on autophagy before the onset of overt dysfunction. The Cyto-ID Autophagy detection kit 2.0 (Enzo Life Sciences) was used to evaluate autophagic vacuoles according to the manufacturer's instructions with the following modifications: CytoID-containing media (1:500) was overlaid onto hydrogels containing activated CD8+ T cells for 30 minutes at 37C. Media was replaced and cells were incubated for 4 hours prior to fixation with 4% PFA. Cells were then stained with Phalloidin-AlexaFluor 546 (1:500) and DAPI (1:1000) to localize individual cells and imaged on a Zeiss LSM780 confocal at 40x magnification. Vacuole number and intensity was quantified using Imaris 8.3.1. p62 immunofluorescent staining was carried out as described above. For LC3B immunoblots, hydrogels containing $CD8^+$ T cells were lysed directly with 6x SDS loading buffer and boiled at 95C for 5 min. 50 uL of the result lysate were loaded onto SDS-PAGE gels for immunoblotting as described above.

**Human samples**

Human sarcoma or non-malignant muscle samples were obtained from surgically resected tumors from patients (de-identified) undergoing therapeutic surgical resection in accordance with protocols approved by the Institutional Review Board at the University of Pennsylvania.

**Oncomine, TCGA, and cBioPortal analyses**

We used the publicly available dataset published in Detwiller et al. through Oncomine Research Premium edition software (version 4.5, Life Technologies) to query ColVI gene expression in sarcoma and normal muscle tissue. Kaplan-Meier survival analyses using TCGA data were conducted using gene expression values ranked according to the rlog transformation. Kaplan-Meier analyses were performed for overall, disease-free, and disease-specific survival of patients. Gene expression correlations were conducting using TCGA gene expression data normalized with DESeq2. Clinical data were downloaded from the "Adult Soft Tissue Sarcomas (TCGA, Cell 2017)" (18) cBioPortal dataset on March 21, 2019. TCGA PanCancer RNA-seq data ("TCGA PanCancer Atlas Studies"; 10,967 samples across 32 studies) were obtained from cBioPortal.

**SUPPLEMENTAL METHODS: REFERENCES**

### APPENDIX: HISTOPATHOLOGIC ANALYSIS OF UNDIFFERENTIATED PLEOMORPHIC SARCOMA SURGICAL SPECIMENS

The goal of this analysis was to use IHC to determine the relationship between collagen VI (COLVI) protein expression and YAP1 activity in human undifferentiated pleomorphic sarcoma (UPS). We used nuclear YAP1 as a surrogate for YAP1 pathway activity in specimens from primary tumors surgically resected at the Hospital of the University of Pennsylvania (HUP). In this Appendix, we describe our digital histopathology approach in detail and provide validation for each method. This document can also be used as a guide for digital histopathologic analyses of histologically complex tumors.

#### *Slide annotation and quantification of COLVI protein expression*

Stained slides were scanned into digital images (20x magnification) using an Aperio VERSA 200 scanner (Leica Biosystems). Images were annotated in Aperio ImageScope to exclude areas with poor resolution or that contained scanning or staining artifacts (e.g., microtome blade marks, DAB deposits, and edge effects). Histologically normal cancer-adjacent tissue (e.g., adipose and skeletal muscle muscle), large deposits of erythrocytes, folded tissue, and debris were also excluded. Furthermore, as many of our biospecimens were sampled near the margins of surgically resected tumors, some slides only contained small foci of or diffusely arranged sarcoma cells. These cases were annotated conservatively in that only areas immediately surrounding regions of frank sarcoma cells were included (**Appendix Fig. 1A-B**). Annotations were reviewed by a pathologist (MH).

Like other extracellular matrix (ECM) proteins, COLVI is synthesized and biochemically modified inside the cell prior to secretion. Therefore, COLVI immunoreactivity is observed both intra- and extracellularly and was quantified with the Aperio ImageScope Positive Pixel Count algorithm (**Appendix Fig. 1C-D**). A “digital” COLVI H-score was calculated as follows. H-score was selected as a metric because it is widely used in the pathology literature and is both normalized to tissue area and weighted for staining intensity.

COLVI H-score = (3 × % strong-positive pixels) + (2 × % moderate-positive pixels*) + (1 × % weak-positive pixels)

*Note that Aperio uses the term “positive pixels” here; however, we believe this term is ambiguous and prefer to use “moderate-positive pixels” instead.

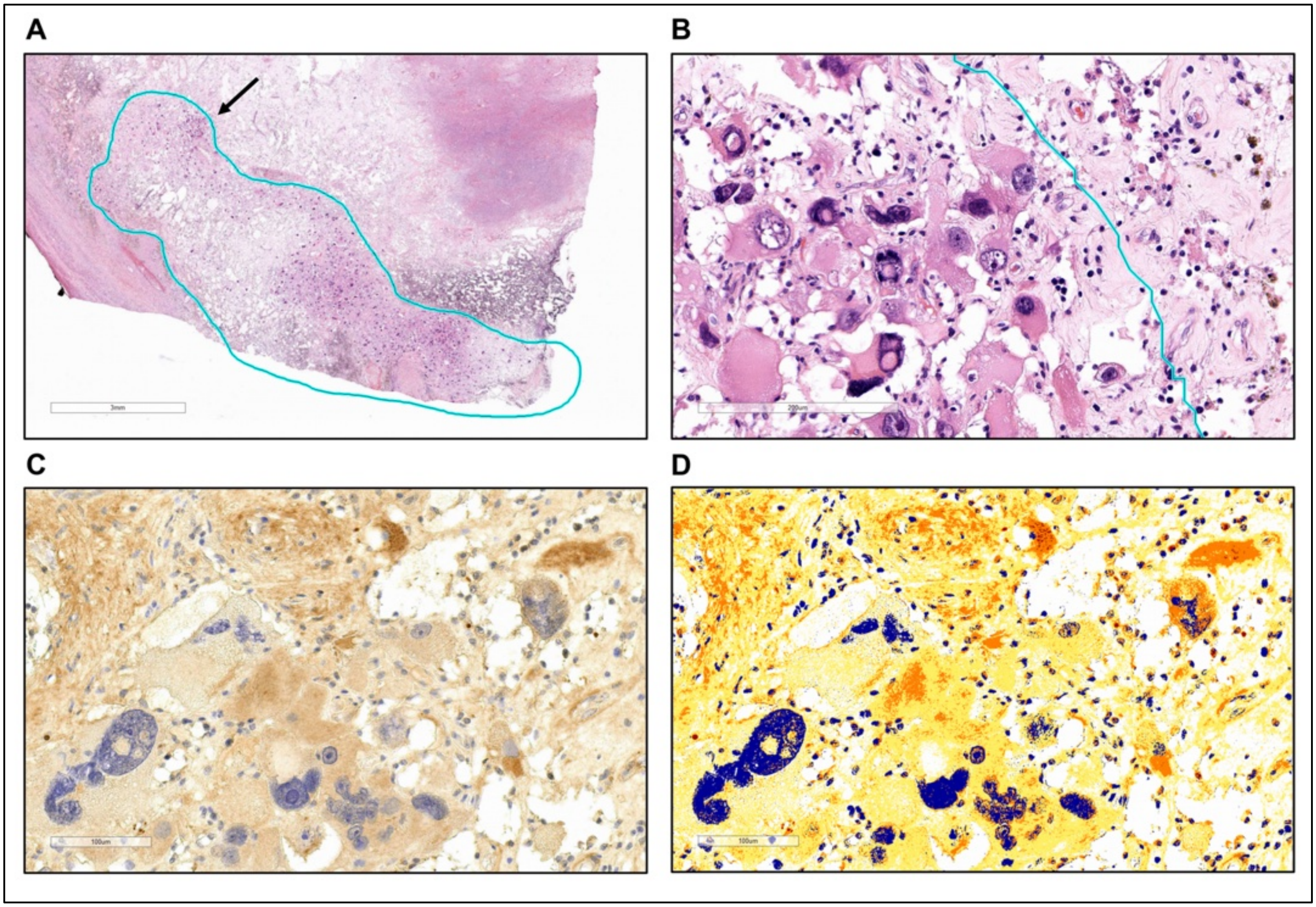


**Appendix Fig. 1. Slide annotations and quantification of COLVI IHC staining. A-B.** Example of a conservatively annotated slide where the biospecimen was obtained near the margins of the parent tumor. Only the area within the blue outline, which immediately surrounds frank sarcoma cells, was included in the analysis. Arrow indicates the approximate location of the high-magnification view shown in panel **B**. **C-D.** COLVI IHC staining (**C)** and the corresponding heat map from the Aperio Positive Pixel Count algorithm (**D**). Blue, yellow, orange, and red indicate negative, weak-positive, moderate-positive, and strong-positive staining, respectively.

***Analysis I: Quantification of nuclear YAP1 staining and correlation with COLVI protein expression***

In UPS, inactivation of the Hippo pathway facilitates nuclear localization of the transcriptional co-regulator YAP1. Upon nuclear translocation, YAP1 binds to TEAD-family proteins and activates pro-proliferative, pro-survival, and ECM-associated genes (19, 20). Therefore, nuclear YAP1 IHC staining was used as a surrogate for YAP1 pathway activation. Due to 1) the pleomorphic nature of UPS nuclei, 2) the strong expression of YAP1 in both the nucleus and cytoplasm of many tumor cells, and 3) the relatively weak hematoxylin staining intensity observed in our specimens, we used the Aperio Genie classifier tool and the Aperio Nuclear v9 algorithm to quantify nuclear YAP1 immunoreactivity.

We first created a Genie classifier that roughly partitioned YAP1-stained slides into the following three components: nuclei, cytoplasm and ECM, and slide glass (**Appendix Fig. 2A-D**). Extremely robust classification of “nuclei” and “cytoplasm and ECM” could not be achieved but was not explicitly required for this step. We then calculated total tissue area:

$$\text{Tissue area}_{\text{YAP1 Genie}} = \text{nuclei area}_{\text{YAP1 Genie}} + \text{cytoplasm and ECM area}_{\text{YAP1 Genie}}$$

The Nuclear v9 algorithm was then tuned to the YAP1-stained specimens and applied only to the Genie “nuclei” class on each slide. Use of Genie in this manner enabled more specific nuclear detection than did the same Nuclear v9 algorithm used in the absence of a classifier, particularly in areas where nuclear and cytoplasmic YAP1 staining intensity were similar (**Appendix Fig. 2E-F**). We then calculated a nuclear YAP1 H-score:

$$\text{Nuclear YAP1 H-score} = (3 \times \%\ 3^{+}\ \text{nuclear area}) + (2 \times \%\ 2^{+}\text{nuclear area}) + (1 \times \%\ 1^{+}\text{nuclear area})$$

Because YAP1 is only located inside the cell, its expression is dependent upon tissue composition; i.e., the percentage of tissue area that is occupied by cells (or nuclei, in the case of nuclear YAP1) vs. ECM and other tissue components. Therefore, we calculated the percent nuclear area of each slide. Note that we could not directly use the “nuclei area$_{\text{YAP1 Genie}}$” parameter from above because the rough segmentation of the Genie classifier would have greatly overestimated nuclear area (**Appendix Fig. 2C**).

$$\%\ \text{nuclear area}_{\text{YAP1 IHC}} = \frac{\text{Nuclear area}_{\text{YAP1 IHC}}}{\text{Tissue area}_{\text{YAP1 Genie}}} \times 100$$

where:

$$\text{Nuclear area}_{\text{YAP1 IHC}} = \text{Total nuclei}_{\text{YAP1 IHC}} \times \text{average nuclear size}_{\text{YAP1 IHC}}$$

Finally, we normalized the nuclear YAP1 H-score of each specimen to its respective percent nuclear area. This value was correlated with COLVI H-score for all available specimens (see **Figure 7J** in the manuscript).

$$\text{Normalized Nuclear YAP1 H-score} = \frac{\text{Nuclear YAP1 H-score}}{\text{\% nuclear area}_{\text{YAP1 IHC}}}$$

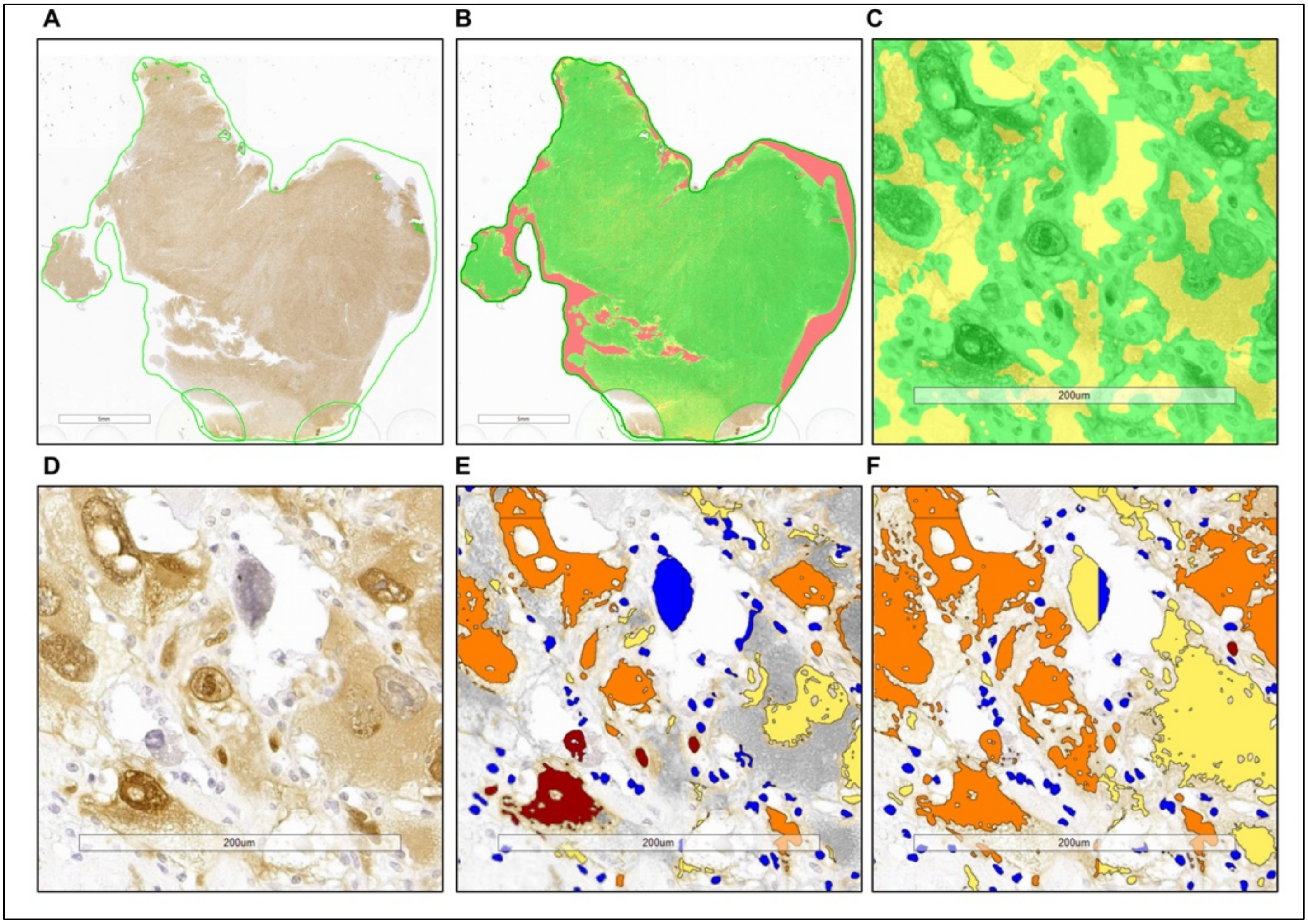


**Appendix Fig. 2. Quantification of nuclear YAP1 staining and correlation with COLVI protein expression. A-B.** Example of a YAP1-stained slide (**A**) roughly partitioned into nuclei (green), cytoplasm and ECM (yellow), and slide glass (pink) (**B**). **C.** High-power view of the YAP1 Genie classifier. **D.** IHC staining corresponding to the image shown in **C**. **E.** Nuclear v9 algorithm applied only to the "nuclei" class of the image shown in **C-D**. **F.** The same Nuclear v9 algorithm shown in **E**, applied to the entire slide rather than just the "nuclei" class of the image shown in **C-D**. Note that more cytoplasm and ECM are detected in panel **F** compared to panel **E**, particularly in cases where nuclear and cytoplasmic staining are similar. For panels **E** and **F**, blue, yellow, orange, and red indicate $0^+$ (negative), $1^+$, $2^+$, and $3^+$ staining, respectively.

***Analysis II: Validation of tissue composition metrics using H&E-stained serial sections***

One challenge of quantitative digital pathology approaches, particularly when performing slide annotations and algorithm tuning, is the requirement to strike a balance between tissue features that may yield false-positive (e.g., sectioning, staining, and scanning artifacts; debris) and false-negative (e.g., faint staining) results (21). In particular, we noticed that our Genie classifier and Nuclear v9 algorithms often omitted nuclei with faint hematoxylin staining, irrespective of whether they appeared positive or negative for YAP1. Because our supply of HUP tissue specimens is extremely limited, we were unable to validate our IHC results using an independent anti-YAP1 antibody (though we have independently validated the specificity of this antibody in other applications). However, we did endeavor to show that the tissue composition metrics obtained with the IHC-based Genie classifier algorithms were representative of our slide set. To this end, we used H&E-stained specimens from the same tissue blocks as the IHC-stained slides in accordance with published data showing that tissue morphometry metrics in a given specimen are highly correlated between replicate measurements (22, 23).

We first created an Aperio Genie classifier to roughly partition H&E-stained specimens into nuclei, cytoplasm and ECM, and slide glass (**Appendix Fig. 3A-B**). These measurements were used to calculate tissue area for each slide:

$$\text{Tissue area}_{\text{H\&E Genie}} = \text{nuclei area}_{\text{H\&E Genie}} + \text{cytoplasm and ECM area}_{\text{H\&E Genie}}$$

We then tuned the Nuclear v9 algorithm to the H&E-stained slides, applying it only to the "nuclei" class of each specimen. We observed that the more intense hematoxylin staining of these specimens facilitated improved nuclear detection compared to the IHC-stained slides (**Appendix Fig. 3C-E**). The assignment of nuclei to $0^+$ (negative), $1^+$, $2^+$, or $3^+$ staining categories was disregarded. The resulting data were used to calculate percent nuclear area for each slide.

$$\%\ \text{nuclear area}_{\text{H\&E}} = \frac{\text{Nuclear area}_{\text{H\&E}}}{\text{Tissue area}_{\text{H\&E Genie}}} \times 100$$

where:

$$\text{Nuclear area}_{\text{H\&E}} = \text{Total nuclei}_{\text{H\&E}} \times \text{average nuclear size}_{\text{H\&E}}$$

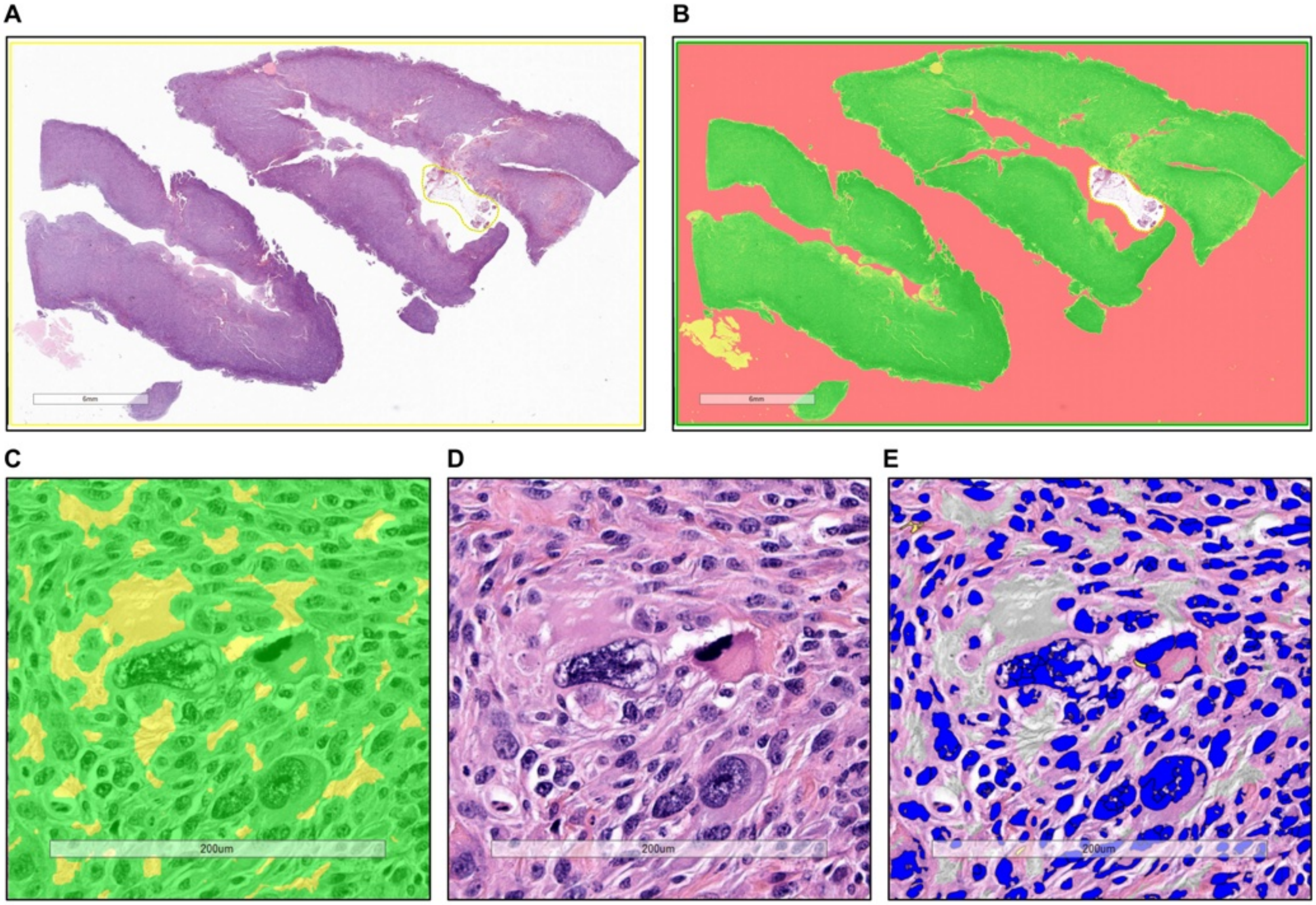


**Appendix Fig. 3. Validation of tissue composition metrics using H&E-stained serial sections. A-B.** Representative whole-slide images of H&E staining and the associated Genie classifier. The classifier segments tissue into nuclei (green), cytoplasm and ECM (yellow), and glass (pink) components. **C.** High-magnification view of the H&E Genie tissue classifier. **D.** H&E staining corresponding to the image shown in **C**. **E.** Nuclear v9 algorithm applied only to the "nuclei" class of the image shown in **C-D**. Note that the assignment of nuclei to $0^+$ (negative; blue), $1^+$ (yellow), $2^+$ (orange), and $3^+$ (red) staining categories was disregarded for this algorithm.

We then correlated the nuclear area metrics obtained from the H&E-stained slides with those from the IHC-stained specimens and found strong associations (**Appendix Fig. 4A-B**).

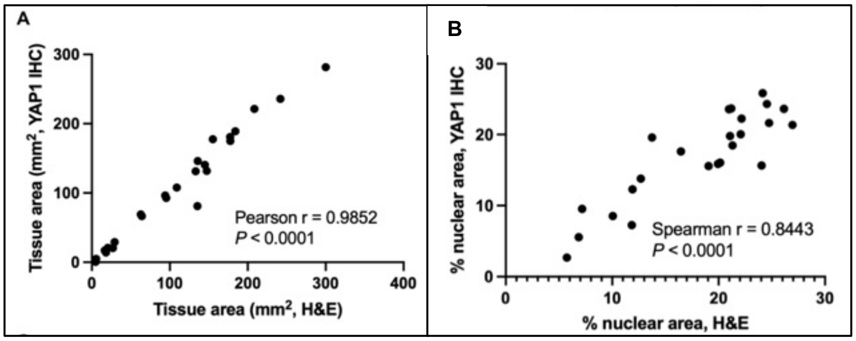


**Appendix Fig. 4. Correlation between tissue composition metrics from IHC- and H&E-stained slides. A.** Correlation between tissue area values obtained from the H&E- and YAP1-stained slides. **B.** Correlation between % nuclear area values obtained from the H&E- and YAP1-stained slides. For both panels, the Pearson correlation coefficient was used if both datasets in a given comparison were normally distributed. The Spearman correlation was used if at least one dataset was not normally distributed. The Shapiro-Wilk test was used to assess normality.

Finally, the YAP1 H-score calculated in analysis I was normalized to the nuclear percent area values obtained from the H&E-stained slides. These normalized values were then correlated with COLVI H-scores for all available specimens.

$$\text{H\&E-normalized Nuclear YAP1 H-score} = \frac{\text{Nuclear YAP1 H-score}}{\text{\% nuclear area}_{H\&E}}$$

Significant correlations between COLVI and nuclear YAP1 were observed when tissue composition metrics were obtained directly from the YAP1-stained slides (**Appendix Fig. 5A**; see also **Figure 7J** in the main manuscript). When tissue composition metrics were calculated from the corresponding H&E-stained sections, the correlation between COLVI and nuclear YAP1 was attenuated but remained statistically significant (**Appendix Fig. 5B**). Taken together, we conclude that COLVI expression in the tumor microenvironment of UPS specimens is significantly associated with YAP1 pathway activity.

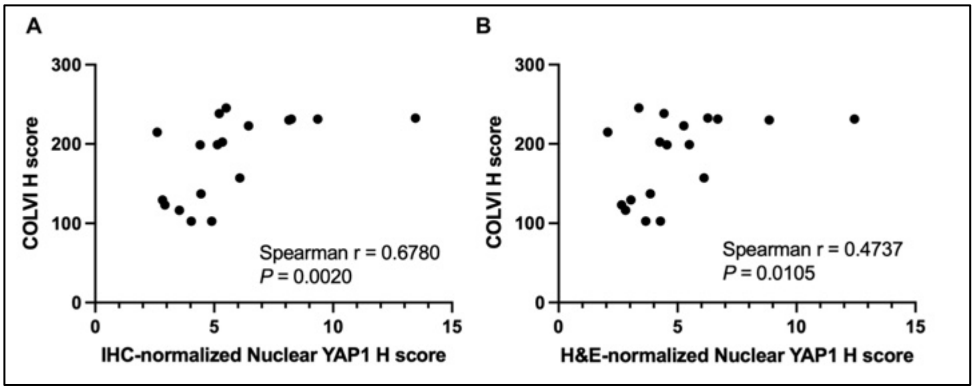


**Appendix Fig. 5. Comparison of nuclear YAP1-COLVI correlations using different normalization methods. A.** Correlation between COLVI H-score and nuclear YAP1 H-score normalized using tissue composition metrics obtained directly from the YAP1-stained slides. **B.** Correlation between COLVI H-score and nuclear YAP1 H-score normalized using tissue composition metrics obtained from the corresponding H&E-stained sections. For both panels, the Pearson correlation coefficient was used if both datasets in a given comparison were normally distributed. The Spearman correlation was used if at least one dataset was not normally distributed. The Shapiro-Wilk test was used to assess normality.

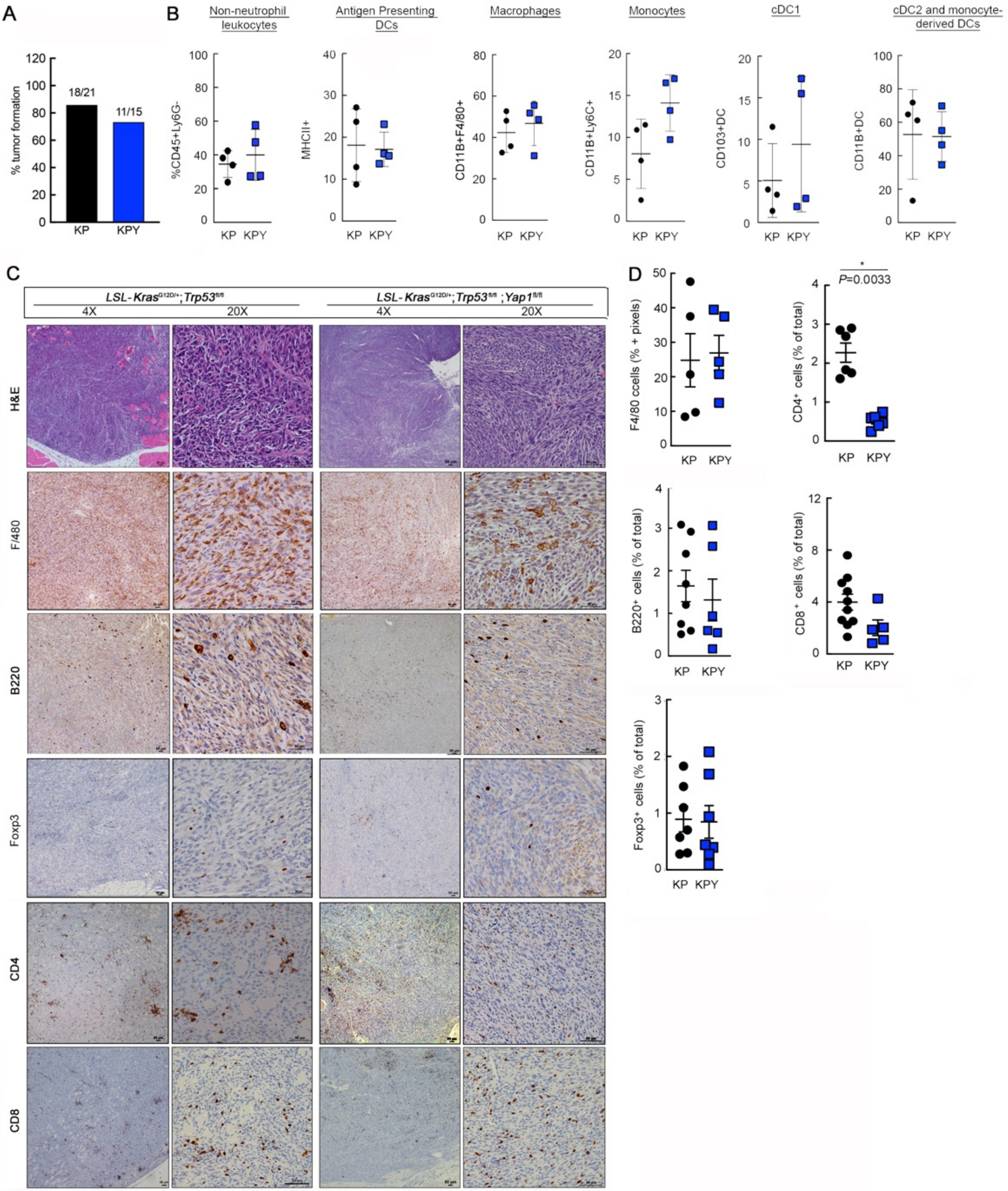
A
% tumor formation
18/21
11/15
KP
KPY
B
Non-neutrophil leukocytes
Antigen Presenting DCs
Macrophages
Monocytes
cDC1
cDC2 and monocyte-derived DCs
%CD45+Ly6G-
MHCII+
CD11B+F4/80+
CD11B+Ly6C+
CD103+DC
CD11B+DC
C
LSL-Kras G12D/+;Trp53 fl/fl
LSL-Kras G12D/+;Trp53 fl/fl ;Yap1 fl/fl
4X
20X
H&E
F/480
B220
Foxp3
CD4
CD8
D
F4/80 ccells (% + pixels)
CD4+ cells (% of total)
P=0.0033
B220+ cells (% of total)
CD8+ cells (% of total)
Foxp3+ cells (% of total)

**Supplementary Figure 1.** (**A**) Tumor formation rates of KP and KPY mice shown in **Fig. 1A.** (**B**) Frequency of non-neutrophil leukocytes, monocytes, macrophages, and DC cell subsets (antigen-presenting DCs, CD103$^+$ DCs, and CD11b$^+$ DCs) in KP and KPY tumors. Each point represents an individual tumor. Two-tailed unpaired t-tests. (**C**) Representative images of H&E and automated IHC staining of F4/80 (macrophages), B220 (B cells), Foxp3 (regulatory T cells), CD4$^+$ T cells, and CD8$^+$ T cells in sections from KP and KPY tumors. Scale bars = 50 $\mu$m. (**D**) Quantification of IHC staining from **C**. Each point represents an individual tumor. Two-tailed unpaired t-tests.

Supplementary Figure 2

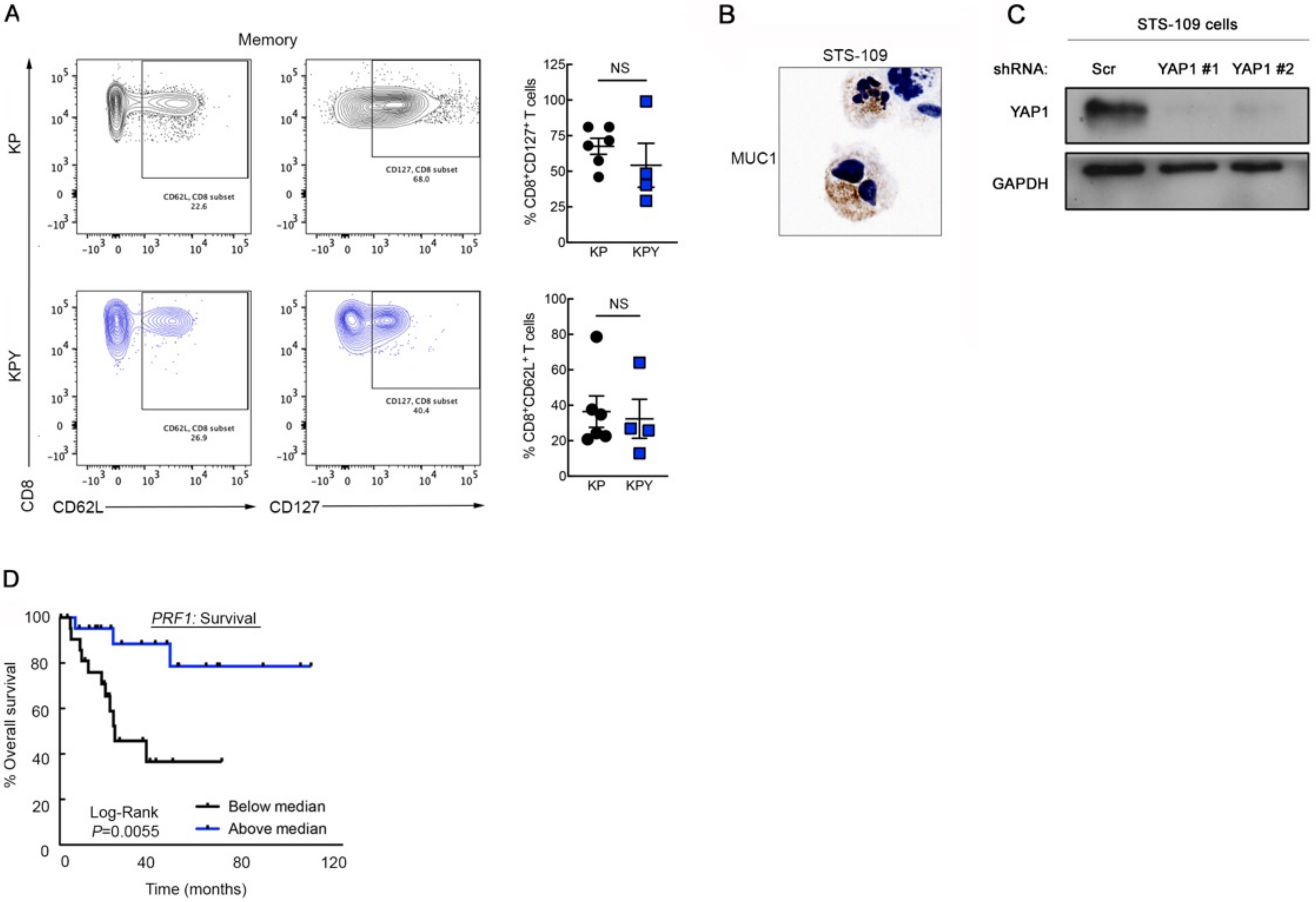

**Supplementary Figure 2.** (**A**) Representative flow cytometry plots and quantification of memory T cell markers (CD62L, CD127) in $CD8^+$ T cells from KP and KPY tumors. Each point in the quantification plots represents an individual mouse tumor. Two-tailed unpaired t-test. Top panel: $P$ = 0.3731; bottom panel: $P$ = 0.7822 (not significant). (**B**) Immunocytochemistry of Tn-MUC1 expression in STS-109 human UPS cells. (**C**) Representative immunoblot of YAP1 expression in STS-109 human UPS cells expressing a control or one of multiple *YAP1*-targeting shRNAs. (**D**) Overall survival of UPS patients in TCGA-Sarcoma (n = 44) stratified by tumor expression levels of *PRF1*. Log-rank test.

Supplementary Figure 3

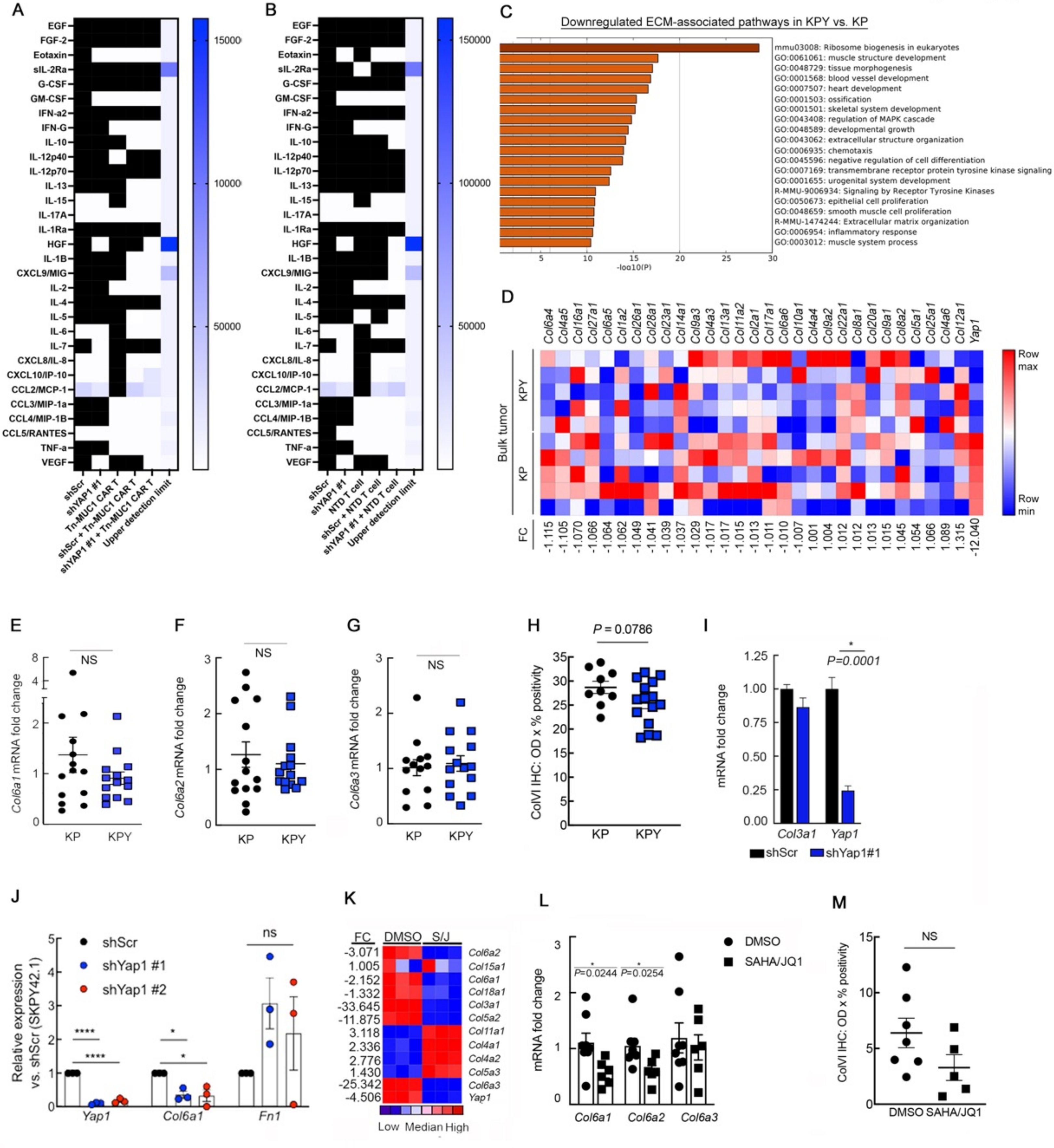

**Supplementary Figure 3.** (**A**) Expression levels of cytokines and chemokines in supernatants from human UPS cell-TnMUC1 CAR T cell co-cultures and the corresponding mono-cultures. Signal from matched co-cultures of UPS cells with un-transduced T cells (non-CAR T cells) is subtracted out. Black squares indicate undetected analytes. (**B)** Expression levels of cytokines and chemokines in supernatants from human UPS cell-normal human donor T (NTD) cell co-cultures and the corresponding mono-cultures. Black squares indicate undetected analytes. (**C**) Metascape pathway enrichment analysis of genes identified by microarray of 5 unique bulk KP and KPY tumors, highlighting extracellular matrix-associated pathways. Analysis includes all genes with >2-fold increase in expression in KP relative to KPY. (**D**) Heat map of gene expression microarray data comparing 5 unique KP and KPY bulk tumors. Map displays the bottom 2/3 of collagen-encoding genes modulated by *Yap1 in vivo* deletion. **Fig. 2A** shows the remaining top 1/3 of collagen-encoding genes. FC = fold change. (**E-G**) qRT-PCR analysis of *Col6a1* (**E**), *Col6a2* (**F**), and *Col6a3* (**G**) gene expression in bulk KP and KPY tumors. Two-tailed unpaired t-test. *P* = 0.4815, 0.8415, and 0.7052 for **D**, **E**, and **F**, respectively (not significant). Each point represents an individual tumor. (**H**) Additional quantification of KP and KPY bulk tumor ColVI IHC shown in **Fig. 2B, C**. Two-tailed unpaired t-test with Welch correction. (**I**) qRT-PCR analysis of *Col3a1* and *Yap1* gene expression in KP cells expressing control or *Yap1* shRNA #1. Two-tailed unpaired t-tests. (**J**) Fibronectin 1 (*Fn1*) gene expression in KP tumor-derived UPS cells expressing a control or one of multiple *Yap1*-targeting shRNAs. *Col6a1* expression in these cells is shown as a control. One-way ANOVA with Dunnett's (vs. shScr) for each gene; n =3. (**K**) Heat map of gene expression microarray data comparing KP cells treated with SAHA (2 μM) + JQ1 (0.5 μM) or vehicle control for 48 hours. (**L**) *Col6a1*, *Col6a2*, and *Col6a3* gene expression in bulk KP tumors from mice treated with 25 mg/kg SAHA + 50 mg/kg JQ1 or vehicle control for 20 days. Two-tailed unpaired t-tests, SEM. Each point represents an individual tumor. (**M**) Quantification of ColVI IHC in SAHA/JQ1-treated KP tumors from **Fig. 2F**. Each point represents an individual tumor.

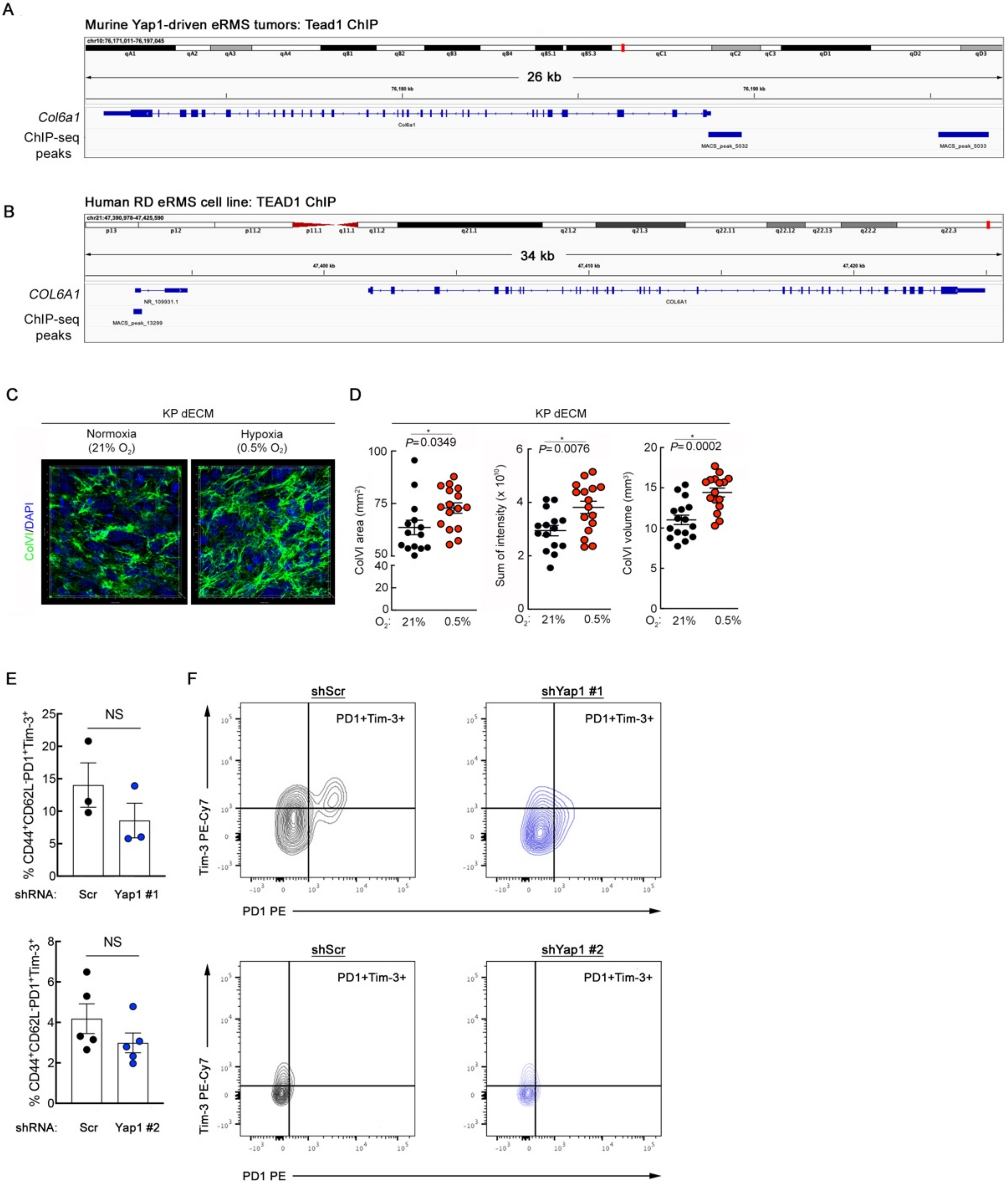
A
Murine Yap1-driven eRMS tumors: Tead1 ChIP
26 kb
Col6a1
ChIP-seq peaks
MACS_peak_5032
MACS_peak_5033
B
Human RD eRMS cell line: TEAD1 ChIP
34 kb
COL6A1
ChIP-seq peaks
NR_109931.1
MACS_peak_13299
C
KP dECM
Normoxia (21% O2)
Hypoxia (0.5% O2)
ColVI/DAPI
D
KP dECM
P=0.0349
P=0.0076
P=0.0002
ColVI area (mm2)
Sum of intensity (x 10^10)
ColVI volume (mm3)
O2: 21% 0.5%
E
NS
% CD44+CD62L-PD1+Tim-3+
shRNA: Scr Yap1 #1
shRNA: Scr Yap1 #2
F
shScr
shYap1 #1
shYap1 #2
PD1+Tim-3+
Tim-3 PE-Cy7
PD1 PE

**Supplementary Figure 4.** (**A**) Tead1-ChIP signal and MACS-based peak calling at the *Col6a1* locus in murine Yap1-driven embryonal rhabdomyosarcoma (eRMS) tumors. (**B**) TEAD1-ChIP signal and MACS-based peak calling at the *COL6A1* locus in cultured human eRMS cells (RD cell line). Data from **A, B** are from the publicly available dataset GSE55186. (**C, D**) Representative confocal micrographs of ColVI immunofluorescent staining (**C**) and quantification (**D**) in KP cell-derived dECMs generated under normoxic or hypoxic conditions. (**E, F**) Quantification (**E**) and representative contour plots (**F**) of Pd1 and Tim-3 co-expression in murine $CD44^{+}CD8^{+}$ T cells incubated on dECMs from control and shYap1-expressing B6-KP cells. Yap1 #1 and Yap1 #2 are shown separately with their respective controls because the assays were performed at different times, at different institutions, and on different instruments. Each point represents T cells isolated from an individual mouse. Two-tailed unpaired t-tests.

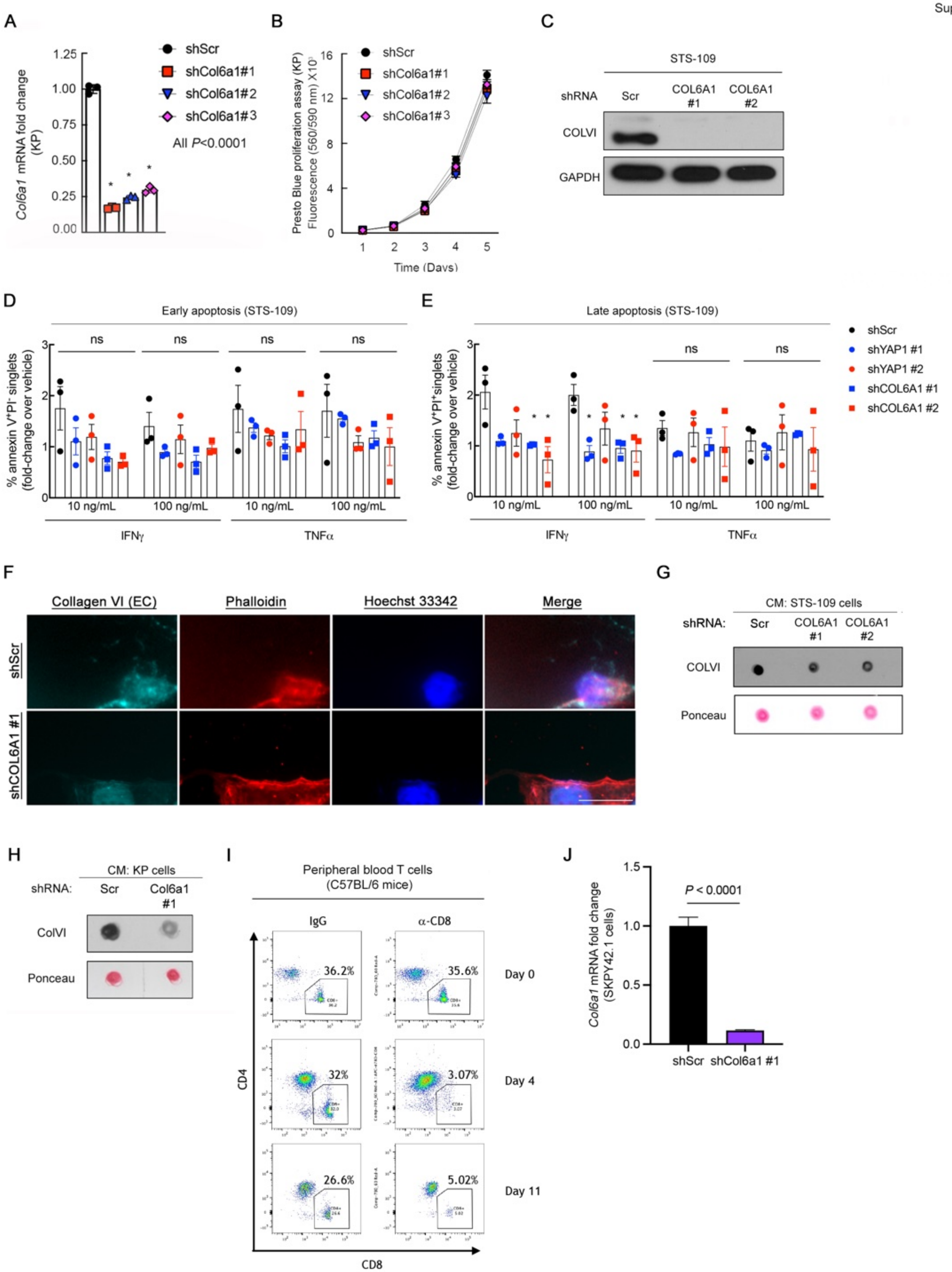
A
shScr
shCol6a1#1
shCol6a1#2
shCol6a1#3
All P<0.0001
Col6a1 mRNA fold change (KP)
B
Presto Blue proliferation assay (KP) Fluorescence (560/590 nm) X10
Time (Days)
C
STS-109
shRNA
Scr
COL6A1 #1
COL6A1 #2
COLVI
GAPDH
D
Early apoptosis (STS-109)
% annexin V+PI- singlets (fold-change over vehicle)
ns
10 ng/mL
100 ng/mL
IFNγ
TNFα
E
Late apoptosis (STS-109)
% annexin V+PI+singlets (fold-change over vehicle)
shScr
shYAP1 #1
shYAP1 #2
shCOL6A1 #1
shCOL6A1 #2
F
Collagen VI (EC)
Phalloidin
Hoechst 33342
Merge
shScr
shCOL6A1 #1
G
CM: STS-109 cells
shRNA:
COLVI
Ponceau
H
CM: KP cells
Col6a1 #1
ColVI
I
Peripheral blood T cells (C57BL/6 mice)
IgG
α-CD8
Day 0
Day 4
Day 11
CD4
CD8
36.2%
35.6%
32%
3.07%
26.6%
5.02%
J
Col6a1 mRNA fold change (SKPY42.1 cells)
P < 0.0001
shScr
shCol6a1 #1

**Supplementary Figure 5.** (**A**) *Col6a1* gene expression levels in cultured KP cells expressing a control or one of multiple *Col6a1*-targeting shRNAs. One-way ANOVA with Dunnett's. (**B**) Presto Blue proliferation assay of KP cells cultured as in (**A**). (**C**) Representative immunoblot of COLVI expression in STS-109 human UPS cells expressing a control or one of multiple *COL6A1*-targeting shRNAs. (**D, E**) Flow cytometric apoptosis assay showing the proportion of early (**D**) and late (**E**) apoptotic STS-109 cells expressing control, *YAP1*, or *COL6A1*-targeting shRNAs after treatment with recombinant human IFN$\gamma$ or TNF$\alpha$. Normalization to vehicle control was done to account for potential differences in baseline apoptosis levels. One-way ANOVA with Dunnett's (vs. shScr) for each condition. N = 3. (**F**) Widefield images confirming extracellular (EC) COLVI expression in STS-109 human UPS cells cultured on cytolysis arrays (as in **Fig. 3D, E**). Phalloidin marks cell boundaries. COLVI staining was performed prior to cell permeabilization in order to primarily detect extracellular COLVI. Scale bar = 15 $\mu$m. (**G**, **H**) Representative immuno-dot blot of ColVI in conditioned media (CM) collected from sub-confluent STS-109 (**G**) and KP (**H**) UPS cells expressing control or *Col6a1*-targeting shRNAs. Ponceau stain shows equal protein loading. Each dot experiment was performed at least 2-3 times. (**I**) Representative flow cytometry plots of $CD4^+$ and $CD8^+$ cells from peripheral blood of C57BL/6 mice following depletion of $CD8^+$ T cells with $\alpha$-CD8$\alpha$. (**J**) qRT-PCR validation of *Col6a1* expression in KP cells derived from C57BL/6 mice (SKPY42.1 cell line) immediately prior to syngeneic orthotopic transplant into the gastrocnemius muscles of C57BL/6 mice (corresponds to **Fig. 3I**).

Supplementary Figure 6

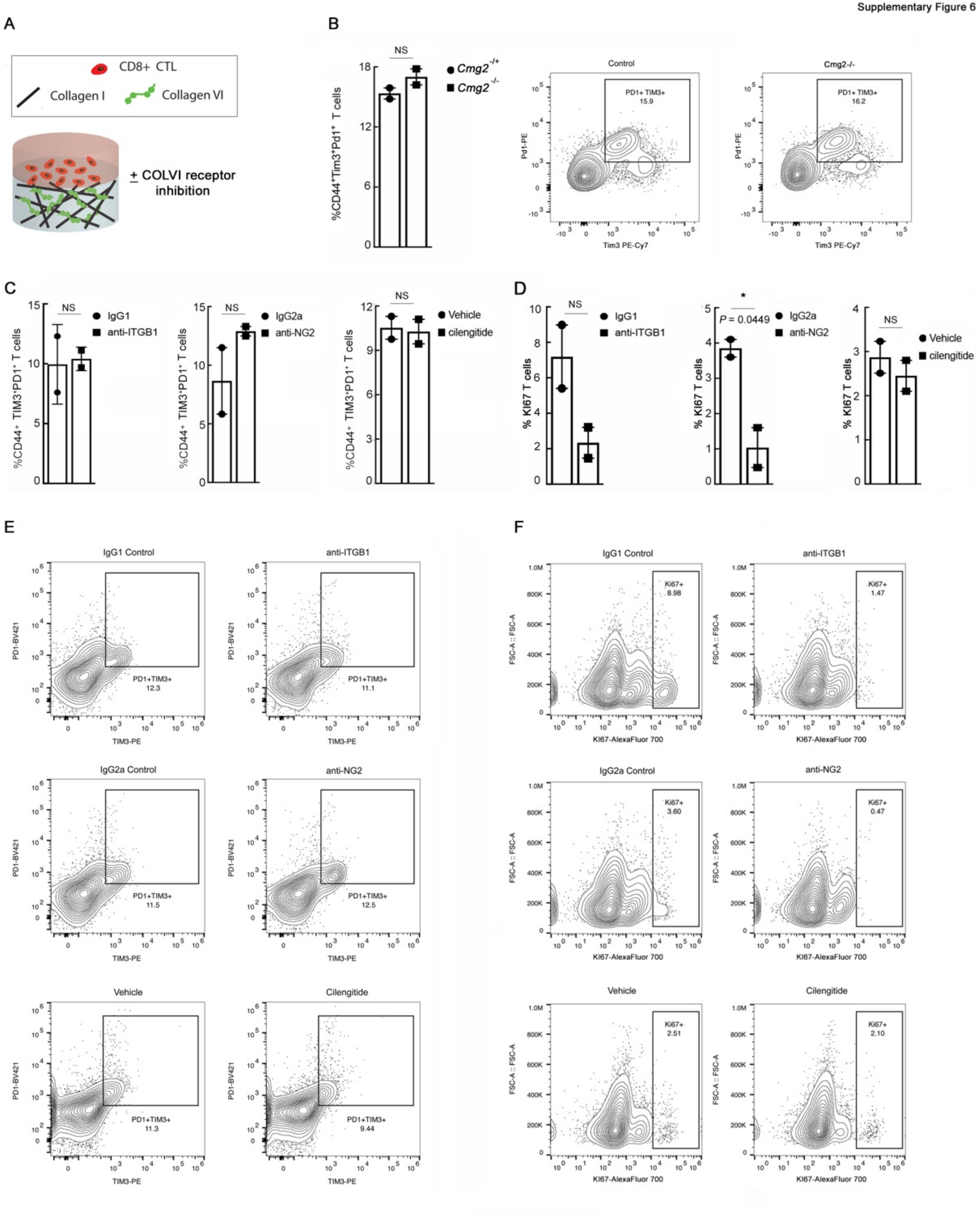

**Supplementary Figure 6.** (**A**) Schematic of hydrogel system to identify COLVI receptors on CD8$^+$ T cells controlling ECM-mediated dysfunction. (**B, C**) Quantification (**B**) and representative flow cytometry plots (**C**) of Tim-3 and Pd1 co-expression in activated *Cmg2* null or control murine splenocytes cultured on hydrogels containing purified COLVI. Each point represents T cells from an individual mouse. Two-tailed unpaired t-test. *P* = 0.2313 (not significant). (**C**) Quantification of TIM-3 and PD-1 co-expression in activated human CD8$^+$CD44$^+$ T cells cultured on purified COLVI-containing hydrogels and treated with ITGB1-blocking antibody, NG2-blocking antibody, isotype control antibody, Cilengitide, or vehicle control. Each point represents an independent experiment. Cells in each replicate were from independent healthy donors. Two-tailed unpaired t-tests. *P* = 0.8698 (ITGB1), *P* = 0.2772 (NG2), *P* = 0.8397 (cilengitide) (not-significant). (**D**) Quantification of KI67 expression in activated human CD8$^+$CD44$^+$ T cells cultured and treated as in **C**. Each point represents an independent experiment. Cells in each replicate were from independent healthy donors. Two-tailed unpaired t-test. *P* = 0.1351 (ITGB1), *P* = 0.4909 (cilengitide) (not-significant). (**E**) Representative flow cytometry plots of data from **C**. (**F**) Representative flow cytometry plots of data from **D**.

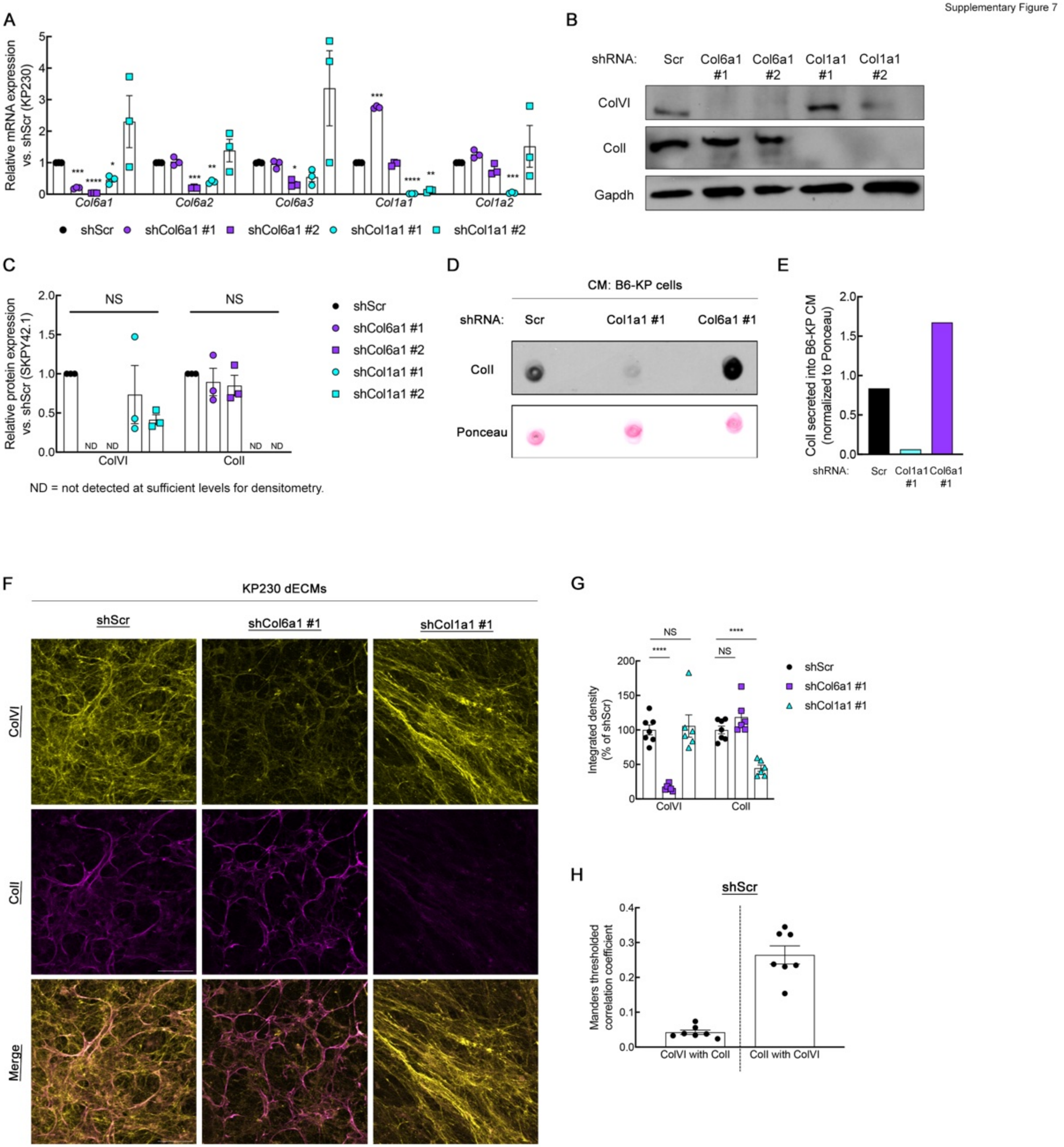
A
Relative mRNA expression vs. shScr (KP230)
Col6a1
Col6a2
Col6a3
Col1a1
Col1a2
shScr
shCol6a1 #1
shCol6a1 #2
shCol1a1 #1
shCol1a1 #2
B
shRNA:
Scr
Col6a1 #1
Col6a1 #2
Col1a1 #1
Col1a1 #2
ColVI
ColI
Gapdh
C
Relative protein expression vs. shScr (SKPY42.1)
NS
NS
ND
ColVI
ColI
ND = not detected at sufficient levels for densitometry.
D
CM: B6-KP cells
shRNA:
Scr
Col1a1 #1
Col6a1 #1
ColI
Ponceau
E
ColI secreted into B6-KP CM (normalized to Ponceau)
shRNA:
Scr
Col1a1 #1
Col6a1 #1
F
KP230 dECMs
shScr
shCol6a1 #1
shCol1a1 #1
ColVI
ColI
Merge
G
Integrated density (% of shScr)
NS
****
NS
****
ColVI
ColI
shScr
shCol6a1 #1
shCol1a1 #1
H
shScr
Manders thresholded correlation coefficient
ColVI with ColI
ColI with ColVI

**Supplementary Figure 7.** (**A**) qRT-PCR analysis of *Col6a1* and *Col1a1* gene expression in KP tumor-derived UPS cells expressing a control or one of multiple *Col6a1-* or *Col1a1*-targeting shRNAs. One-way ANOVA with Dunnett's for each gene (vs. shScr). N = 3. (**B, C**) Representative immunoblot (**B**) and (**C**) quantification of ColVI and ColI protein expression in KP tumor-derived UPS cells expressing a control or one of multiple *Col6a1-* or *Col1a1*-targeting shRNAs. One-way ANOVA with Dunnett's for each protein (vs. shScr). ND = not detected at sufficient levels for robust densitometry analysis. N = 3. (**D, E**) Representative immuno-dot blot (**D**) and quantification (**E**) of ColI in conditioned media (CM) collected from sub-confluent KP UPS cells expressing control, *Col1a1-*, or *Col6a1*-targeting shRNAs. Ponceau stain shows equal protein loading. Quantification is representative of n $\geq$2 experiments. (**F, G**) Representative confocal micrographs (maximum-intensity Z-projections) (**F**) and quantification (**G**) showing ColI and ColVI co-immunofluorescence in dECMs from KP cells. Scale bars = 50 $\mu$m. Quantification of each protein was performed in 7 representative fields across 3 independent dECMs per condition. One-way ANOVA with Dunnett's (vs. shScr) for each gene. (**H**) Mander's correlation coefficients depicting ColI and ColVI colocalization in images from **F, G**. Colocalization was determined for each of 7 representative fields across 3 independent dECMs per condition.

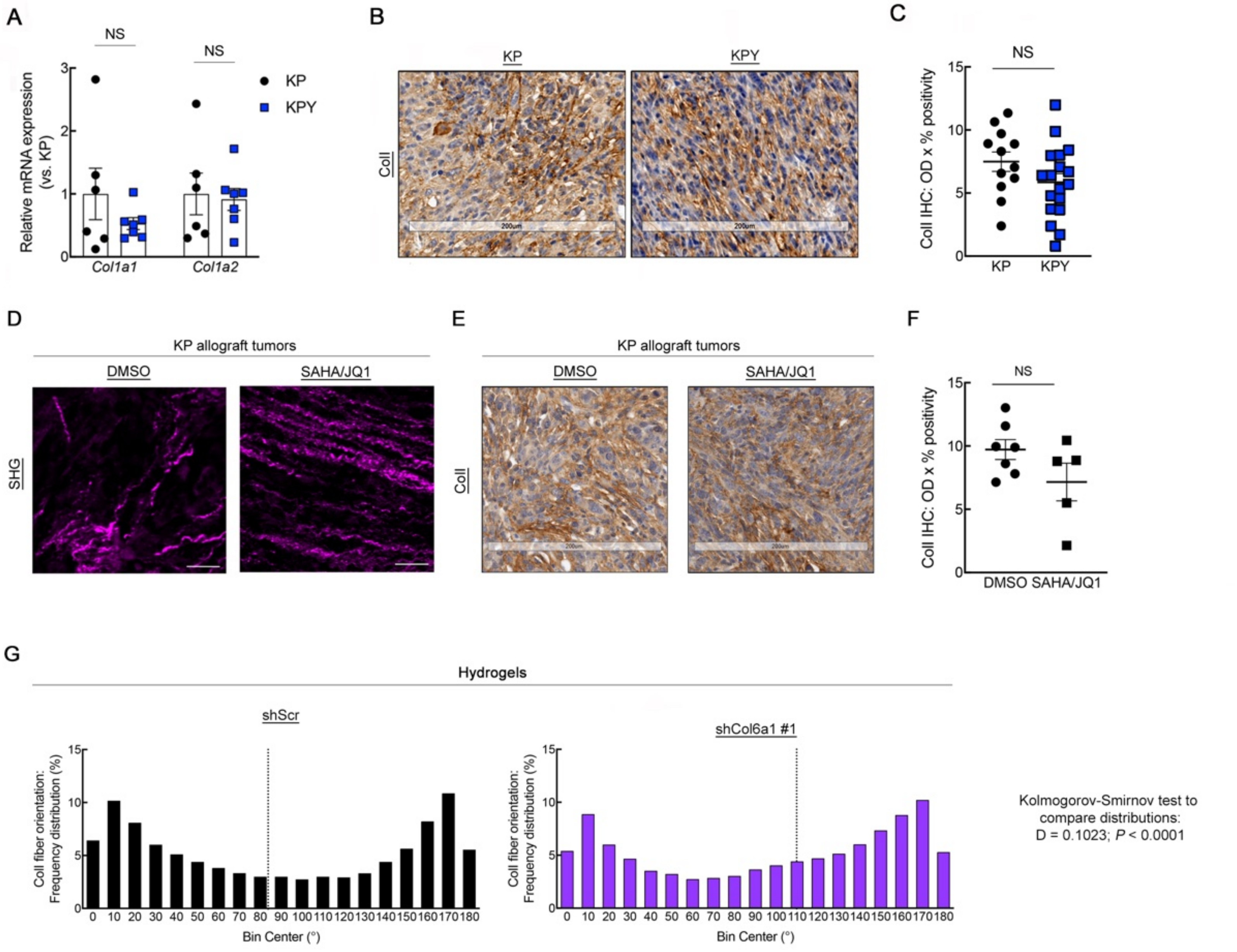
A
NS
NS
KP
KPY
Relative mRNA expression (vs. KP)
Col1a1
Col1a2
B
KP
KPY
ColI
C
NS
ColI IHC: OD x % positivity
KP
KPY
D
KP allograft tumors
DMSO
SAHA/JQ1
SHG
E
KP allograft tumors
DMSO
SAHA/JQ1
ColI
F
NS
ColI IHC: OD x % positivity
DMSO
SAHA/JQ1
G
Hydrogels
shScr
shCol6a1 #1
ColI fiber orientation: Frequency distribution (%)
Bin Center (°)
Kolmogorov-Smirnov test to compare distributions:
D = 0.1023; P < 0.0001

**Supplementary Figure 8.** (**A**) qRT-PCR of *Col1a1* and *Col1a2* gene expression in bulk KP and KPY tumors. Each point represents an individual tumor. Two-tailed unpaired t-tests. (**B**) Representative ColI IHC staining in KP and KPY tumors. (**C**) Quantification of KP and KPY tumor ColI IHC shown in **B**. N = 3-5 mice per genotype with 3 sections per mouse. Two-tailed unpaired t-test with Welch correction. (**D**) Representative multiphoton second-harmonic generation (SHG) images (maximum-intensity Z-projections) of sections from KP tumors treated with 25 mg/kg SAHA + 50 mg/kg JQ1 or vehicle control for 20 days. All scale bars = 50 $\mu$M. (**E, F**) Representative images (**E**) and quantification (**F**) of ColI IHC staining in tumors from mice treated as in D. Not significant by two-tailed, unpaired t-test. (**G**) Frequency distribution histograms of ColI fiber orientation in hydrogel images from **Fig. 4K**. Dotted vertical line in each histogram indicates the median fiber orientation angle.

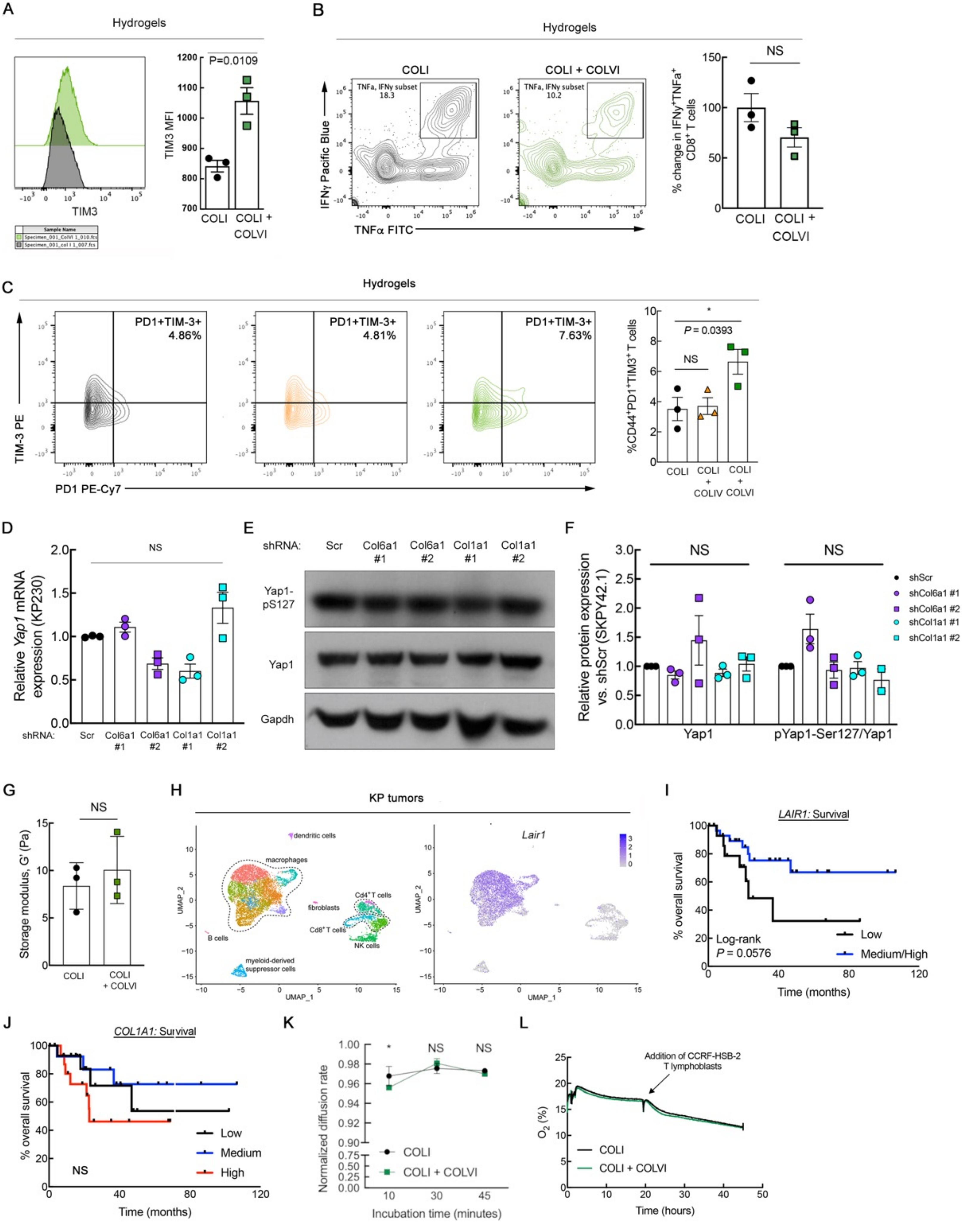
A
Hydrogels
TIM3
TIM3 MFI
P=0.0109
COLI
COLI + COLVI
B
Hydrogels
COLI
COLI + COLVI
TNFa, IFNy subset 18.3
TNFa, IFNy subset 10.2
IFNγ Pacific Blue
TNFα FITC
% change in IFNγ+TNFa+ CD8+ T cells
NS
C
Hydrogels
PD1+TIM-3+ 4.86%
PD1+TIM-3+ 4.81%
PD1+TIM-3+ 7.63%
TIM-3 PE
PD1 PE-Cy7
%CD44+PD1+TIM3+ T cells
P = 0.0393
NS
COLI
COLI + COLIV
COLI + COLVI
D
Relative Yap1 mRNA expression (KP230)
NS
shRNA: Scr Col6a1 #1 Col6a1 #2 Col1a1 #1 Col1a1 #2
E
shRNA: Scr Col6a1 #1 Col6a1 #2 Col1a1 #1 Col1a1 #2
Yap1-pS127
Yap1
Gapdh
F
Relative protein expression vs. shScr (SKPY42.1)
NS
Yap1
pYap1-Ser127/Yap1
shScr
shCol6a1 #1
shCol6a1 #2
shCol1a1 #1
shCol1a1 #2
G
Storage modulus, G' (Pa)
NS
COLI
COLI + COLVI
H
KP tumors
dendritic cells
macrophages
fibroblasts
Cd4+ T cells
Cd8+ T cells
NK cells
B cells
myeloid-derived suppressor cells
UMAP_1
UMAP_2
Lair1
I
LAIR1: Survival
% overall survival
Log-rank P = 0.0576
Low
Medium/High
Time (months)
J
COL1A1: Survival
% overall survival
NS
Low
Medium
High
Time (months)
K
Normalized diffusion rate
NS
COLI
COLI + COLVI
Incubation time (minutes)
L
Addition of CCRF-HSB-2 T lymphoblasts
O2 (%)
COLI
COLI + COLVI
Time (hours)

**Supplementary Fig. 9.** (**A**) TIM-3 median fluorescence intensity (MFI) in activated human CD8$^+$ T cells incubated on hydrogels containing purified COLI, with or without purified COLVI. Two-tailed unpaired t-test. (**B**) Representative contour plots and quantification of IFN$\gamma$ and TNF$\alpha$ co-expression in activated human CD8$^+$ T cells incubated on hydrogels containing purified COLI, with or without purified COLVI. Two-tailed unpaired t-test. (**C**) Representative contour plots and quantification of PD1 and TIM-3 co-expression in activated human CD8$^+$ T cells incubated on hydrogels containing purified COLI, with or without purified COLIV or purified COLVI. One-way ANOVA with Dunnett's (vs. COLI-alone). Each point represents CD8$^+$ T cells procured from an individual healthy human donor. (**D**) *Yap1* gene expression levels in KP-derived UPS cells (KP230 cell line) expressing a control or one of multiple *Col6a1*- or *Col1a1*- targeting shRNAs. One-way ANOVA with Dunnett's. N = 3. (**E, F**) Representative immunoblot (**E**) and quantification (**F**) of Yap1 protein expression and S127 phosphorylation in SKPY42.1 cells treated as in **D**. One-way ANOVA with Dunnett's for each protein. N = 3. (**G**) Rheometry analysis of COLI hydrogel stiffness, with or without the addition of COLVI. Two-tailed student's t-test. (**H**) Single-cell RNA-seq data of CD45$^+$ leukocyte populations from KP tumors showing cell type distributions of *Lair1* gene expression. *Lair1* is primarily expressed on tumor-associated macrophages and only minimally on CD8$^+$ T cells. Data are from the publicly available dataset GSE144507. (**I, J**) Kaplan Meier survival curve of UPS patients in TCGA-Sarcoma dataset (n = 44) based on *LAIR1* (**I**) and *COL1A1* (**J**) gene expression. Each tertile (low, medium, high), represents 1/3 of patients. Log-rank test. (**K**) Rhodamine B assay showing normalized diffusion rates throughout COLI hydrogels over time, with or without the addition of purified COLVI. Two-way ANOVA with Sidak's. (**L**) Trace curves showing non-invasive oxygen readings of the COLI hydrogel system, with or without the addition of purified COLVI, before and after the addition of a T lymphoblast cell line.

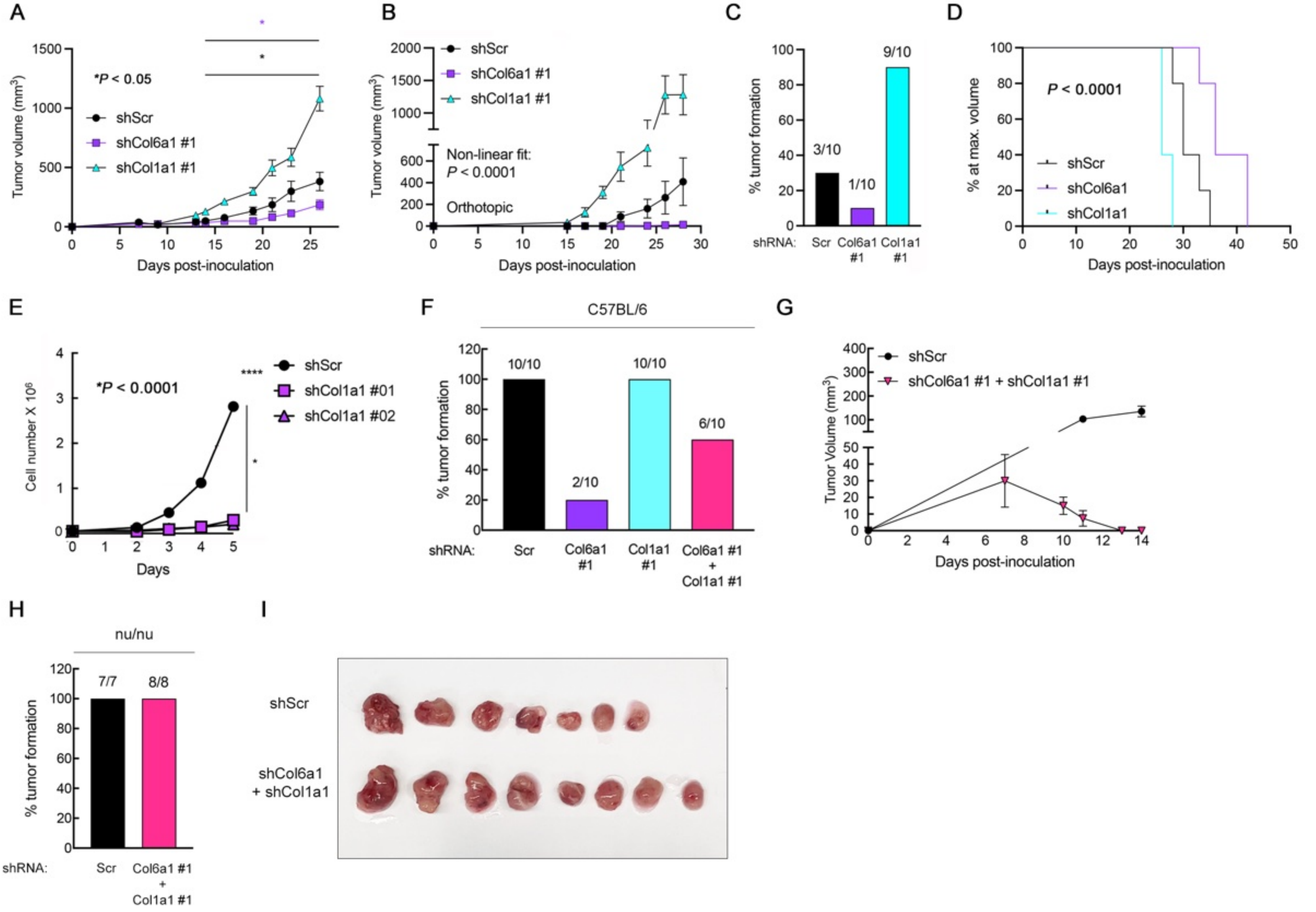
A
*P < 0.05
shScr
shCol6a1 #1
shCol1a1 #1
Tumor volume (mm3)
Days post-inoculation
B
shScr
shCol6a1 #1
shCol1a1 #1
Non-linear fit:
P < 0.0001
Orthotopic
Tumor volume (mm3)
Days post-inoculation
C
3/10
1/10
9/10
% tumor formation
shRNA: Scr Col6a1 #1 Col1a1 #1
D
P < 0.0001
shScr
shCol6a1
shCol1a1
% at max. volume
Days post-inoculation
E
*P < 0.0001
shScr
shCol1a1 #01
shCol1a1 #02
Cell number X 10^6
Days
F
C57BL/6
10/10
2/10
10/10
6/10
% tumor formation
shRNA: Scr Col6a1 #1 Col1a1 #1 Col6a1 #1 + Col1a1 #1
G
shScr
shCol6a1 #1 + shCol1a1 #1
Tumor Volume (mm3)
Days post-inoculation
H
nu/nu
7/7
8/8
% tumor formation
shRNA: Scr Col6a1 #1 + Col1a1 #1
I
shScr
shCol6a1 + shCol1a1

**Supplementary Fig. 10.** (**A**) Tumor growth curves from orthotopic (gastrocnemius muscle) syngeneic transplant of 2.5 x $10^5$ KP cells (SKPY42.1 cell line) expressing control, *Col6a1*, or *Col1a1*-targeting shRNAs in C57BL/6 mice. Two-way repeated-measures ANOVA with Dunnett's (vs. shScr). SEM. (**B**) Tumor growth curves from orthotopic (gastrocnemius muscle) syngeneic transplant of 5 x $10^5$ KP cells (SKPY42.1 cell line) expressing control, *Col6a1*, or *Col1a1*-targeting shRNAs in C57BL/6 mice. Non-linear regression (exponential fit) to compare slopes. (**C**) Tumor formation rates from **B**. (**D**) Kaplan-Meier survival curves of C57BL/6 mice bearing control, shCol6a1, or shCol1a1 syngeneic KP tumors (SKPY42.1 cell line). Log-rank test. (**E**) *In vitro* proliferation assay of KP cells expressing a control or one of multiple *Col1a1*-targeting shRNAs. Two-way ANOVA, SEM. (**F**) Tumor take rates from subcutaneous syngeneic transplant of 1 x $10^6$ KP cells (SKPY42.1 cell line) expressing control, *Col6a1*, and/or *Col1a1*-targeting shRNAs in C57BL/6 mice. (**G**) Tumor volume data from **Fig. 5N** zoomed in to show regression in dual shCol6a1/shCol1a1 tumors. (**H**) Tumor take rates from subcutaneous transplant of 1 x $10^6$ KP cells (SKPY42.1 cell line) expressing control or *Col6a1 and Col1a1*-targeting shRNAs in nu/nu mice. (**I**) Photograph of excised SKPY42.1 tumors from nu/nu mice (corresponds to data in **Fig. 5O** and **Supp. Fig. 10H**).

A

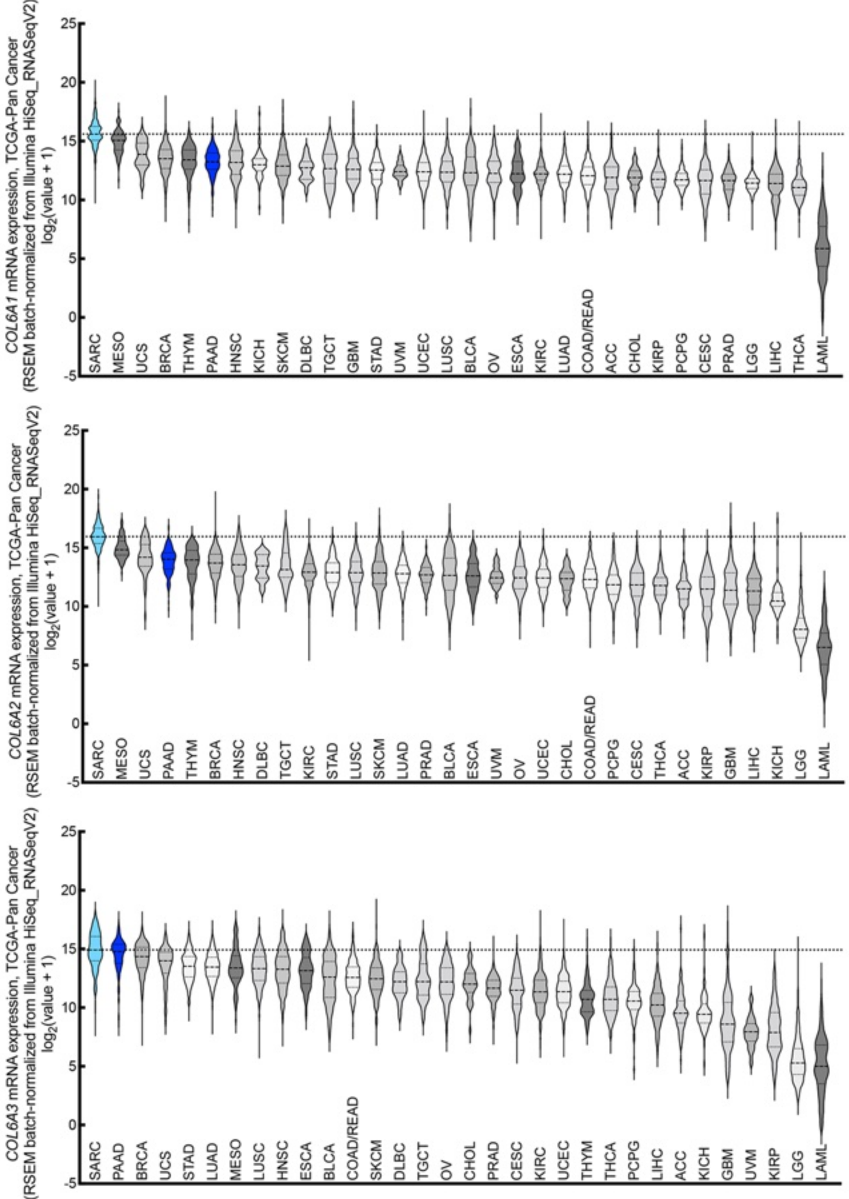


| TCGA study code | Cancer type |
|---|---|
| ACC | Adrenocortical carcinoma |
| BLCA | Bladder urothelial carcinoma |
| BRCA | Breast invasive carcinoma |
| CESC | Cervical squamous cell & endocervical adenocarcinoma |
| CHOL | Cholangiocarcinoma |
| COAD | Colon adenocarcinoma |
| DLBC | Diffuse large B cell lymphoma |
| ESCA | Esophageal carcinoma |
| GBM | Glioblastoma multiforme |
| HNSC | Head and neck squamous cell carcinoma |
| KICH | Kidney chromophobe |
| KIRC | Kidney renal clear cell carcinoma |
| KIRP | Kidney renal papillary cell carcinoma |
| LAML | Acute myeloid leukemia |
| LGG | Brain low-grade glioma |
| LIHC | Liver hepatocellular carcinoma |
| LUAD | Lung adenocarcinoma |
| LUSC | Lung squamous cell carcinoma |
| MESO | Mesothelioma |
| OV | Ovarian serous cystadenocarcinoma |
| PAAD | Pancreatic adenocarcinoma |
| PCPG | Pheochromocytoma and paraganglioma |
| PRAD | Prostate adenocarcinoma |
| SARC | Sarcoma |
| SKCM | Skin cutaneous melanoma |
| STAD | Stomach adenocarcinoma |
| READ | Rectum adenocarcinoma |
| TGCT | Testicular germ cell tumor |
| THCA | Thyroid carcinoma |
| THYM | Thymoma |
| UCEC | Uterine corpus endometrial carcinoma |
| UCS | Uterine carcinosarcoma |
| UVM | Uveal melanoma |

B

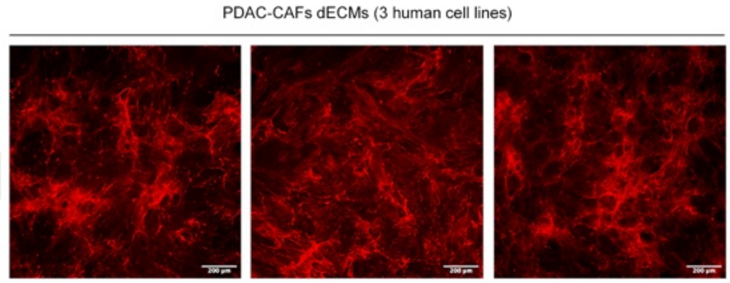


C

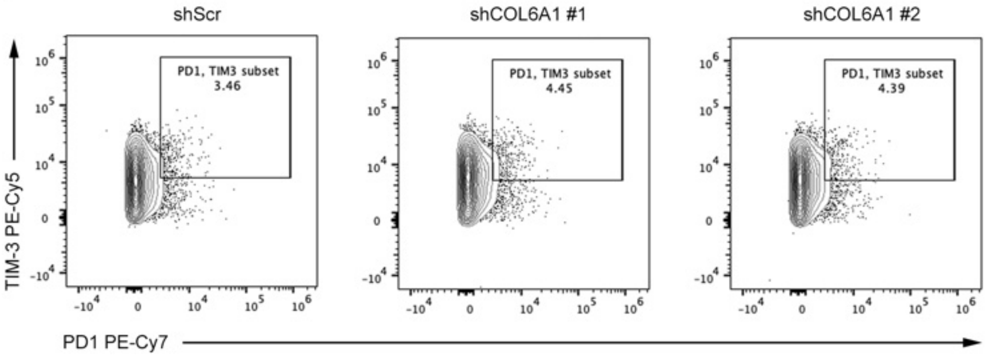


D

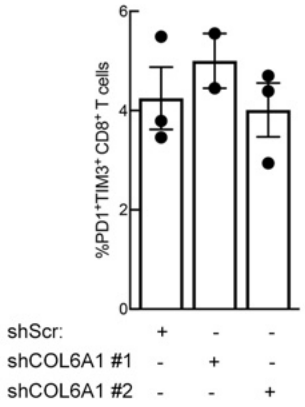


E

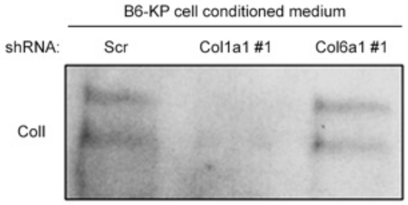

**Supplementary Fig. 11.** (**A**) TCGA PanCancer data (cBioPortal) showing *COL6A1*, *COL6A2*, and *COL6A3* expression levels in 32 human tumor types. Abbreviations are defined in the table. Sarcoma and pancreatic ductal adenocarcinoma (PDAC) tumors are shown in light blue and dark blue, respectively. (**B**) Representative micrographs showing COLVI immunofluorescence in dECMs generated from human PDAC-cancer-associated fibroblasts (CAFs), confirming their ability to secrete COLVI. Scale bars = 200 $\mu$m. (**C, D**) Representative contour plots (**C**) and quantification (**D**) of PD1 and TIM-3 co-expression in activated human $CD8^+$ T cells incubated on control or shCOL6A1 dECMs from PDAC CAFs. One-way ANOVA. (**E**) Immunoblot of COLI secreted into culture media by KP-derived UPS cells expressing a control or *Col6a1*-targeting shRNA. Two bands denote the presence of Col1a1 and Col1a2. shCol1a1-expressing cells are shown to demonstrate antibody specificity.

**Supplementary Movies S1-S3.** 3D rotating view of decellularized extracellular matrix (dECM) from control (shScr) KP cells (**Mov. S1)** co-stained for COLI (cyan) and COLVI (red). dECMs from shCol6a1 (**Mov. S2**) and shCol1a1 cells (**Mov. S3**) are also shown. A representative z-stack image of each dECM was surface-rendered into a movie rotating about the y-axis using Leica Las X 3D viewer software.

**Supplementary Movies S4-S6.** 3D rotating view of depth-coded multiphoton second-harmonic generation images from explanted KP (**Mov. S4**), KPY (**Mov. S5**), and human UPS (**Mov. S6**) tumors. Red = SHG signal farthest away from the objective/greatest relative depth in the tissue, blue= SHG signal closest to the objective/shallowest relative depth in the tissue.
